# An image-based transcriptomics atlas reveals the regional and microbiota-dependent molecular, cellular, and spatial structure of the murine gut

**DOI:** 10.1101/2025.07.21.665958

**Authors:** Rosalind J. Xu, Paolo Cadinu, Phillip B. Nicol, Uli S. Herrmann, Tyrone Lee, Ludwig Geistlinger, Rafael A. Irizarry, Jeffrey R. Moffitt

## Abstract

The gastrointestinal environment is home to a massive diversity of diet-, host-, and microbiota-derived small molecules, collectively sensed by a remarkable variety of cells. To explore the cellular and spatial organization of sensation, we used MERFISH to profile receptor expression across 2.1 million cells in multiple regions of the murine gut under specific-pathogen-free (SPF) and germ-free (GF) conditions. This atlas revealed expected and novel cell types—including a candidate murine homolog of human BEST4⁺ enterocytes—demonstrated cell-type regional specialization, discovered extensive location-dependent spatial fine-tuning in mucosal cell expression, and suggested cell-type specific mediators of the effects of microbiota-derived small molecules. In addition, this atlas revealed that, aside from immune cell abundance, many aspects of the murine gut are host-intrinsic and modified only modestly in the absence of a microbiota. Collectively, this atlas provides a valuable resource for understanding the cellular and spatial organization underlying small molecule sensation in the gut.

## INTRODUCTION

The gastrointestinal (GI) tract plays a complex role as one of the largest organs in the body. In addition to facilitating digestion, nutrient absorption, and immune response, it is home to a complex microbial community that contributes to a diverse small molecule environment through its metabolic activity^1,2^. A wide array of cell types, spanning multiple cell classes including epithelial, immune, fibroblast, endothelial, smooth muscle cells, as well as cells of the enteric nervous system, are responsible for executing these functions, and their behavior and interactions are shaped by the intricate spatial structure of the gut, including not just regional variations between the small and large intestine but local variations within the varied anatomical structures of the gut^3–5^.

Of its many functions, one central role of the gut is to sense and detect these diverse small molecules as well as other microbial- and host-derived signals^6^. These functions are mediated, in part, through the expression of a vast array of small molecule and signal receptors, including G-protein-coupled receptors (GPCRs), nuclear receptors, ligand-gated ion channels, pattern recognition receptors, complement receptors, enzyme-linked receptors, and cytokine receptors. The differential cell-type expression of these receptors, in turn, shapes how the gut responds to these molecular cues. Therefore, an understanding of gut sensation will require a mapping of the cell type of expression of these receptors.

To this end, single-cell RNA sequencing (scRNA-seq) has revolutionized our understanding of small molecule sensation in the gut by enabling detailed cataloging of both the cell type diversity and their gene expression profiles^3,5,7–13^. These efforts have led to important findings, including the identification of previously unrecognized epithelial^7–9^, immune^11^, fibroblast^12^, and enteric nervous system cell subsets^10^ and the discovery of region-specific transcriptional differences between the small and large intestine^14^. Nonetheless, several limitations have hindered a comprehensive understanding of sensory capabilities in the gastrointestinal tract. Receptor expression and cell type composition vary across gut regions, yet many existing atlases focus on single regions, leaving regional specialization—i.e. small or large intestine-specific gene expression within cell types—incompletely characterized. Additionally, limited cell numbers have reduced the resolution of rare but potentially important cell types and subtypes. These gaps highlight the need for a large-scale, pan-regional, single-cell atlas to comprehensively map gut cell diversity, sensory capabilities, and regional specialization.

Beyond regional specialization, multiple gut cell populations are known to fine-tune their gene expression over micron-scale lengths within the gut mucosa. For example, single-cell RNA sequencing in combination with laser-capture microdissection or targeted single-RNA-molecule imaging has revealed spatial gradients in gene expression within epithelial populations and mucosal fibroblasts along the axis stretching from the top to the bottom of the mucosa^15–18^. These gradients give rise to functional fine tuning over the micron-scale, and, in the case of morphogen expression can have important consequences in cell development and fate^15–18^. Yet aside from the direct measurement of a handful of genes, the transcriptome wide adaptation of gene expression within cells of the mucosa remains only inferred not directly measured, limiting our understanding of the degree of niche-dependent functional fine tuning of cells within the gut.

In addition to mediating rapid responses to local environmental cues, gut sensing also regulates gene expression, cellular recruitment, and decisions regarding cell fate and development. Over longer timescales, these sensing mechanisms have the potential to reshape the structural and functional features of the gut. A key driver of such remodeling is the gut microbiome, which influences the host by regulating immune cell development, nutrient absorption and metabolism, central nervous system functions, and susceptibility to disease^19,20^. As the largest, most complex, and metabolically active commensal community in the host, the gut microbiome produces a diverse array of small molecules, such as short-chain fatty acids, tryptophan metabolites, as well as secondary bile acids^1,2^. Therefore, host sensing of the activity of the microbiota is one of the ways in which the microbiome may shape gut physiology. For example, lacking the diverse molecular cues provided by the microbiota, germ-free mice have an altered mucosal barrier, metabolism, and altered abundance of gut cell types^21–26^. Nonetheless, the remodeling of the gut by the microbiota has not been systematically characterized across all cell types and compartments; thus, the role that the microbiome and its metabolites play in shaping structural and transcriptomic features of the gut remains unclear.

Image-based transcriptomic methods, with the ability to characterize the expression of large fractions of the transcriptome within single cells within intact tissues, promise the ability to define the sensory gene expression of cell types, place these cell types in their region of origin, and identify subtle spatial variations in expression and, thus, offer a new avenue to address these questions. Indeed, studies leveraging a variety of image-based methods—MERFISH^27,28^, CosMx^29^, or Xenium^30^—have already provided insights into the molecular, cellular, and spatial structure of individual gut regions in health and inflammatory disease^31–34^. Here we apply MERFISH to provide new insights into the cellular and spatial organization of sensation in the murine gut by profiling the expression of a near comprehensive set of murine receptors in 2.1 million cells from 105 cross-sectional slices taken from four regions of the gut, covering both the small and large intestine, in both specific-pathogen-free (SPF) and germ-free (GF) conditions. The resulting atlas provides a rich resource for understanding the cellular compartmentalization of small molecule sensation in the gut, the spatial specialization of these capabilities both across gut regions and within the length of the mucosa, and the modulation of the molecular, cellular, and spatial structure of the gut by the microbiota.

## RESULTS

### Construction of a cellular atlas across the lower mouse GI tract with MERFISH

To define the diversity of cells in the GI tract, their sensory capabilities, regional specialization, potential spatial fine-tuning, and the role of the microbiome on the molecular and cellular structure of the gut, we created a MERFISH library targeting 1,922 genes (**Table S1**, STAR Methods). The first library portion focused on cellular identity and included ∼200 markers of cell types. The second portion focused on the sensory capabilities of these cells and comprised essentially all major classes of annotated receptors in the mouse genome—such as G-protein-coupled receptors (GPCRs), ligand-gated ion channels (LGICs), enzyme-linked receptors (ELRs), nuclear receptors (NRs), pattern recognition receptors (PRRs), complement receptors, and cytokine receptors—as well as their host-derived ligands, including cytokines and ligands from pathways such as Wnt, Hedgehog, Notch, Bmp, and Hippo. Collectively, these genes were targeted with 2,070 39-bit combinatorial barcodes. Additionally, we measured 12 highly expressed cell markers with non-combinatorial barcodes (STAR Methods).

We used MERFISH with this panel to construct an atlas of four regions of the mouse lower gut—ileum, cecum, and proximal and distal colon (**Figure 1A**). These regions were selected because they span both the large and small intestine and have the highest microbial density^35^. Tissue was harvested from male C57BL/6NTac mice aged 8-12 weeks harboring either a specific-pathogen-free (SPF) microbiota or no microbiota (germ-free [GF]), using a fresh-frozen protocol optimized for RNA stability^36,37^ (STAR Methods). In total, gene expression was measured in 105 cross-sectional slices taken from at least 3 mice per region and condition (**Figure 1B**; STAR Methods). As receptor mRNAs can be lowly expressed even when functionally relevant, we developed a method to decrease false positive levels by defining a quality score for each identified RNA molecule, lowering our detection expression limit to ∼1 count per 500 cells (**Figures S1A-S1C**; STAR Methods). Supporting the accuracy of our identified RNAs, markers were identified in the expected spatial locations (**Figure 1B**), average expression per slice correlated strongly with bulk RNA-sequencing from the same regions (**Figure S1C**), and MERFISH measurements from different mice correlated strongly within each region (**Figures S1D**-**S1E**).

**Figure 1.**
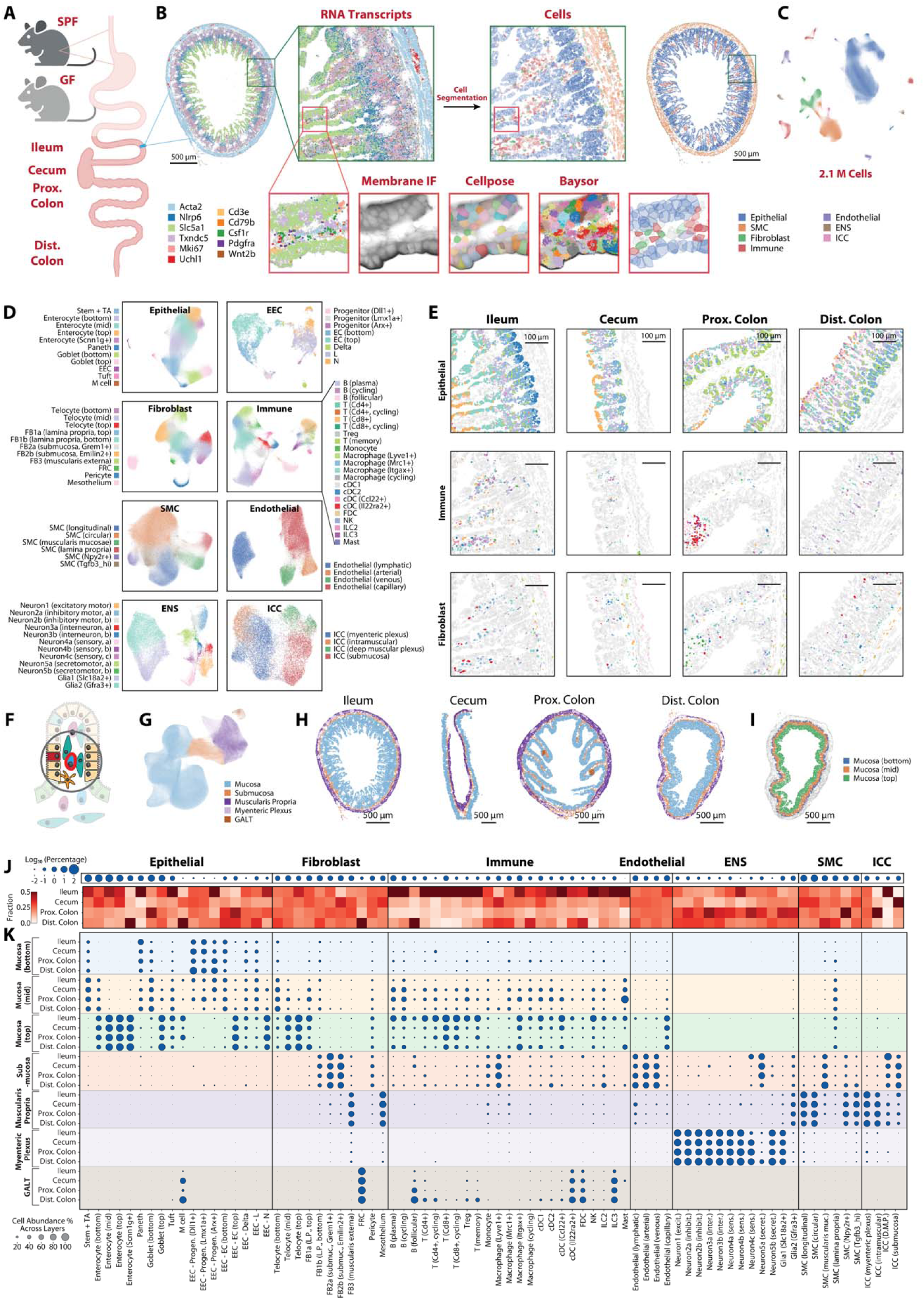
MERFISH defines the cellular diversity and spatial organization of the lower mouse gastrointestinal tract. (A) Illustration of the gut regions profiled with MERFISH in specific pathogen free (SPF) or germ-free (GF) mice. (B) Spatial distribution of 11 out of 1,922 mRNAs (top left) and the boundaries of identified cells colored by major cell class (top right) in an ileal cross-section from a SPF mouse. Images of the distribution of these RNAs, immunofluorescence (IF) against a pan-membrane marker, the boundaries identified via Cellpose, RNAs colored by the cell to which they were assigned by Baysor, and the estimated 3D boundaries of identified cells projected in 2D (bottom, left to right). Scale bar: 500 µm. (C) Uniform manifold approximation and projection (UMAP) of the 2.1 million cells characterized colored by the major cell class. ENS: Enteric nervous system. SMC: smooth muscle cell. ICC: interstitial cell of Cajal. Cells from all regions and microbiome states are included. (D) UMAP of each major cell class colored by the cell types identified within those classes. EEC: enteroendocrine cell. EC: enterochromaffin cell. FB: fibroblast. FRC: fibroblastic reticular cell. cDC: conventional dendritic cell. FDC: follicular dendritic cell. NK: natural killer cell. ILC: innate lymphoid cell. Cells from all regions and microbiome states are included. (E) Spatial distribution of cells in each of the four characterized regions for three major cell classes in SPF mice. Boundaries are colored as in (D) for each class or gray if they are not in the listed major cell class. Scale bars: 100 µm. (F) Illustration of neighborhood analysis approach for dissecting gut layers (STAR Methods). (G) UMAP based on the neighborhood composition for individual cells, colored by the anatomical region. Cells from all regions and microbiome states are included. (H) Spatial distribution of cells in representative slices from each of the four regions in SPF mice colored by neighborhood identity. Scale bars: 500 µm. (I) Spatial distribution of cells in a representative slice of the SPF ileum colored by mucosal region (STAR Methods). (J) The average fractional abundance of each cell type in each region across all slices measured in SPF mice in log-base-10 of the percentage (top) with the normalized fraction of each cell type in each of the four different regions profiled in SPF mice (bottom). (K) The relative abundance of each cell type in each anatomical layer for SPF mice. Relative abundance is normalized to the fractional abundance in the corresponding region in (J).

To segment cells within these measurements, we included with each MERFISH measurement immunofluorescence images of a pan-cell surface marker, the Na^+^/K^+^-ATPase, and used Cellpose^38^ to define the 3D boundaries of cells based on these images (**Figure 1B**; STAR Methods). To refine these boundaries and to identify cells that could not be captured with Cellpose, we used the distribution of the RNAs themselves with Baysor^36^ (STAR Methods). As minor cell segmentation errors could confound our profiling of lowly expressed genes, we developed a segmentation optimization pipeline that uses doublet-detection tools^39^ to provide a measure of the segmentation quality. We used this pipeline to guide segmentation and parameter choices that produced a noticeable improvement in segmentation quality (**Figures S1F-S1K**; STAR Methods). Putative cells generated by this pipeline were then filtered through a combination of volume, RNA copy number, and residual doublet score (STAR Methods) to remove small cell fragments and segmentation errors, producing 2,060,051 cells collectively containing 280 million RNAs.

To manage the effects of gene expression differences due to regional specialization and microbiota dependence, we used Harmony^40^ to jointly embed and label the cells within this atlas (STAR Methods). A first tier of clustering revealed all anticipated cell classes, which then divided into 78 distinct cell populations with additional tiers of clustering (**Figures 1B-1E**; STAR Methods). These clusters showed a high degree of marker purity (**Figure S1L**); co-integrated both across regions and microbiome states (**Figures S1M** and **S1N**); co-integrated with published scRNA-seq measurements (**Figures S1O-S1X**); were found in the expected locations (**Figures 1E** and **S2A-S2B**); and expressed the expected markers (**Figure S2C-S2K**), supporting our segmentation, clustering, and cell-type labeling.

To then define the spatial structure of the gut, we statistically identified anatomical features through recurrent patterns in the composition of the local neighborhood of cells^32^ (**Figure 1F**; STAR Methods). This analysis identified all major gut anatomical structures, including the mucosa, the submucosa, the muscularis propria, the myenteric plexus, and patches of gut associated lymphoid tissue (GALT; **Figures 1G** and **1H**). We then refined this analysis to capture the known zonation of the mucosa by classifying top, middle, and bottom mucosal regions (**Figure 1I**; STAR Methods). This tissue neighborhood mapping allowed us to determine the fraction of each cell type within each anatomical feature across all four regions (**Figures 1J** and **1K**). Importantly, this analysis revealed cell types in the expected regional and anatomical locations, further supporting our analysis.

In summary, MERFISH identified 78 distinct cell populations, characterized their abundances across the four regions of the lower GI tract, and mapped their spatial organization within gut anatomical features, both in the presence and absence of the microbiome. This atlas complements recent spatial omics atlases of portions of the mouse or human gut^4,33,34,41^ by combining high spatial resolution, high cell-type resolution, and broad regional coverage. As we anticipate that this atlas will be a rich resource for the community, we developed a web-based viewer to facilitate access and exploration (Resource Availability).

### Cellular identity and spatial organization are largely conserved across gut regions

The cellular composition of the mammalian gut differs between the small and large intestine and varies along the longitudinal axis within each region. Moreover, cell types can display region-specific expression profiles reflecting specialization to the unique local molecular environment. While there exist several pan-region murine single-cell atlases focused on specific intestinal cell populations^4,5,8,10,12,14^, the extent to which cell identity and spatial organization are conserved across region remains unclear. To address this question, we first examined the cell type diversity in our integrated large-and-small-intestine atlas in SPF mice.

Despite strong regional specialization in epithelial cell function, our atlas defined a diverse set of epithelial cells (**Figure 1D**) with most cell types seen in all four profiled GI regions (**Figure 1J**). We identified the expected maturation of absorptive enterocytes, starting with stem and transit amplifying (TA) cells, and proceeding through increasing maturation from the bottom to the top of the mucosa (**Figures 1D**, **1E**, **S2B** and **S2C**). The joint integration of enterocyte development suggests that, despite the differences in enterocyte function and lamina propria (LP) organization between the small and large intestine, there is a strong, regionally conserved maturation program among enterocytes (**Figure S1M**).

We observed a similar conservation in secretory epithelial cells, with the identification of two divisions of goblet cells (bottom and top), tuft cells, and Paneth cells (**Figures 1D**, **1E**, and **S2C**). We also observed a rich diversity of enteroendocrine cells (EEC; **Figures 1D, S2D**, and **S2E**), with an early progenitor population, and two late progenitor states committed either to enterochromaffin (EC) or delta, L, or N cell development, respectively. Finally, we also observed microfold (M) cells—specialized epithelial cells that line gut associated lymphoid tissues (GALT)—in all four regions with the expected GALT association (**Figures 1J, 1K**, and **S2C**).

We identified four families of fibroblasts (**Figures 1D**, **1E**, **1J**, **1K**, and **S2F**). As there is still some debate regarding fibroblast naming, we adopted telocyte and FB1, FB2, and FB3, as names for these families. As expected, telocytes were identified lining the epithelial layer in the lamina propria (LP) and were marked by *Pdgfra*, *Bmp5*, and *Wnt5a*. FB1 were found within the lamina propria but further away from the epithelium, and were marked by *Sfrp1* and *Wnt2b*; FB2 were found in the submucosa and were marked by *Cd34*, *Wnt2b*, and a variety of atypical chemokine receptors (*Ackr1-4*); and FB3 were found exclusively in the muscularis externa in the vicinity of the myenteric plexus and were marked by a variety of ENS-related genes (e.g., the neuropeptide receptor, *Adcyap1r1*, and nerve growth factor, *Ngf*).

Shared gene expression, spatial location, as well as co-embedding with scRNA-seq (**Figures 1D**, **1E**, **1J**, **1K**, **S1Q-S1T**, and **S2F**) allowed us to embed these fibroblast states in the diverse literature nomenclature^12,42–45^. FB1 corresponds to cells previously referred to as crypt-base fibroblasts, or alternatively *CD90*+, *Fgfr2*+, or *Igfbp5*+ fibroblasts in the LP. FB2 corresponds to cells previously described as trophocytes, *Pi16*+, or *Ackr4*+ fibroblasts. FB3 represents a recently described ENS-associated fibroblast subtype previously reported only in the distal colon^32,46^, which our measurements now reveal in the small intestine as well. Illustrating the impact of our improved segmentation, we were able to resolve the telocyte-FB1 division better than in previous MERFISH studies^32^. Importantly, within the telocyte, FB1, and FB2 fibroblast families, we also identified divisions that were conserved across all four regions. In the case of telocytes and FB1, these subdivisions had a clear spatial organization (**Figures 1D**, **1E**, **1J**, **1K**, and **S2F**). By integrating gene expression, spatial location, and scRNA-seq co-embedding, we conclude that top FB1 likely corresponds to the recently described *Fgfr2*+ fibroblast while bottom FB1 likely corresponds to *Thy1*+ fibroblast in the colon or *Igfbp5*+ fibroblast in the small intestine^12^ (**Figures 1D**, **1E**, **1J**, **1K**, **S1R**, **S1T**, and **S2F**). In addition to these core fibroblast families, we also identified a series of other stroma-like cells that we grouped with fibroblasts, including pericytes, fibroblast reticular cells (FRC), and the mesothelium found, as expected, in the LP, GALT, and a single-layer surrounding the muscularis propria, respectively (**Figures 1D**, **1E**, **1J**, **1K**, and **S2F**).

Similarly, we identified a striking immune diversity (**Figures 1D**, **1E, 1J**, **1K**, and **S2G**). Among lymphoid cells, we observed plasma, follicular, and cycling B cells. Plasma and cycling B cells were found predominantly in the LP while follicular B cells were heavily enriched in GALT, as expected. We also observed a diversity of T cells, including *Cd4*+, *Cd8*+, and cycling versions of both as well regulatory T cells (Treg) and memory T cells (*Ccr7*+)—all predominantly localized within the mucosa. Type 2 and 3 innate lymphoid cells (ILC2 and ILC3) were also present; ILC2s were mainly mucosal while ILC3s were strongly enriched in GALT. Among myeloid cells, we observed monocytes; macrophage subsets expressing *Lyve1*, *Mrc1*, or *Itgax*; multiple conventional dendritic cell populations (including cDC1, cDC2, an activated migratory population marked by *Ccl22* and *Ccr7*^47^, and a recently defined GALT-associated *Il22ra2*+ population^48,49^); follicular dendritic cells (FDC); natural killer cells (NK); and Mast cells. The abundance of immune populations varied dramatically across gut regions, with ileum containing far more immune cells compared to the large intestine (**Figures 1J** and **1K**).

In addition, we defined a diversity of other stromal populations, including smooth muscle cells (SMC), interstitial cells of Cajal (ICC), and endothelial cells (**Figures 1D**, **1E**, **1J**, **1K**, and **S2H**-**S2J**). The SMCs included both longitudinal and circular SMCs in muscularis propria, SMCs found within the lamina propria, and SMCs of the muscularis mucosae. Interestingly, we observed two additional SMC populations marked by high levels of *Tgfb3* and *Npy2r*, respectively (**Figure S2I**). The latter was found lining the boundary between the submucosa and the inner muscular layer, was most prominent in the proximal colon (**Figure 1K**), and based on location, likely represents a recently discovered mouse SMC in the superficial muscularis propria^50^. Its expression of neuropeptide receptors (*Npy2r* and *Cckar*) as well as the neuro-related adhesion GPCR *Adgrd1* suggest a previously unnoticed potential for interactions with EECs and the ENS (**Figure S2I**). Within endothelial cells, we found all four expected divisions: lymphatic, arterial, venous, and capillary, the last of which was the only population appreciably found outside of the submucosa (**Figures 1K** and **S2H**). Finally, we identified multiple populations of interstitial cells of Cajal (ICCs) (**Figure S2J**). ICC was the only major cell class that failed to co-integrate across all four profiled regions (**Figure S1M**), consistent with the established regional specialization^51^. The myenteric plexus ICC was found in all regions while the intramuscular, deep muscular plexus, and submucosal ICCs had more prominent regional enrichment (**Figures 1J** and **1K**).

We also distinguished the expected ENS populations in our atlas. We identified five major neuronal families, which co-embedded well with expression profiles of previous studies^10^ (**Figures 1D**, **1J**, **1K**, **S1U**, **S1V** and **S2K**). Specifically, we identified *Chat*+ excitatory motor neurons (Neuron 1), *Nos1*+ inhibitory motor neurons (Neurons 2a and 2b), *Penk*+ interneurons (Neurons 3a and 3b), *Calcb*+ sensory neurons (also called intrinsic primary afferent neurons [IPAN]; Neurons 4a, 4b, and 4c), and *Glp2r*+ secretory motor neurons (Neurons 5a and 5b). We also identified two glial populations marked by *Slc18a2* and *Gfra3*, respectively.

Neuronal cell bodies are found within two major anatomical regions in the gut: the myenteric plexus, situated between the inner and outer muscular layers; and the submucosal plexus, located in the submucosa. Our measurements complement previous single-cell profiling by unambiguously mapping these neuronal populations to their respective anatomical structures^10^ (**Figure 1K**). As expected, the myenteric plexus contains the greatest diversity of neuronal types, with all identified populations present, while the submucosal plexus consists predominantly of IPANs and secretory motor neurons^52^ (**Figure 1K**). Interestingly, the neuronal composition of these plexuses varies across gut regions. Most notably, sensory neuron 4c was prominent in the submucosal plexus in the ileum but not the large intestine, while secretory motor neuron 5a was found almost exclusively in the submucosal plexus in almost all regions except the distal colon. Our measurements also revealed distinct spatial organization of glial cell populations. *Gfra3*+ glia was found outside the myenteric plexus—in the surrounding muscle layer, the submucosa, and, in small numbers, the mucosa—consistent with previous findings^53^. In contrast, *Slc18a2*+ glia were strongly enriched in the myenteric plexus. As these cells have been proposed to have neurogenic potential^53^, their localization may have implications for the regenerative capacity within the myenteric plexus.

### MERFISH reveals a potential mouse homolog of the human BEST4+ enterocyte

While examining cell populations identified in our atlas, we uncovered an enterocyte population that has not been clearly described in mouse to our knowledge. This population was characterized by the expression of all three subunits of the epithelial sodium channel (*Scnn1a*, *Scnn1b*, and *Scnn1g*) and was most clearly marked by *Scnn1g*; hence, we termed these *Scnn1g*+ enterocytes (**Figures 2A**). *Scnn1g*+ enterocytes were found predominantly in the distal colon (**Figure 1J**), almost exclusively in the top mucosa (**Figures 1K** and **2B**), and expressed markers shared with mature enterocytes (e.g., *Apob* and *Slc5a1*, **Figures 2C** and **S2C**), suggesting that *Scnn1g*+ enterocyte represents a subpopulation of distal colon mature enterocytes. Importantly, beyond the expression of the epithelial sodium channel, these enterocytes display transcriptional signatures indicative of unique functions. For example, high *Nt5e* expression suggests a role in extracellular AMP metabolism; *Ptger3* indicates a sensitivity to prostaglandin signaling; and the expression of *Bmp3*—rather than *Bmp8a*, which is expressed in other mature enterocytes—suggests a unique role in morphogen signaling (**Figures 2C** and **S2C**).

**Figure 2.**
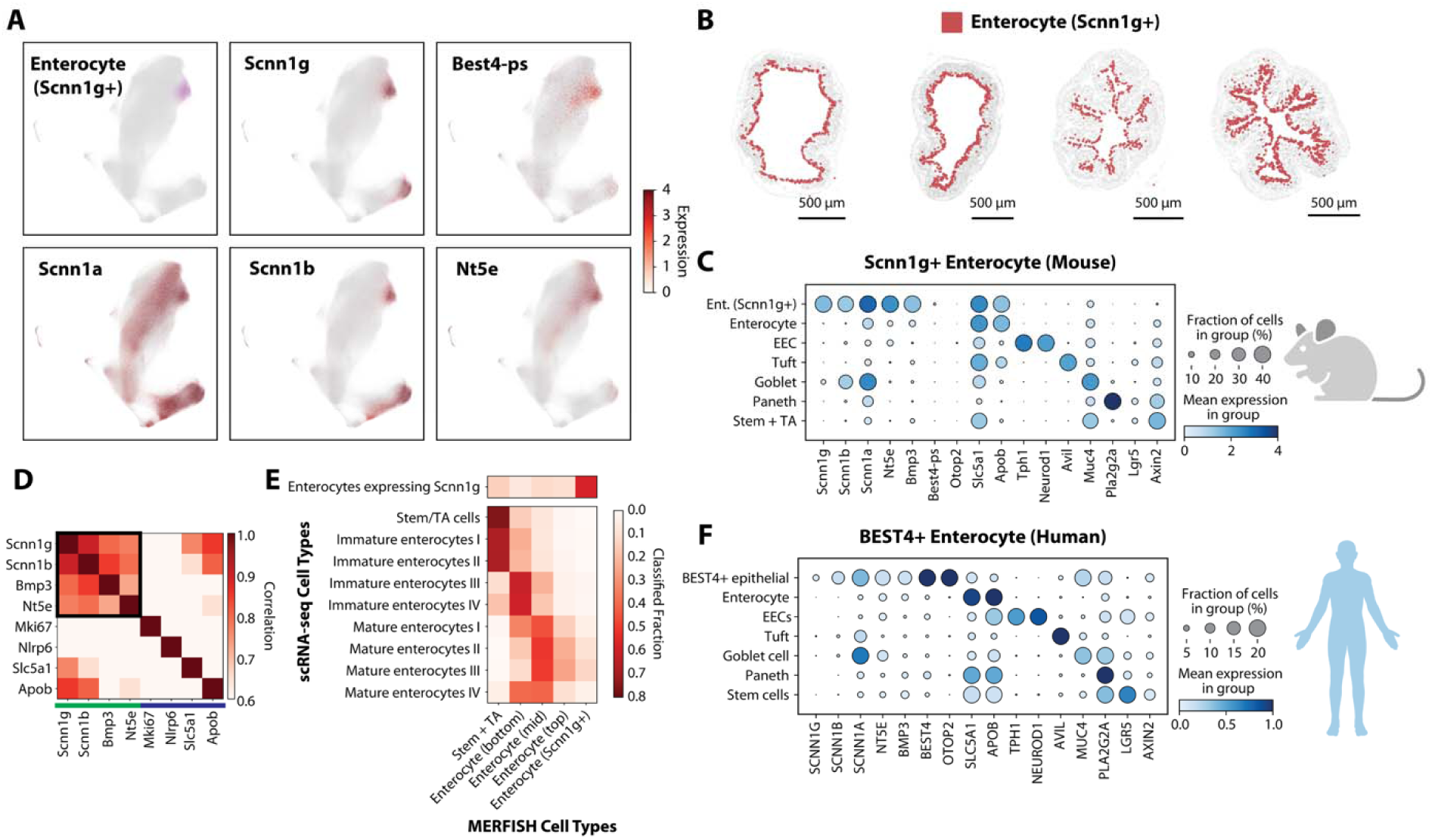
MERFISH defines a potential mouse homolog to the human BEST4+ enterocyte. (A) UMAP of epithelial cells colored by the Enterocyte (*Scnn1g*+) cluster (upper right) or the logarithmic normalized expression of the indicated marker genes. (B) Spatial distribution of cell centroids for *Scnn1g*+ enterocytes (red) in representative slices taken from distal colon in SPF mice. Gray represents all other cells. Scale bars: 500 µm. (C) Average gene expression within key epithelial populations from SPF mice. Color indicates the logarithmic normalized expression with the cluster and the dot size indicates the fraction of cells that express at least one count of the listed gene. (D) Pairwise Pearson correlation coefficients for the gene expression within all enterocytes in published scRNA-seq^33^. Green and purple lines indicate markers of different epithelial populations, and boxed genes represent co-variation of markers of *Scnn1g*+ enterocytes. (E) Fraction of enterocytes measured with scRNA-seq^33^ that are mapped to each of the MERFISH epithelial cell type labels based on co-embedding (bottom; STAR Methods). scRNA-seq enterocytes with at least one copy of *Scnn1g* mRNA were treated as a separate group for mapping to MERFISH (top). (F) As in (C) but for human intestinal epithelial populations published previously^5^.

To provide further support for this population, we examined prior murine gut single-cell studies and found evidence for a subpopulation of mature enterocytes that co-express all subunits of the epithelial sodium channel as well as *Nt5e* and high levels of *Bmp3* in a recent study^33^ (**Figures 2D** and **S3A**). Although this subpopulation had not been explicitly identified in that dataset, co-embedding revealed that our *Scnn1g*+ cluster aligned with the published cells expressing *Scnn1g*, confirming the presence of this subpopulation (**Figures 2E**, **S3B**, and **S3C**). The sensitivity of MERFISH, along with the large number of cells profiled in our study, may have enabled us to cluster and identify this population in our atlas where smaller datasets could not.

We next asked whether the *Scnn1g*+ enterocytes in mice resembled epithelial populations in other organisms. Using a recent human GI single-cell atlas^5^, we found that the human BEST4+ enterocyte displays a similar expression profile to the *Scnn1g*+ enterocyte in mice (**Figure 2F**). Notably, the human BEST4+ enterocyte was the only population to express appreciable levels of all three subunits of the epithelial sodium channel and was also the major source of NT5E and BMP3 (**Figure 2F**). Moreover, these populations shared striking spatial similarities: BEST4+ enterocytes are located at the top of the mucosa^54^ and are primarily found in the distal colon^54^, mirroring the distribution of *Scnn1g*+ enterocytes in mouse (**Figures 1J**, **1K** and **2B**).

Although BEST4+ enterocytes have been observed in a variety of organisms, including macaque, pig, rabbits, and zebrafish^54^, it has been suggested that mice lack a homolog due to an in-frame stop codon that renders *Best4* a pseudogene^54,55^. However, we hypothesized that if the *Scnn1g*+ enterocyte is a homolog of the BEST4+ enterocyte, residual regulatory mechanisms might still drive *Best4-ps* expression. Indeed, bulk RNA sequencing detected *Best4-ps* expression in the mouse distal colon at a level consistent with our MERFISH data (**Figure S3D**), and, importantly, the *Scnn1g*+ enterocyte was the only population to express *Best4-ps*, albeit at low levels (**Figures 2A** and **2C**). Collectively, these observations suggest that the *Scnn1g*+ enterocyte may represent the mouse homolog of the human BEST4+ enterocyte.

Nonetheless, our data also indicate a potential functional divergence between these populations: in addition to the non-functional *Best4-ps*, the mouse homolog of the only other commonly cited marker of human BEST4+ cells included in our MERFISH panel, OTOP2, was not expressed by the corresponding mouse population (**Figures 2C** and **2F**).

### Cells in the intestinal mucosa are spatially organized along the crypt-to-villus axis

The intestinal mucosa is highly organized, with cell abundance gradients along the axis from the submucosa to the lumen^4,34,41^. Previous studies have lacked either full regional coverage, cellular resolution, or have focused on limited cell types^4,34,41^, leaving the spatial organization of mucosal populations across GI regions incompletely defined. Therefore, we next used our atlas to provide a cellular-scale map of the organization of the intestinal mucosa.

One challenge to defining this organization is the imperfect and variable orientation of crypts and villi that arises as a natural result of sectioning. To address this challenge, we were inspired by the strong gene expression variation that occurs within epithelial populations as they mature from the crypt to the top of the mucosa^15,16,18^, which we reasoned could serve as a natural mucosal ruler, defining mucosal location via gene expression instead of absolute position. Indeed, multiple epithelial genes showed clear zonation in expression (**Figure 3A**), and we observed a clear continuous gradient in gene expression in epithelial cells from stem to mature enterocytes (**Figure 3B**). To quantify the transcriptomic variation in enterocytes along this developmental axis, we performed pseudotime analysis on the co-integrated stem, TA cells, and enterocytes from all profiled regions and found excellent agreement between this pseudotime and both the continuous axis of transcriptomic variation captured in our epithelial UMAP (**Figures 3B**-**3E**; STAR Methods) and the spatial location of enterocytes within all four regions (**Figures 3F** and **3G**). Importantly, this clear agreement underscores the remarkable fidelity between epithelial maturation and mucosal position (**Figure 3G**).

**Figure 3.**
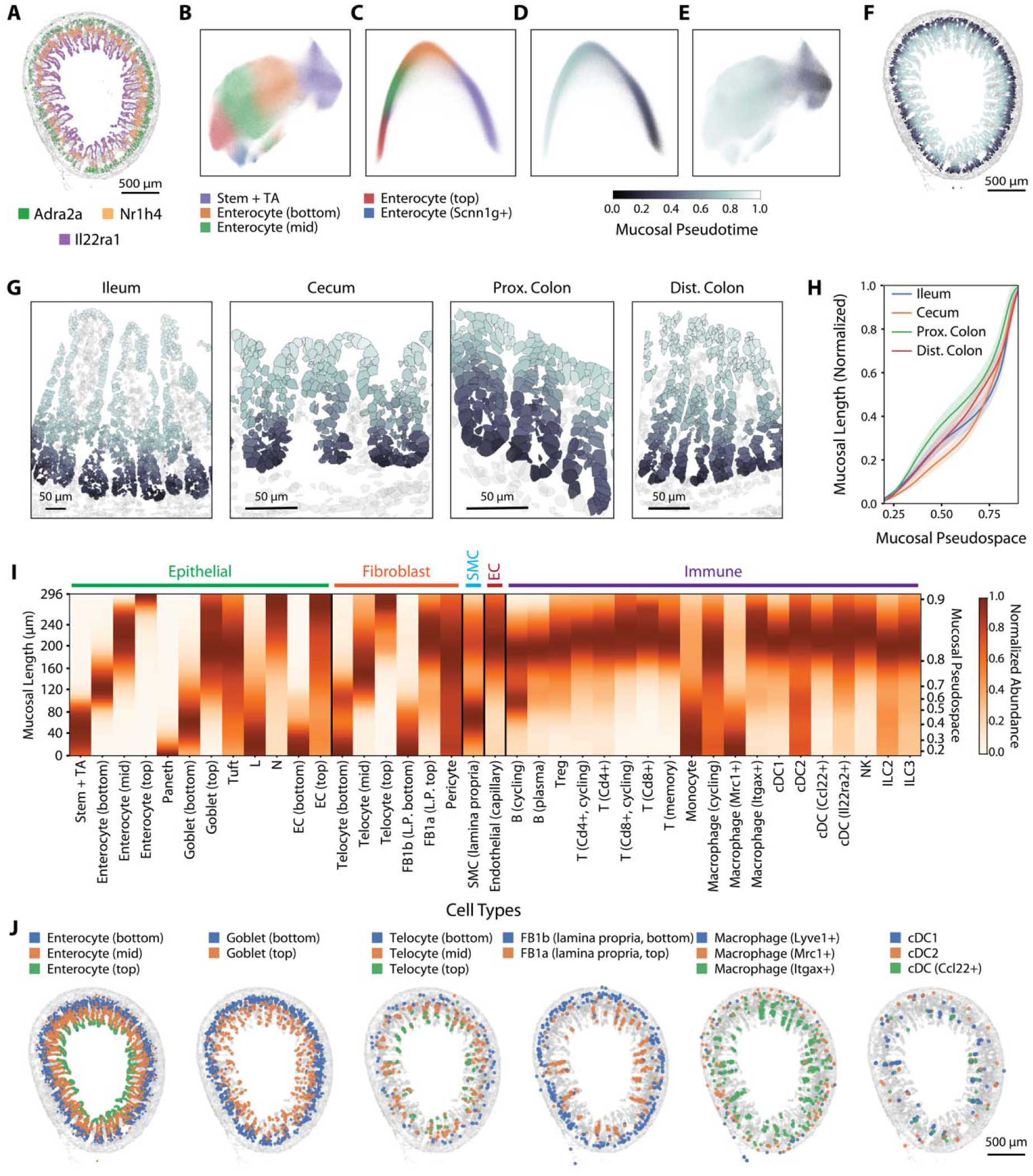
Cells within the intestinal mucosa are organized along the mucosal axis. (A) Spatial distribution of three example mRNAs (color) over all mRNAs (gray) measured in a representative slice of the ileum from SPF mice. Scale bar: 500 µm. (B) UMAP representation of all absorptive epithelial cells in SPF mice colored by major populations. (C) The first two diffusion components of the epithelial cells in (B) colored as in (B). (D) As in (C) but colored by the pseudotime variable that captures epithelial maturation. (E) As in (B) but colored by pseudotime. (F) As in (A) but with all absorptive epithelial cells colored by pseudotime. Scale bar: 500 µm. (G) The spatial distribution of cells in representative mucosal regions from four slices from the listed regions in SPF mice. Gray patches represent non-epithelial cells, and colored patches represent absorptive epithelial cells colored by pseudotime. Scale bars: 50 µm. (H) The relationship between mucosal pseudospace and actual mucosal length, normalized by the mucosal thickness of the specified region. Lines represent the average across all slices and shaded regions represent the 95% confidence interval (slice numbers in **Figure S2A**). (I) Average distribution of all major mucosal cell populations along the length of the ileal mucosa of SPF mice determined by mucosal pseudospace (left) or mucosal distance from the crypt (right). Only mucosal populations with sufficient abundances are shown (STAR Methods). (J) Spatial distribution of cell centroids for the listed example populations (color) on top of all cells (gray) in a representative slice of the SPF ileum. Scale bar: 500 µm.

To then use this gene expression ruler to define spatial positions within the mucosa, we developed a mucosal pseudospace axis by propagating the pseudotime value from the nearest enterocytes to all other mucosal cells (STAR Methods). We then calibrated this mucosal pseudospace axis to actual mucosal position by calculating a cumulative distribution of epithelial cell numbers along this axis and estimating the average thickness of the mucosa for each region (**Figures 3H** and **S4A**; STAR Methods). All regions showed a very similar progression of the pseudospace axis along the length of the mucosa; however, anatomical differences between the large and small intestine were clear. For example, pseudospace grew linearly with mucosal position in the ileum until roughly 80 µm, after which the pseudospace value increased quickly over a short mucosal length, before growing very slowly over a large stretch of the mucosa that corresponds to the mature villus region. This transition at 80 µm reflects the established transition between crypt and villus and underscores the rapidity of transcriptomic remodeling at this transition (**Figures 3G**, **3H**, and **S4A**).

Using this pseudospace axis, we computed the spatial distribution of all mucosal cells, revealing a rich, cell-type specific organization in all four regions (**Figures 3I, 3J,** and **S4B**-**S4D**; **Table S2**). Supporting this analysis, we observed the expected zonation of stem/TA, bottom, mid, and top enterocytes as well as other cell type divisions already named based on spatial location, e.g. bottom and top goblet, EC, FB1, and telocytes (**Figures 3I** and **S4B**-**S4D**). Interestingly, we noted subtle differences in the extent of bottom, mid, and top enterocytes as well as bottom, mid, and top telocytes between all four regions (**Figures 3I** and **S4B**-**S4D**), suggesting region-specific variation in the pacing of enterocyte development perhaps shaped by regional differences in the spatial distribution of the mucosal morphogens secreted by telocytes^43^.

This analysis also revealed a rich and differential organization of the immune system across gut regions. In the ileum, most immune populations were seen enriched at the mid-to-top villus. However, there were notable exceptions (**Figure 3I**). Monocytes were found predominantly within the crypts, possibly reflecting their recent recruitment from blood vessels in the submucosa; macrophages were arrayed from bottom to top reflecting the *Mrc1*+ to *Itgax*+ functional maturation^32,56^; and cDC2, while predominantly in the villus, contained a small population in the crypts (**Figures 3I** and **3J**). Interestingly, this strong top mucosal enrichment in the ileum was not as prominent in all regions of the large intestine. While *Cd8*+ T cells and *Itgax*+ macrophages remained the populations closest to the crypt top, possibly reflecting their roles in microbiome interactions, most other populations were found lower in the crypts. The cecum was a particular outlier with most immune populations predominantly found in the base of the crypt (**Figures S4B**-**S4D**).

### Mucosal location modulates gene expression in diverse cell populations

In addition to the cellular scale organization of the mucosa, multiple studies have suggested that some mucosal cell populations adjust their gene expression along the length of the mucosa^15–18,34,57^. However, these studies had either limited cellular resolution or focused on specific cell types; thus, the general degree to which cells continuously adjust their gene expression based on their location in the mucosa and the fidelity between gene expression and location remain poorly defined.

Therefore, we used our atlas to examine how mucosal cell populations modulate their gene expression as a function of location within the mucosa. As expected, our data show clear spatial gene expression variation in cell types where such gradients have been well established, including epithelial cells, telocytes, and macrophages (**Figures 3A**, **4A**, and **4B**). To determine whether other mucosal populations also adjust their gene expression based on position, we projected pseudospace values onto both our MERFISH gene expression UMAPs and, as a control, published scRNA-seq UMAPs, using our co-embedding to impute pseudospace values for scRNA-seq cells (**Figures 4C**, **4D**, and **S5A-S5F**; STAR Methods). These projections revealed that location-dependent fine-tuning of gene expression is a common feature across most mucosal cell types, with clear co-variation between UMAP features and pseudospace in goblet cells, EECs, telocytes, FB1 fibroblasts, pericytes, and capillary endothelial cells. Furthermore, these results suggest that the location-based clusters we identified, e.g. bottom, mid, and top telocytes, represent discrete approximations of a continuous gene expression gradient. For the purposes of this analysis, we therefore regrouped all such divisions.

**Figure 4.**
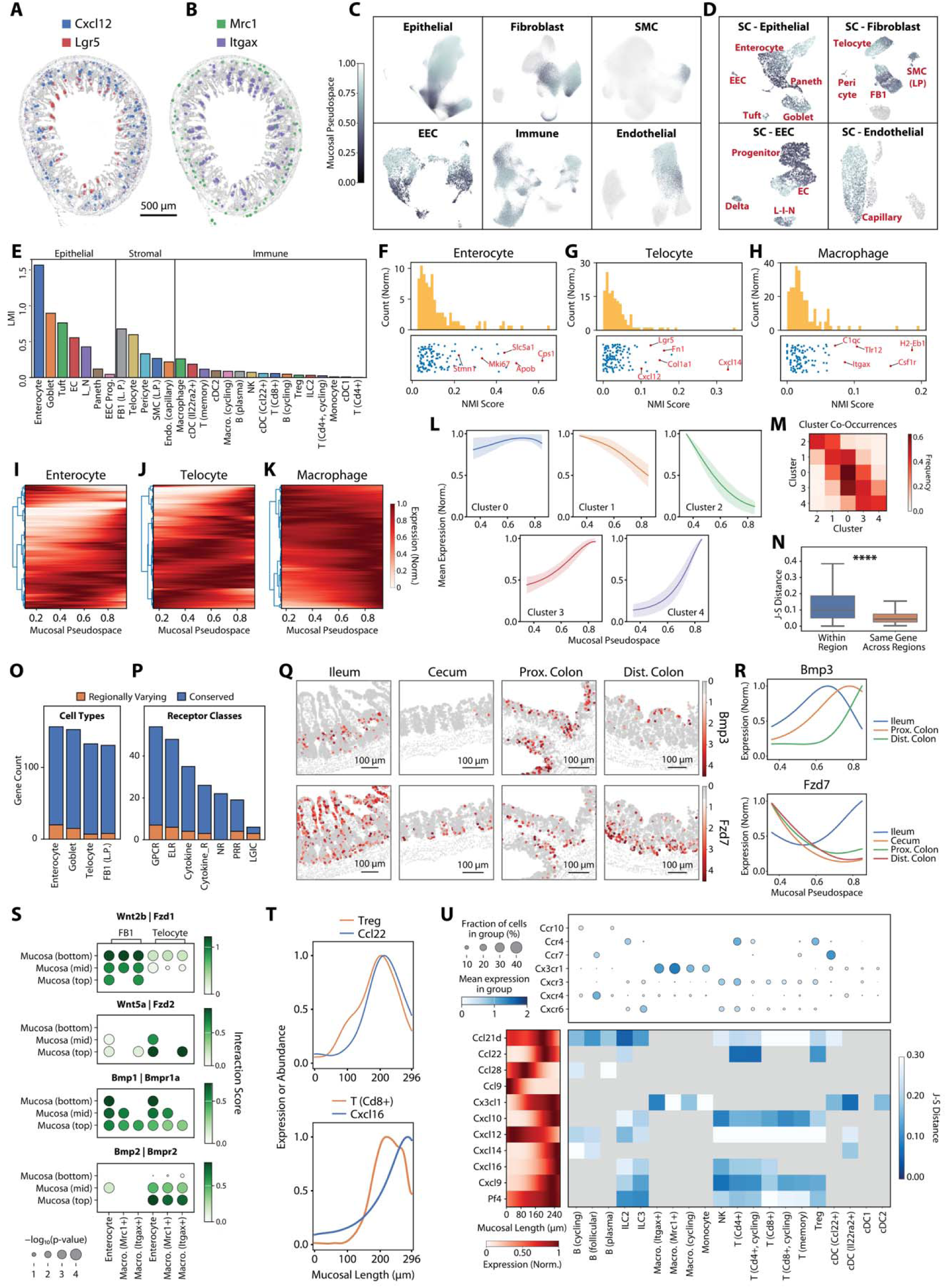
Position along the mucosal axis shapes the gene expression in most mucosal cell types. (A) Spatial distribution of two example mRNAs (color) expressed in telocytes over all mRNAs expressed in all cells (gray) measured in a representative slice of the ileum from SPF mice. Scale bar: 500 µm. (B) As in (A) but for two example mRNAs expressed in macrophages. (C) UMAP representation of all cells of the listed major classes in SPF mice colored by mucosal pseudospace if found within the mucosa; gray if not. (D) UMAP representation of major cell classes measured with scRNA-seq colored by imputed mucosal pseudospace (STAR Methods). (E) Latent mutual information (LMI) between the transcriptome and mucosal pseudospace for listed mucosal cell populations in the ileum of SPF mice. (F) Histogram of the normalized mutual information (NMI) scores for all genes in enterocytes in the ileal mucosa of SPF mice (top) with the distribution of NMI for individual genes (bottom). Notable high NMI genes are listed. (G) As in (F) but for all telocytes. (H) As in (F) but for all macrophages. (I) Average expression of individual genes versus mucosal pseudospace for all enterocytes in the ileal mucosa of SPF mice (right) sorted via hierarchical clustering (left). (J) As in (I) but for all telocytes. (K) As in (I) but for all macrophages. (L) The average gene expression for the five identified clusters of genes showing similar dependence of expression on mucosal pseudospace. Solid lines represent averages and shaded areas represent the 25% to 75% quantile across all slices (slice numbers in **Figure S2A**). (M) The frequency with which genes clustered into one spatial pattern are identified with another spatial pattern in another region. (N) The distribution of Jensen-Shannon (J-S) distances between the spatial patterns for individual genes within a region and between the same gene in different regions (STAR Methods). Line: median. Box: 25% - 75% quantile. Whiskers: 0% - 95% quantile. **** indicates a p-value less than 10^-4^ as determined by a t-test. (O) The number of genes that are conserved (blue) or varying (orange) in their spatial distributions across regions, among all genes expressed in at least two regions within the listed cell populations (STAR Methods). (P) As in (O) with genes tabulated not by cell population but by receptor category. (Q) Two example mRNAs expressed in enterocytes that change their spatial distribution between regions. Scale bars: 100 µm. (R) Average expression of the genes in (Q) versus mucosal pseudospace for the regions in which the gene is expressed above background (STAR Methods). (S) Spatially prioritized receptor ligand interaction scores (STAR Methods) between mucosal fibroblasts (FB1 and telocytes) and different epithelial and macrophage populations. Dot size represents significance while color indicates the interaction score. (T) Average abundance of Treg (orange, top) or *Cd8*+ T cells (orange, middle) versus the average expression across all mucosal cells of chemokines known to attract these cells (*Ccl22*, top; *Cxcl16*, bottom). (U) Average expression of key chemokine receptors in the listed mucosal cell populations (top). Average spatial expression of chemokines across all mucosal cells versus mucosal position (left). J-S distance (color) between the chemokine expression profile and the spatial distribution of cell types expressing the cognate chemokine receptor. Gray indicates that the corresponding cell type does not express the specific chemokine receptor above background levels (STAR Methods). Included are only cells from the ileal mucosa of SPF mice.

To quantitatively compare how different cell populations fine-tune their gene expression based on location, we next calculated the mutual information (MI) between pseudospace and gene expression within each population (**Figures 4E** and **S5G**-**S5I**). We chose MI over other methods as it does not assume a specific form of relationship with position, and we used the recently introduced latent mutual information (LMI) to address the computational challenges of calculating MI for thousands of genes^58^ (**Table S3**). Epithelial cells showed the greatest LMI of any cell class across all four regions, with enterocytes, as expected, showing the largest LMI among all populations. Among other epithelial cells, goblet cells showed substantial location-dependent variation, as did Tuft cells and ECs (**Figures 4E** and **S5G**-**S5I**). Importantly, this analysis revealed extensive stromal remodeling based on location (**Figures 4E** and **S5G**-**S5I**). Telocytes exhibited the greatest LMI, followed closely by FB1, with vascular cells (pericytes and capillary endothelial cells) also showing appreciable location-dependent variation. In contrast, LP SMCs had consistently low LMI across regions. Finally, immune populations generally had modest LMI values, indicating limited spatial fine-tuning of expression among the genes we profiled (**Figures 4E** and **S5G**-**S5I**). While memory T cells in the ileum showed one of the highest LMIs among immune cells, consistent with recent studies^34^, the values were still lower than most epithelial or stromal populations (**Figure 4E**).

To identify genes driving this spatial patterning, we next calculated normalized mutual information (NMI) between gene expression and pseudospace (**Table S3**). Supporting this analysis, we noted that the relative range of NMI values reflected the LMI hierarchies, and top NMI genes aligned with known biology (**Figures 4F**-**4H** and **S5J**-**S5N**). For example, *Mki67*—a marker of dividing cells—scored highly in dividing epithelial cells (enterocytes, goblet cells, and EECs) but not stromal or immune cells. Similarly, other high NMI genes included *Apob* (enterocytes), *Lgr5* (telocytes), and *Itgax* (macrophages), consistent with their spatially restricted expression in the top mucosa (**Figures S2C**, **S2F**, and **S2G**). Broadly, high NMI genes suggested functional fine-tuning with spatial location along the mucosa, such as ECM modulation by telocytes (*Fn1* and *Col1a1*) and differential MHCII expression and PAMP detection by macrophages (*H2-Eb1* and *Tlr12*; **Figures 4G** and **4H**).

We next explored spatial gene expression patterning along the mucosa and its conservation across regions, focusing on four high-LMI populations: enterocytes, telocytes, FB1s, and macrophages. All showed a diversity of spatial patterns among hundreds of genes (**Figures 4I**-**4K** and **S5O-S5P**), indicating wide-spread spatial fine-tuning of expression. Hierarchical clustering revealed five major expression patterns: constant (cluster 0), off-on or on-off (clusters 2 or 4), or modest decreases or increases along the mucosal axis (clusters 1 and 3; **Figures 4L** and **S5Q**). A few genes, mainly among enterocytes, peaked at intermediate pseudospace values (e.g. *Nr1h4*; **Figure 3A**). However, genes with this spatial expression pattern were too rare to form a separate cluster.

To explore regional conservation, we quantified how often genes switched spatial expression clusters across regions, considering genes that are sufficiently expressed in at least two regions (**Figure 4M**). Cluster switching was rare and mostly occurred between similar patterns (e.g., cluster 1 to cluster 2 or cluster 0 to cluster 1; **Figures 4L** and **4M**), indicating that spatial gene expression is largely conserved within each cell population across gut regions. A cluster-free analysis using the Jensen-Shannon (J-S) divergence as a measure of distance between spatial patterns further supported this conclusion (STAR Methods). While gene pairs within a cell type showed diverse spatial patterns, the same gene across regions had consistently lower J-S distances, reinforcing regional conservation of spatial expression patterns (**Figure 4N**). Using J-S distance, we identified a small set of outlier genes with large regional differences in spatial expression patterns within a given cell type (**Figure 4O**; STAR Methods). While these genes were not enriched for specific receptor categories (**Figure 4P**), some, like key morphogens and their receptors such as *Bmp3* and *Fzd7* in enterocytes, may play key roles in shaping regional variation (**Figures 4Q** and **4R**).

We next explored how spatial fine-tuning of gene expression impacts mucosal organization and interactions, focusing on receptor-ligand interactions along the pseudospace axis. Indeed, receptor-ligand analysis performed on different mucosal zones revealed a variety of zonal morphogen interactions that may represent key mechanisms in shaping cell identities along the mucosa (**Figures 4S** and **S5R**; **Table S4**; STAR Methods). Beyond interactions between mucosal zones, we reasoned that the pseudospace axis could help generate spatially prioritized hypotheses from receptor–ligand interactions. To investigate the role of chemokine gradients in organizing motile immune cells along the mucosa, we examined spatial chemokine expression patterns and receptor profiles. For instance, all T cells express *Cxcr3*, whose ligand *Cxcl16* peaks at the tip of the ileal villus (**Figures 4T** and **4U**), consistent with the enrichment of T cells in that location (**Figure 3I**). In contrast, *Cd4*+ T cells, including Tregs, localize to slightly lower mucosal pseudospace values than *Cd8*+ T cells, perhaps due to the expression of *Ccr4* by *Cd4*+ T cells, but not *Cd8*+ T cells. The ligand of *Ccr4*, *Ccl22*, peaks lower in the villus, possibly driving the lower mucosal localization of *Cd4*+ T cells (**Figures 3I**, **4T**, and **4U**).

Inspired by this observation, we reasoned that the spatial similarity between chemokine expression and the distribution of receptor-expressing cells could serve as a quantitative metric to prioritize receptor-ligand interactions influencing cellular organization. To formalize this notion, we computed the J-S distance between the spatial distributions of all chemokines and their cognate receptor expressing cell types, generating a rich set of hypotheses for spatially driven interactions and their regional differences (**Figures 4U** and **S5S-S5U**). For example, this analysis suggests that *Ccl21d* and *Pf4*; *Cxcl9* and *Cxcl10*; and *Ccl22* and *Cxcl9* play more prominent roles than other chemokines in shaping the distributions of ILC2, NK cells, and *Cd4*+ T cells in the ileum, respectively. Similar trends emerged in the large intestine, most notably that chemokines enriched at the crypt base in the cecum may drive the preferential localization of immune populations to this region (**Figures S5S-S5U**). More broadly, quantitative comparisons of gradients in gene expression and cell type distributions, such as J-S distance, may prove valuable for prioritizing the rich interaction hypotheses generated from spatial transcriptomic data.

### Spatial expression variation identifies properties of interferon response and rare cell states

After characterizing gene expression variation along the mucosal base-to-top axis, we next investigated whether similar variations exist along the circumferential axis in cross-sectional slices. We developed a computational method to define circumferential coordinates and detect potential localized spatial gene expression variability (STAR Methods). In all four gut regions, this analysis identified a series of epithelial genes that showed clear spatial patchiness in a subset of slices (**Figure 5** and **Table S5**). These genes fell into two categories. The first included *Dhx58*, *Ddx58*, and *Ifih1*, all of which are interferon-stimulated genes (ISGs). In the ileum, we observed co-expression of these genes within individual villi, with expression patches often spatially isolated and confined to one or two adjacent villi (**Figure 5A**). In the cecum, proximal colon, and distal colon, co-expression was instead seen within small groups of adjacent crypts (**Figures 5B-5D**). Previous studies have shown that certain ISGs, such as *Ifit1* and *Isg15*, are expressed in epithelial cells within individual villi in the mouse small intestine in an interferon-lambda-dependent and microbiota-dependent manner and suggest that such ISG patches modulate susceptibility to enteric viral infection^59,60^. Our observations now suggest that such patchy expression extends to these additional genes and also exists in the large intestine.

**Figure 5.**
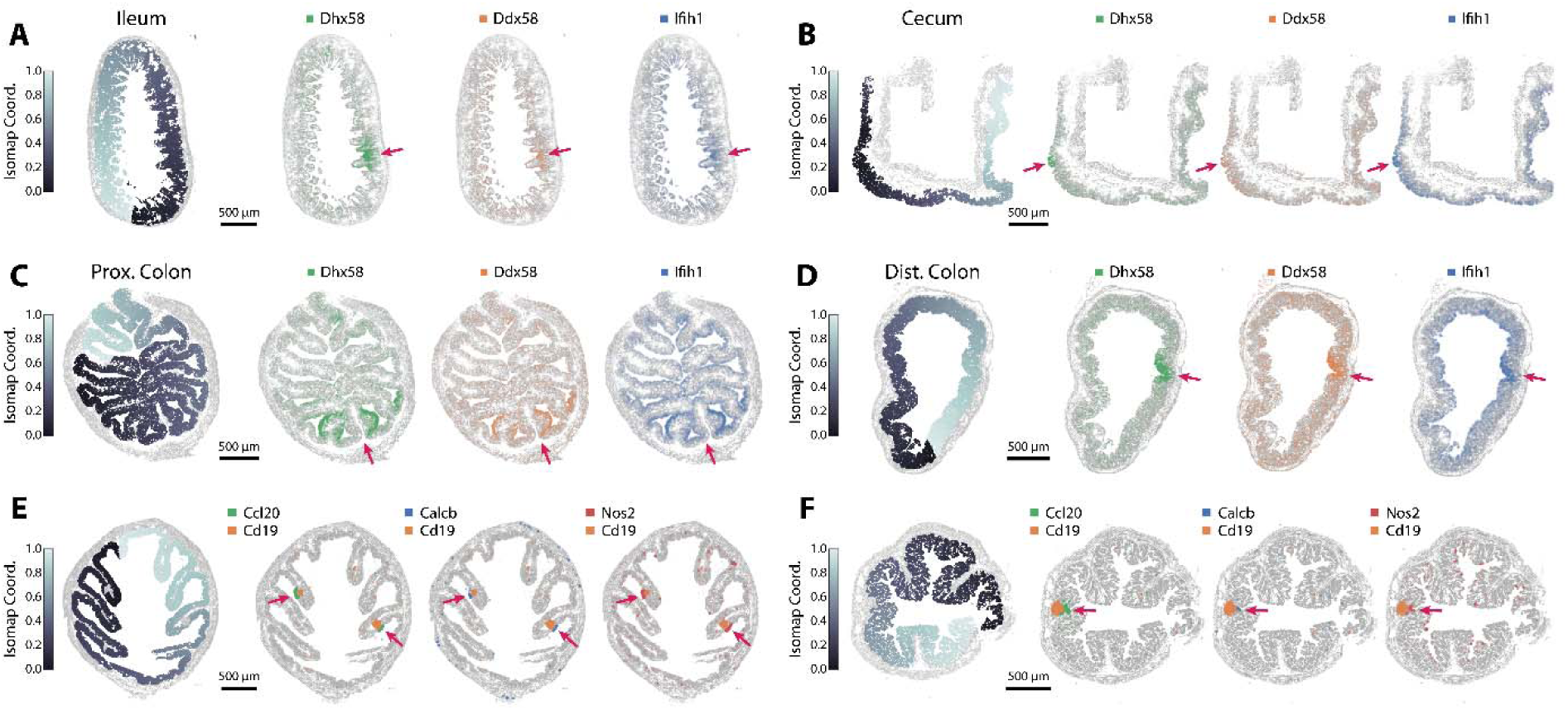
Spatially heterogeneous gene expression along the circumferential gut axis reveals properties of ISG expression and follicle-associated epithelial states. (A) Spatial distribution of mucosal cells colored by circumferential Isomap coordinates (left; STAR Methods) on top of non-mucosal cells (gray) with the spatial distribution of RNAs (right) colored by the listed genes on top of all RNAs (gray) in an example ileum slice from SPF mice. Arrows mark a patch of coordinated interferon-stimulated gene (ISG) expression within epithelial cells. Scale bar: 500 µm. (B) As in (A) but for the cecum from an SPF mouse. (C) As in (A) but for the proximal colon from an SPF mouse. (D) As in (A) but for the distal colon from an SPF mouse. (E) Similar to (A) but for the proximal colon in an SPF mouse. Arrows mark a patch of genes expressed in follicle-associated epithelial cells. (F) As in (E) but for the distal colon from an SPF mouse.

The second category of genes was also found in epithelial cells and included chemokine *Ccl20*, inducible nitric oxide synthase *Nos2*, and calcitonin signaling gene *Calcb*. We observed rare but distinct patches of these genes in the proximal and distal colon (**Figures 5E** and **5F**). Unlike the ISG patches, which lacked structural correlates, these genes were consistently expressed in non-M-cell, follicle-associated enterocytes located on the surface of small GALT patches (as marked by *Cd19*; **Figures 5E** and **5F**). Therefore, these cells likely represent a distinct enterocyte state induced in the presence of GALT, which escaped detection via standard clustering because of their rarity. Importantly, *Ccl20* has been implicated in recruiting *Ccr6*+ B cells during early GALT formation^61^, suggesting a potential role for this rare follicle-associated epithelial state.

### Regional variation in gene expression may shape sensory capabilities and microbial metabolite response

We next explored the spatial, regional, and cell-type variation in genes associated with small molecule sensing. While our initial Harmony-based embedding enabled joint cell-type labeling across regions and microbiome states, it purposefully obscured regional expression differences (**Figure 1C**). Removing batch correction revealed a hierarchy of regional specialization across major cell classes (**Figures 6A** and **6B**; STAR Methods). As expected, epithelial cells showed the greatest regional variation, with pronounced differences across most regions except cecum and proximal colon, which maintained a degree of partial co-embedding (**Figures 6A** and **6B**).

**Figure 6.**
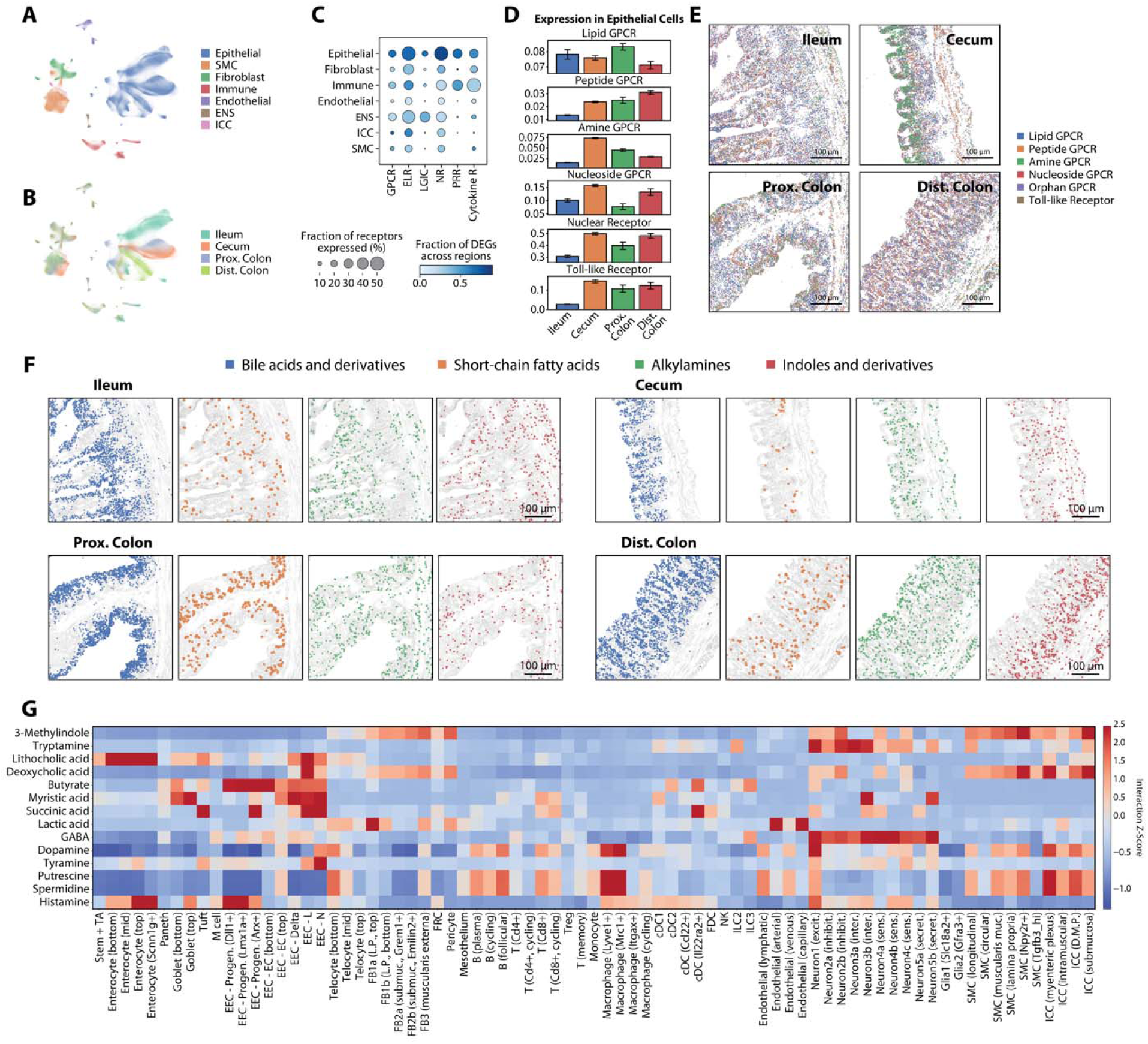
Differential expression of sensory genes may shape regional specialization and interaction with microbially derived small molecules. (A) UMAP representation of all cells imaged in all four regions of SPF mice without Harmony batch correction, colored by major class. (B) As in (A) but colored by region. (C) Regional expression of receptor categories across major cell classes. Circle size represents the fraction of receptors in each category expressed by at least one cell type within each major cell class in at least one region. Color indicates the fraction of these receptors that are differentially expressed across regions (STAR Methods). (D) Average expression of receptor categories within all epithelial cells across regions in SPF mice. Bars and error bars represent the mean and standard error across slices (slices in Figure S2A). (E) Spatial distribution of RNAs encoding receptors of the listed classes in different regions from SPF mice. (F) Spatial distribution of RNAs encoding receptors for specific categories of microbiota-derived metabolites in a representative slice from each region of the SPF mouse. Scale bar: 100 µm. (G) Predicted interaction strength (represented as z-scores across cell types; STAR Methods) between specific microbiota-derived metabolites and gut cell types in all regions of the SPF mouse.

To quantify these variations, we identified genes differentially expressed within a given cell type in one region compared to the same cell type across all other regions (**Table S6**; STAR Methods). This analysis revealed that across all major cell classes, epithelial cells showed the most extensive regional specialization, with over 50% of expressed receptors showing differential expression in at least one region (**Figure 6C**). This degree of specialization is unsurprising given the substantial variation in the small molecule environment across regions and is consistent with prior scRNA-seq findings^5,8,33^. Nonetheless, our work extends this previous understanding by revealing a surprising degree of specialization across many other cell populations. Immune cells, fibroblasts, cells of the ENS, and, to a lesser degree, ICCs, also exhibited differential regional expression within large fractions of their expressed receptors (**Figure 6C**). In contrast, endothelial cells and SMCs had both more modest levels of sensory gene expression and regional specialization (**Figure 6C**).

To explore whether this regional specialization aligns with the shifting small molecule landscape or microbial load along the GI axis, we examined the average expression levels of specific subsets of GCPRs responsive to common gut small molecules (lipids, peptides, amines, nucleosides) as well as receptors responsive to microbial cues (toll-like receptors; TLRs). Remarkably, despite substantial variations in the nutrient content along the GI tract, we observed no clear correspondence between small molecule distribution and sensory gene expression in epithelial cells, fibroblasts, immune cells, or the ENS though we noted a substantial increase in TLR expression in the large intestine compared to the small intestine, mirroring regional differences in microbial density (**Figures 6D** and **S6A**). Moreover, while these classes of molecules are distributed differently across the different anatomical features of the gut, with the epithelial layer encountering the greatest diversity of small molecules, remarkably, we observed no clear spatial compartmental enrichment in the expression of different classes of GPCRs and PRRs (**Figure 6E**). Collectively, these observations suggest that the varying small molecule landscape within the gut is not reflected in similar broad spatial compartmentalization of the corresponding receptor genes.

### Differential receptor expression shapes the potential cell-type-specific response to microbial metabolites

The gut harbors a rich array of microbiota metabolites, including short-chain fatty acids, secondary bile acids, amines, and indole derivatives^1,2^, which act on diverse receptors such as GPCRs or nuclear receptors (NRs) to regulate host digestion, metabolism, immune system, and neural functions^1,2,62,63^. While the systemic effects of these metabolites are well-documented, the specific cell populations that respond to these small molecules are less well understood. To this end, we set our gut atlas in the context of the diverse array of microbiome-derived small molecules by constructing a custom microbiota metabolite—receptor database (MMRDB; STAR Methods). We constructed the MMRDB by combining GPCR—ligand (GLASS^64^) and NR— ligand (NRLiSt BDB^65^) databases with multiple microbiota-metabolite databases (MiMeDB^66^, gutMGene^67^), which we then supplemented with additional interactions curated from gutMGene^67^ (**Table S7**; STAR Methods).

We next integrated the MMRDB with our gut receptor atlas using the drug2cell^68^ pipeline (**Table S7**; STAR Methods). This integration revealed spatially distinct enrichment patterns, suggesting that microbial metabolite sensing is functionally compartmentalized, with specific metabolite classes preferentially detected in distinct gut niches. For instance, receptors for bile acid derivatives and SCFAs were consistently enriched in the mucosa, with SCFA receptors especially abundant in the colonic mucosa but also present in the myenteric and submucosal plexuses. In contrast, receptors for alkylamines and indoles were more broadly distributed. While such patterns were mostly regionally conserved, there were some exceptions. Notably, indole receptors were more highly expressed and shifted toward the lower mucosa in the distal colon (**Figure 6F**).

In terms of cell type specialization, receptors for microbiota metabolites were represented in all gut cell classes, with epithelial, ENS, interstitial, and immune cells, in particular, serving as major sensory hubs (**Figure S6B**). Interactions with microbiota metabolites were largely conserved across regions but highly cell-type-specific (**Figure 6G**, **S6C**, and **S6D**). For example, *Mrc1*+ macrophages selectively sensed alkyl polyamines like putrescine and spermidine, but this interaction was lost as they migrated to the top of the mucosa and differentiated into *Itgax*+ macrophages (**Figure 6G**), echoing recent findings that microbiota-derived polyamines regulate macrophage differentiation^69,70^. Succinic acid, a common microbial metabolite^62,63^, was sensed by tuft cells, L and N enteroendocrine cells, and their *Arx*+ progenitors (**Figure 6G**). In particular, the sensing of succinate by tuft cells is consistent with prior findings that succinate-mediated activation of *Sucnr1* in tuft cells triggers ILC2-driven immunity^71^. Since microbiota-derived succinate also improves glucose homeostasis^72,73^, its detection by glucagon-like peptide-1 (GLP1)–secreting L cells via *Sucnr1* suggests a potential metabolic mechanism, as GLP1 is an incretin that promotes insulin production^74^. Additionally, we uncovered underexplored interactions, such as deoxycholic acid signaling in colonic submucosal ICCs (**Figure 6G**), implicating a potential microbiome role in regulating pacemaker cell function. Overall, our atlas highlights specific cell types as mediators of gut–microbiota metabolite interactions and offers a rich hypothesis-generating resource for future mechanistic studies.

### The absence of the microbiome selectively remodels the gut on a structural, cellular and transcriptomic level

GF mice exhibit well-documented physiological differences from mice harboring a native microbiota—including reduced mucosal thickness, Paneth cell depletion, immune underdevelopment, metabolic alterations, and increased susceptibility to colitis^19,21,75^. However, a comprehensive, cell-type- and tissue-wide characterization of the differences in the GF remains lacking. Thus, to investigate the role of the microbiota in shaping the molecular, cellular, and spatial structure of the gut, we next analyzed the portion of our atlas derived from germ-free (GF) mice.

We first examined gut morphology across all four regions and found that, although the absence of the microbiome made villi thinner, more slender, and structurally fragile—a well-documented effect^76^, its impact on neighborhood composition was relatively modest. (**Figures 7A** and **7B**). As expected, our neighborhood analysis identified all major gut anatomical regions in GF mice, with region sizes—measured by the area associated with assigned cells—largely unchanged as compared to that observed in SPF mice. However, a few modest but statistically significant differences were observed: the mucosa was slightly reduced in the ileum and distal colon; the submucosa was reduced in the cecum and proximal colon; the muscularis propria was reduced in the cecum; and the myenteric plexus was also reduced in the cecum (**Figure 7B**). The cecum is known to have substantial structural changes in GF mice, where it is often several-fold larger than in mice harboring a microbiota^77^, consistent with our observation that, while subtle, many compartments of the cecum showed statistically significant differences from SPF controls (**Figure 7B**).

**Figure 7.**
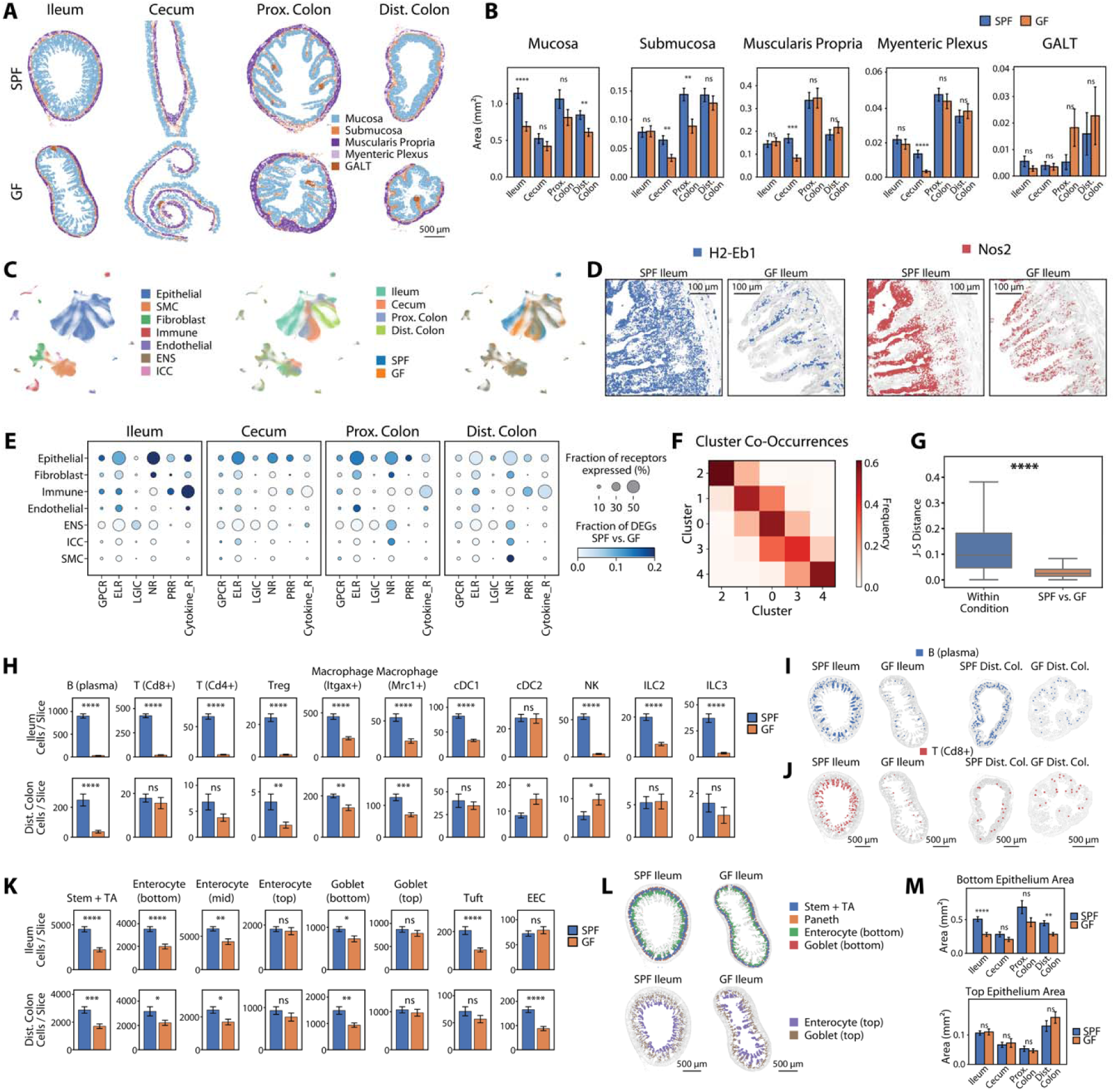
The absence of the gut microbiota alters cell-type abundance while having only modest effects on receptor expression. (A) Spatial distribution of cells in representative slices from each of the four regions in SPF or GF mice, colored by tissue neighborhood assignment. Scale bars: 500 µm. (B) Area of each neighborhood in SPF and GF conditions. Bars and error bars represent the mean and standard error across slices (slice numbers in **Figure S2A**). (C) UMAP representation of all cells imaged in all four regions of SPF and GF mice without Harmony batch correction, colored by major class (left), region (middle), or microbiome state (right). (D) Spatial distribution of RNAs of the specified genes (color) on top of all RNAs (gray) in a representative ileal slice of SPF or GF. Scale bar: 100 µm. (E) Expression of receptor categories across major cell classes between microbiome states in each gut region. Circle size represents the fraction of receptors in each category expressed by at least one cell type within each major cell class in the given region in either SPF or GF mice. Color indicates the fraction of these receptors that are differentially expressed between SPF and GF conditions in the given region (STAR Methods). (F) The frequency with which mucosal gene expression gradients clustered into one spatial pattern in a given region of the SPF mice is identified with another spatial pattern in the same region of the GF mice. (G) The distribution of Jensen-Shannon (J-S) distances between the spatial patterns for individual mucosal gene gradients within a region and microbiome state and between the same gene in the same region across different microbiome states (SPF vs. GF) (STAR Methods). Line: median. Box: 25% - 75% quantile. Whiskers: 0% - 95% quantile. **** indicates a p-value less than 10^-4^ as determined by a t-test. (H) Average number of select immune cells per slice in ileum or distal colon of the SPF or GF mice. Bars and error bars represent the mean and standard error across slices (slices numbers in **Figure S2A**). *, **, ***, and **** indicate p-values less than 0.05, 10^-2^, 10^-3^, and 10^-4^, respectively, while ns indicates a p-value greater than 0.05 as determined by a t-test. (I) Spatial abundance and distribution of plasma B cells (color) on top of all cells (gray) in ileum or distal colon of the SPF or GF mice. Scale bars: 500 µm. (J) As in (I) but for *Cd8*+ T cells. (K) As in (H) but for select epithelial cells. (L) As in (I) but for select cell types in top or bottom ileal mucosa. (M) Area of top and bottom epithelium in SPF and GF conditions. Bars and error bars represent the mean and standard error across slices (slice numbers in **Figure S2A**).

We next examined the role of the microbiota in modulating gene expression within cell types. Indeed, without batch correction, gene expression differences were readily visible in the UMAP representation, highlighting that the microbiota influences expression within our targeted gene panel (**Figure 7C**). We observed pronounced shifts in the expression of a small number of genes in a cell type- and location-specific fashion^78^. Notably, interferon-gamma stimulated genes such as MHC class II (*H2-Eb1*), its positive regulator *Ciita*, and nitric oxide synthase *Nos2* were strongly downregulated in the absence of the microbiome, while *Gpr108*, a negative regulator of TLR signaling, was upregulated (**Figures 7D**, **S7A**, and **S7B**).

However, beyond these more prominent differences, the overall transcriptional response to the absence of the microbiome was relatively modest. Specifically, the number of differentially expressed genes between microbiota states was smaller within individual cell classes than the number observed between gut regions in SPF mice (**Figures 6C** and **7E**; **Table S6**, STAR Methods). Among cell classes, epithelial cells exhibited the most extensive microbiota-dependent remodeling, although immune cells and fibroblasts also showed notable differences (**Figure 7E**). Interestingly, this microbiota-dependent remodeling varied by gut region, with ileum showing the greatest differences and the distal colon the least. Although the colon harbors higher microbiota densities than the ileum, the increased differences within the ileum may reflect a greater degree of microbe-host interactions due to the thinner mucus layer in the small intestine relative to that of the large intestine. In parallel, the absence of the microbiota had minimal impact on the spatial gene expression gradients. Major high-LMI populations—including enterocytes, goblet cells, FB1 fibroblasts, and telocytes—continued to display five distinct gradient types, with minimal switching of expression patterns between SPF and GF states (**Figure 7F**). Spatial gradients remained highly similar based on J-S distance (**Figure 7G**), and very few genes exhibited substantial spatial differences (**Figure S7C**). Notably, we observed ISG expression patches in all four gut regions in GF mice (**Figure S7D**), suggesting that, in addition to microbiota-dependent cues previously associated with these patches^59,79^, other non-microbial cues may also be involved. Together, these results indicate that the sensory capabilities of gut cells are largely host-intrinsic, and that changes in the abundance or metabolic activities of the microbiota are likely to induce only modest alterations in host cell function.

Despite modest changes in gene expression, we observed pronounced shifts in cell type composition in GF mice. Mucosal cell types, especially in the epithelial and immune compartments, were notably depleted (**Figure S7E**). In contrast, other cell classes—including fibroblasts, smooth muscle, the ICCs, and the enteric nervous system—showed no significant changes in cell abundance in the GF gut, suggesting their development is largely microbiota-independent (**Figure S7E**). Interestingly, remodeling of the mucosal immune compartment was both cell type-selective and regionally dependent. The ileal mucosa was most affected, with broad depletion of both adaptive and innate immune cells (**Figures 7H** and **S7F**), while the large intestine showed a more modest and cell type-selective pattern of immune depletion (**Figures 7H** and **S7F**). For instance, in the distal colon, plasma cells, macrophages, and Tregs decreased in abundance, while other T cell subsets and ILCs remained unchanged. In contrast, cDC2 and NK cell abundance increased (**Figure 7H**). These findings highlight the strong regional specificity of the immune response to the microbiota and the distinct remodeling that occurs in its absence.

In addition to the changes in the immune compartment, we found that the developmental process of the gut epithelium is also modulated by the microbiome. In both ileum and distal colon, the abundance of bottom epithelial cells, including Paneth in ileum and stem and bottom enterocytes and goblet cells in both regions, decreased in number per cross-section, whereas the abundance of top or mature epithelial cells, including top enterocytes and goblet cells, remained unchanged (**Figures 7K**, **S7G**, and **S7H**). This effect was present but less significant in cecum and proximal colon (**Figures 7K**, **S7G**, and **S7H**). The decrease in Paneth cell numbers confirmed previous reports^26^, whereas our atlas revealed a systemic depletion across all bottom or immature epithelial cell populations. Indeed, we observed that the band of bottom epithelial cells visually contracted in depth in the ileum and distal colon in GF compared to SPF state, while the same trend did not hold true for the band of top epithelial cells (**Figures 7L** and **S7I**). This observation was further validated by comparing the area per cross-section for bottom versus top epithelial zones (**Figure 7M**). Furthermore, by comparing the progression of actual mucosal length with positions along the mucosal pseudospace axis, we observed a slower growth in real length at lower pseudospace positions, indicating a relative shortening of the lower mucosa in the absence of the microbiome (**Figure S7J**). Likewise, cell abundance distributions along the mucosal pseudospace axis showed a relative downward shift of epithelial populations, including enterocytes and goblet cells, consistent with the shortening of the lower mucosa (**Figure S7K**). These findings collectively suggest that gut epithelial development is modulated by the microbiome, and that its absence may accelerate epithelial development, resulting in the depletion of immature and a relative enrichment of mature cell populations.

## DISCUSSION

The gut contains a diverse array of cell types, each tuned to sense distinct host-, diet-, and microbe-derived molecules. These sensory roles vary across regions and are shaped by local spatial heterogeneities. The microbiome can further reshape this molecular landscape, influencing the cellular and spatial organization of the gut. To systematically explore these features, we used MERFISH to generate a spatial transcriptomic atlas of the expression profile of most sensory-related genes in over 2 million cells across four regions of the mouse lower digestive tract, under both SPF and GF conditions. Through a combination of improved background filtering and segmentation, we defined the expected cell types across all layers of the gut, including rare populations, and defined previously unappreciated divisions within several populations. By providing a high-cellular- and -spatial-resolution view of multiple regions of the gut in two microbiome states, we envision that this atlas may prove to be a useful resource for a deeper understanding of the spatial organization of gut sensation in a range of questions.

Several observations illustrate this point. For instance, by combining detection sensitivity with cellular throughput, our MERFISH-based atlas revealed a population of mature enterocytes predominantly in the distal colon marked by the expression of the epithelial sodium channel subunit *Scnn1g*. To our knowledge, this population has not been previously identified in mice. Interestingly, this population expresses multiple genes in common with a human mature enterocyte population, which is marked by the calcium-dependent chloride channel BEST4 and similarly localized predominantly in the distal colon top mucosa. Multiple functions have been proposed for the BEST4+ enterocytes, including ion transport, mucus hydration, and regulation of the luminal pH^54^. However, as there has been no previously defined homolog in a convenient model organism, it has been difficult to validate the hypothesized functions of these cells^54^. Like the BEST4+ enterocytes, the *Scnn1g+* enterocytes may play a similar role in ion transport and mucus hydration given its expression of all three subunits of the non-voltage-gated epithelial sodium channel. While the lack of a functional Best4 and some BEST4+ enterocyte markers may indicate potential functional distinctions between these populations, genetic manipulation of this mouse population may prove useful in the exploration of BEST4+ enterocyte functions.

Another major insight provided by our atlas is that there is an extensive degree of fine-tuning of sensory gene expression and, thus, potentially sensory function within most cell types across multiple spatial scales within the gut. At broad spatial scales, we observed that while the overall cellular identities are conserved across different regions of the gastrointestinal tract, most cell types exhibit region-specific receptor expression, reflecting a potential tuning of their sensation to the differences in the small molecule environment within different gut regions. As the cells in direct contact with the lumen, it is perhaps not surprising that epithelial cells showed the greatest regional specialization, with over half of their receptors varying by region, particularly in receptors for lipids, peptides, amines, and microbial cues. However, these patterns showed little correlation with known variations in small molecules along the GI tract, suggesting that specialization is not solely shaped by local ligand availability.

At finer spatial scales, prior studies had described mucosal bottom-to-top gene expression gradients in epithelial and some fibroblast populations^12,15–18,57^. Our atlas now demonstrates that such spatial fine-tuning is widespread across most mucosal cell types, including stromal and immune populations, with the most pronounced gradients seen in epithelial and fibroblast subsets, and the least in immune cells. These gradients appear largely conserved across gut regions and microbiome conditions, suggesting they are intrinsic features of each cell type. Importantly, these gradients suggest that most cells adapt their gene expression to micron-scale variations within the mucosa. Importantly, our study offers a more precise framework for interpreting cellular heterogeneity in the gut. Subsets previously thought to represent distinct cell types—such as top and bottom goblet cells, telocyte subtypes, or other fibroblast subpopulations (FB1s)—are more accurately described as positions along spatially continuous gene expression gradients. This refined view provides a unifying perspective on gut cellular diversity and highlights the power of spatial transcriptomics to resolve long-standing ambiguities in tissue organization.

In addition to spatially cued gene expressions, cells in the intestinal mucosa are intricately organized, but this cellular-scale architecture has been difficult to define comprehensively across regions and cell types. Leveraging the conserved epithelial maturation process as a spatial reference, we applied a pseudospace analysis to align mucosal organization across the gut. This revealed broadly conserved spatial patterns in epithelial and fibroblast populations, while immune cell organization varied more substantially. Notably, the cecum stood out with distinct immune cell localization at the base of the crypt, suggesting region-specific roles in microbial interaction. By precisely mapping gene expression gradients and cell localization, we also developed a spatial prioritization analysis for ligand–receptor interactions, linking chemokine expression levels to immune cell distributions in the mucosa. As image-based single-cell transcriptomics become more widely adopted, such analysis approaches can offer powerful frameworks for dissecting cellular organization and interaction within intact tissues.

The microbiota has broad effects on host physiology, influencing processes such as immune development, epithelial differentiation, and disease susceptibility^19,21,75^. To systematically map these effects, our atlas included a germ-free (GF) component, enabling molecular, cellular, and tissue-scale comparisons with specific-pathogen-free (SPF) controls. We found that most features of gut organization are host intrinsic: cell types were largely conserved, and both receptor expression profiles and spatial gene expression patterns were mostly unchanged in the absence of the microbiome. The most pronounced changes were observed in immune cell abundance. The ileum showed widespread depletion of both adaptive and innate immune cells, while the large intestine exhibited more selective shifts—such as reduced plasma cells, Tregs, and macrophages in the distal colon, alongside increased cDC2 and NK cells. Epithelial development was also affected, with a depletion of immature populations in the gut epithelium of GF mice. These findings highlight the critical yet selective role the microbiome plays in shaping immune composition and epithelial dynamics. Taken together, these measurements reveal a rich spatial and regional organization of the mammalian gut and provide a valuable foundation for future mechanistic studies of host–microbe interactions and gut physiology.

## Limitations of the study

As image-based transcriptomic methods are inherently targeted, our conclusions are necessarily limited to the genes within our selected panel, which is enriched for genes associated with sensory functions. Therefore, our findings regarding the host-intrinsic nature of receptor expression in the absence of the microbiome may not extend to other cellular processes. Moreover, we restricted our analysis to male C57BL/6 mice fed a standard diet (NIH-31M); therefore, the conclusions we draw from our atlas will not reflect sex, strain, or dietary differences, all of which can be important modulators of the sensory organization of the gut and its interaction with the microbiome. Additionally, because this is a transcriptomic study, our functional interpretations are inferred from the presence of transcripts rather than direct measurements of protein expression or cellular function. Finally, our spatial expression gradient analysis focused on radial and circumferential organization of the mucosa, reflecting the orientation of the gut slices collected. However, anatomical transitions between gut regions— such as the proximal to distal colon—may not be discrete. Thus, while our atlas revealed regional differences, it does not capture the underlying longitudinal gradients that may underlie more continuous variation between regions. Future studies that capture the full longitudinal axis using techniques such as Swiss rolling will be important for providing a more continuous and comprehensive view of regional specialization.

## ACKNOWLEDGMENTS

We thank Brianna R. Watson for assistance in microscope construction and maintenance, Richard Shim and Josh J. Luce for assistance for reagent preparation, Evan Yang for assistance with bulk RNA sequencing, Gokul Gowri and Allon Klein for suggesting and proof-of-concept work with latent mutual information, and members of the Moffitt laboratory and members of the Harvard Medical School Rodent Histopathology core for their support and helpful comments. This work was supported by grants to J.R.M. from the Chan Zuckerberg Initiative, the Helmsley Charitable Trust, and the NIH (R01GM143277 and P30DK034854); to P.C. from the Charles A. King Trust Postdoctoral Research Fellowship Program, Bank of America, N.A., Co-Trustees; and to U.S.H. from a postdoctoral mobility fellowship from the Swiss National Science Foundation and the Novartis Foundation for Medical-Biological Research. Portions of this research were conducted on the O2 High Performance Computer Cluster, supported by the Research Computing Group at Harvard Medical School. Some figure items were created with BioRender.

## AUTHOR CONTRIBUTIONS

Conceptualization, R.J.X. and J.R.M.

Methodology, R.J.X., P.C., P.B.N., U.S.H., R.A.I., and J.R.M.

Software, R.J.X., T.L., L.G., and J.R.M.

Investigation, R.J.X.

Formal analysis, R.J.X. and P.B.N.

Data curation, R.J.X.

Resources, L.G., R.A.I., and J.R.M.

Visualization, R.J.X., T.L., and L.G.

Writing – original draft, R.J.X. and J.R.M.

Writing – review & editing, R.J.X., P.C., P.B.N., U.S.H., T.L., L.G., R.A.I., and J.R.M.

Supervision, J.R.M.

Funding acquisitions, J.R.M.

## DECLARATION OF INTERESTS

J.R.M. is a co-founder of, stakeholder in, and advisor for Vizgen, Inc. J.R.M. is an inventor on patents associated with MERFISH applied for on his behalf by Harvard University and Boston Children’s Hospital. J.R.M.’s interests were reviewed and are managed by Boston Children’s Hospital in accordance with their conflict-of-interest policies. P.C. is an inventor on patents associated with MERFISH applied for on his behalf by Boston Children’s Hospital.

## RESOURCE AVAILABILITY

### Lead contact

Further information and requests for resources and reagents should be directed to and will be fulfilled by the Lead Contact, Jeffrey Moffitt.

### Materials availability

This study did not generate unique reagents.

### Data and code availability

All MERFISH measurements are available on Dryad (DOI listed in the key resources table). Bulk RNA-seq data generated in this paper are available on GEO (accession number listed in the key resources table). The accession numbers for all publicly available scRNA-seq data used in the analysis in this paper are in the key resources table. A web-based viewer of this atlas is available at moffittlab.github.io/visualization/2025_XuMurineGIAtlas/.

Any additional information required to reanalyze the data reported in this work is available from the lead contact upon request.

## STAR⍰METHODS

### KEY RESOURCES TABLE

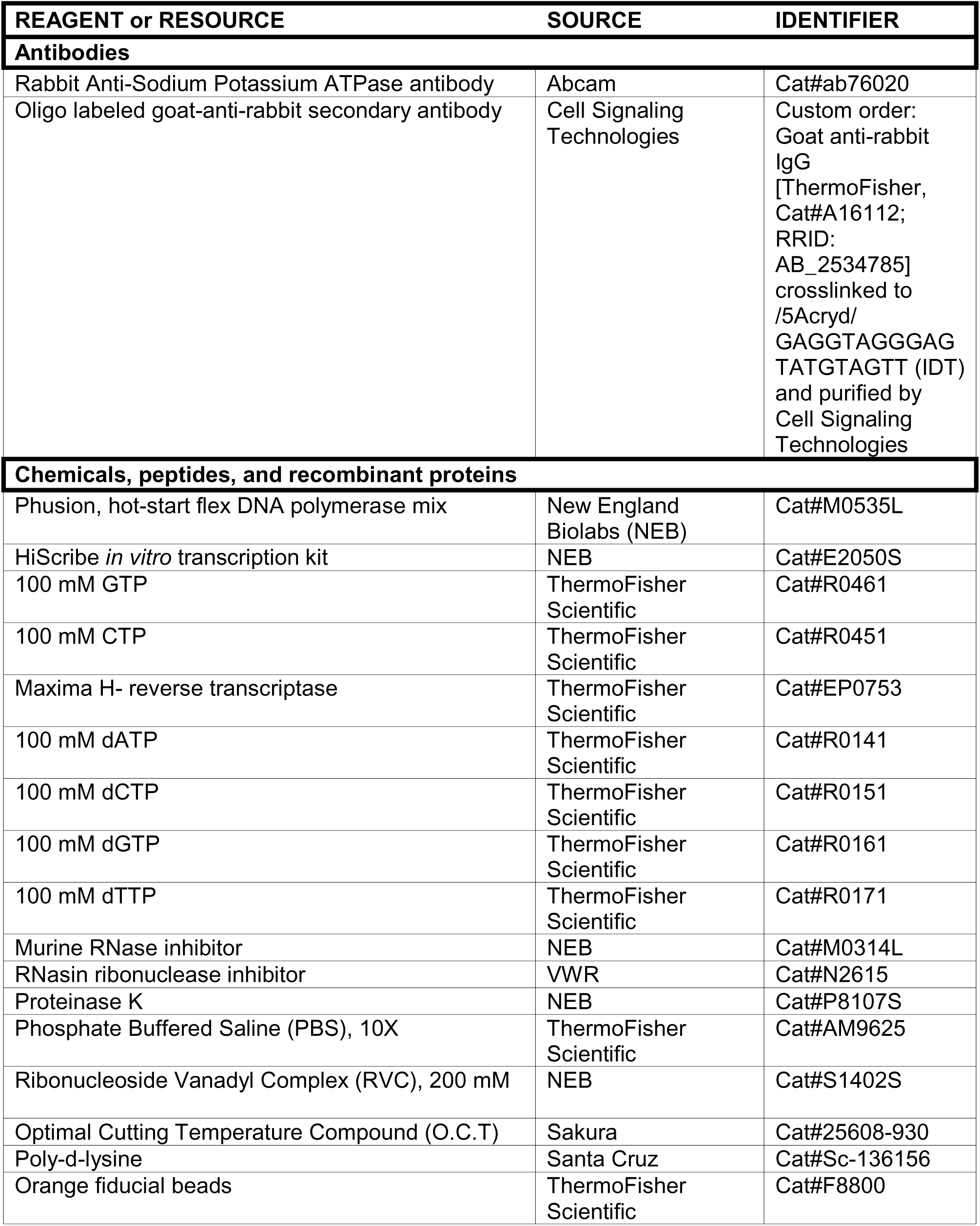
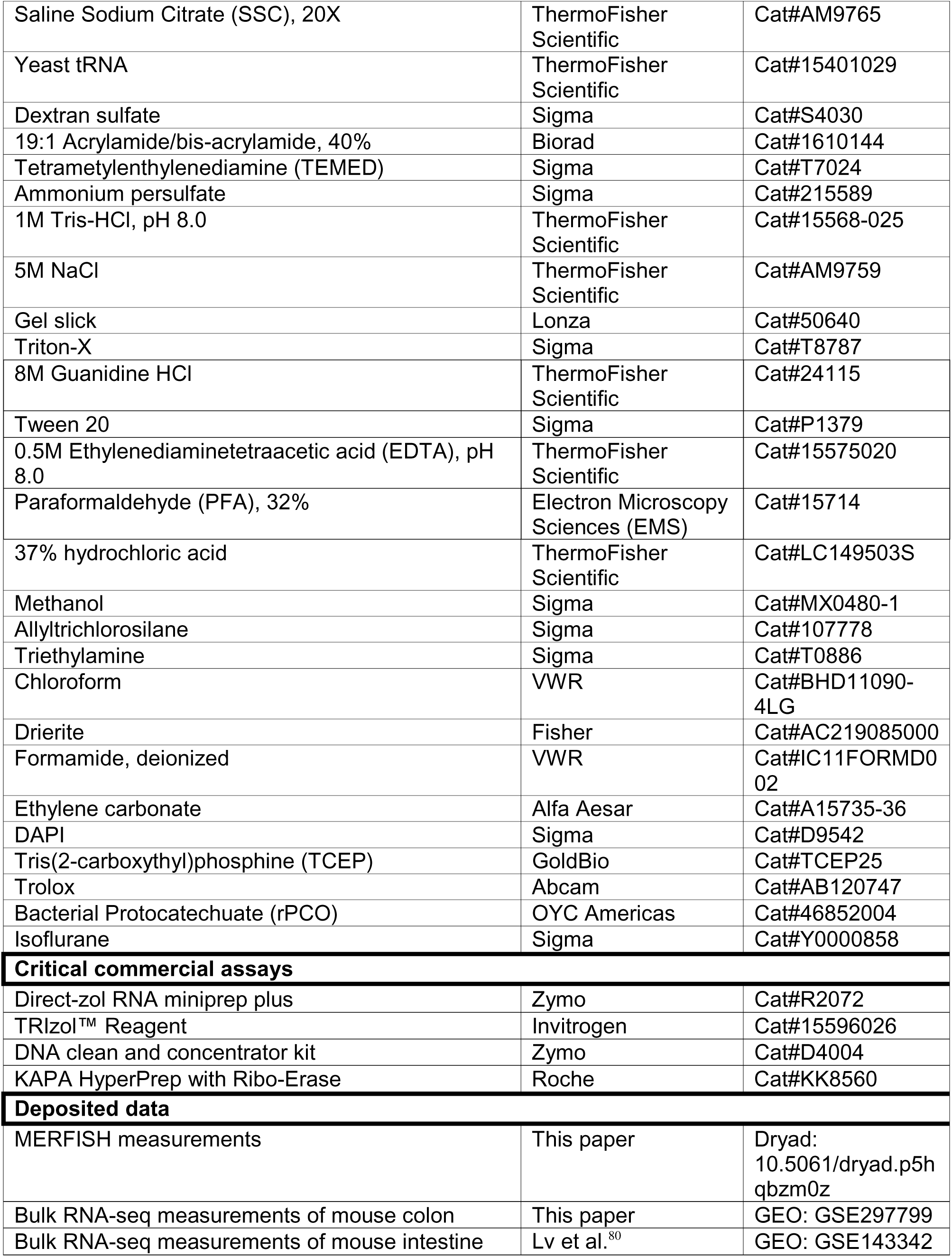
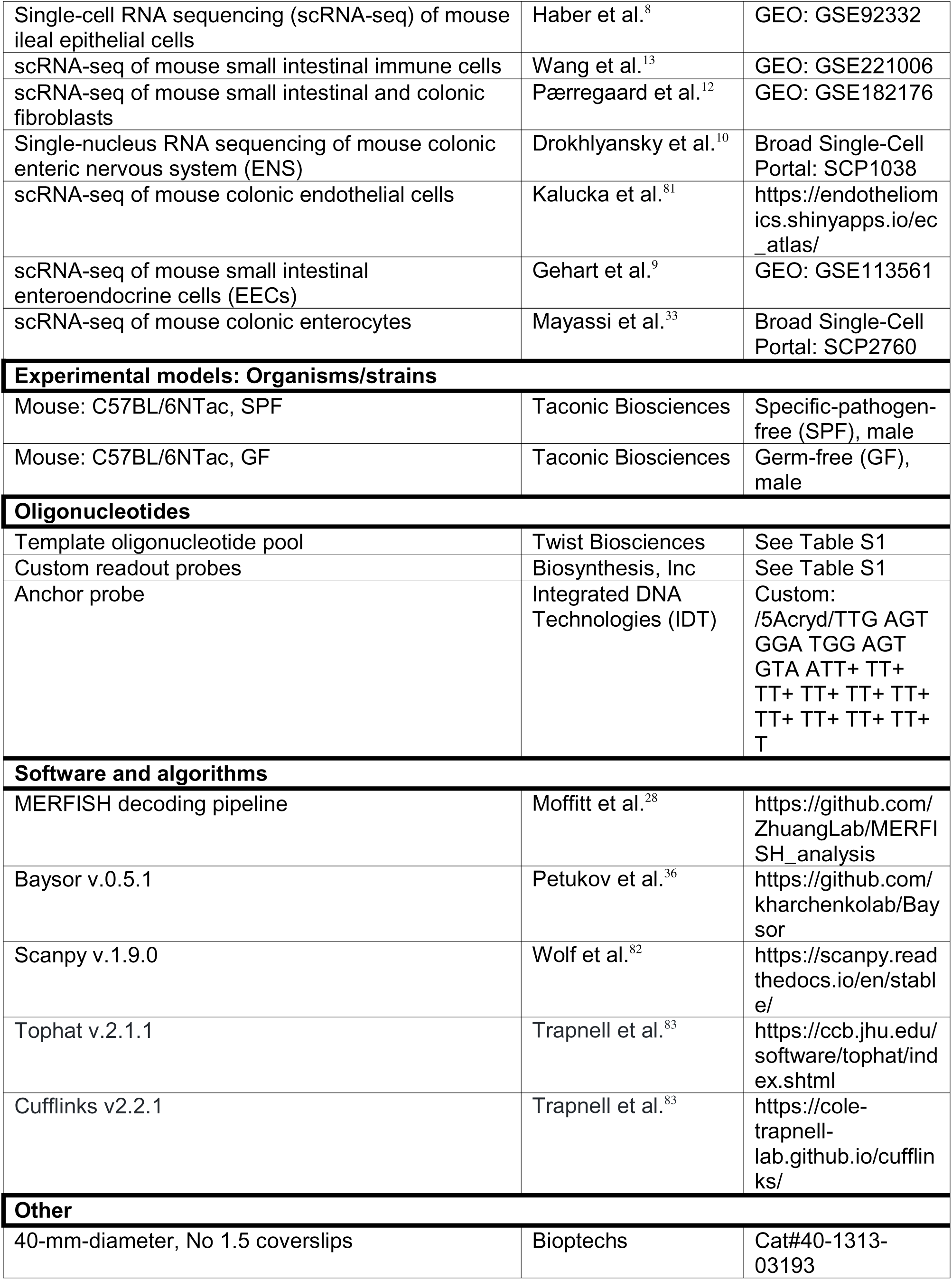
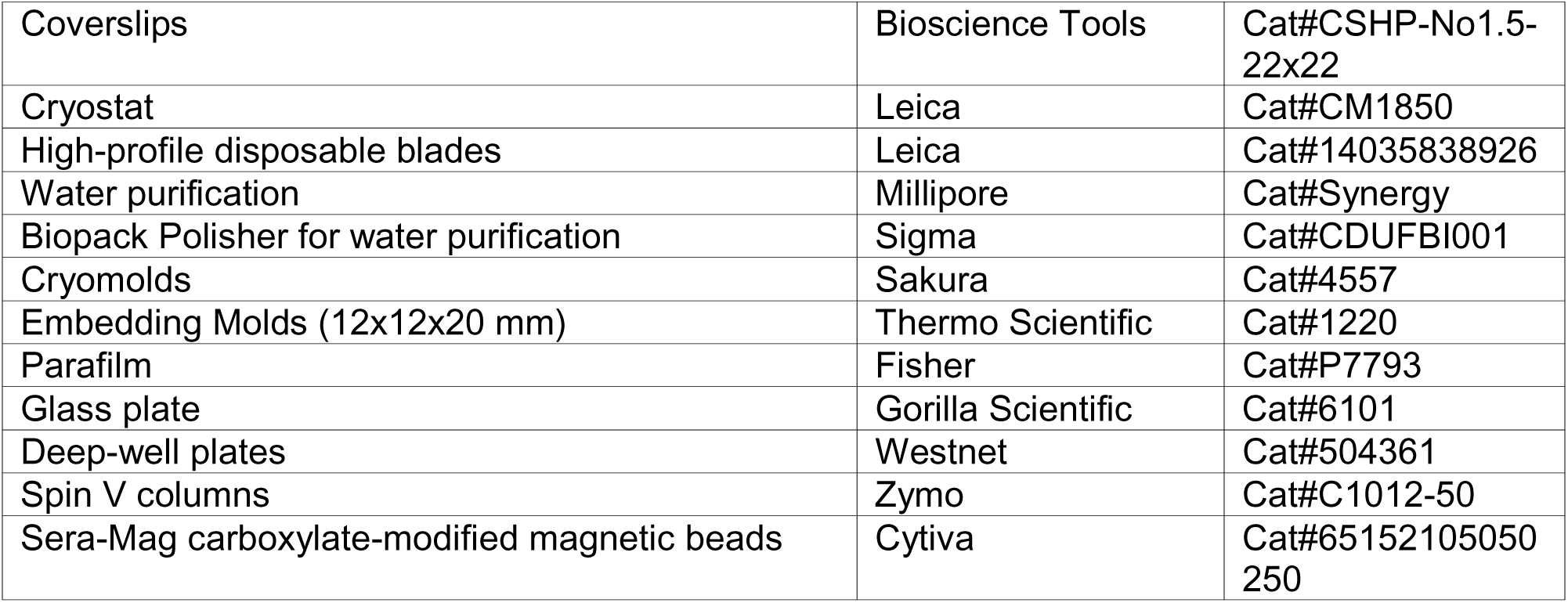

### RESOURCE AVAILABILITY

#### Materials availability

This study did not generate new materials.

#### Data and code availability

All MERFISH measurements are deposited on Dryad (DOI listed in the key resources table). Bulk RNAseq data generated here are deposited on GEO (accession number listed in the key resources table). This paper also analyzed publicly available bulk and scRNA-seq data (accession numbers listed in the key resources table).

### EXPERIMENTAL MODEL AND STUDY PARTICIPANT DETAILS

#### Animals

C57BL/6NTac mice were purchased from Taconic Biosciences as either specific pathogen free (SPF) or germ free (GF). The SPF mice were housed at the Harvard Center for Comparative Medicine for 3 or more days after receipt and prior to sacrifice while the GF mice were sacrificed upon receipt. Adult male mice aged between 8-12 weeks were used. All animals were on a 12-hour-light/12-hour-dark cycle with *ad libitum* access to the NIH-31M diet and water. All animal experimentation was performed in compliance with NIH guidelines and was performed with protocols approved by the Harvard Medical School Institutional Animal Care and Use Committee (IACUC) under protocol number IS00003215.

### METHOD DETAILS

#### Bulk RNA sequencing

As a validation of MERFISH measurements, we performed bulk RNA sequencing of the mouse intestine harvested from SPF mice. Specifically, distal colon tissue was harvested from C57BL/6NTac mice, flushed and rinsed with 4 mM ribonucleoside vanadyl complex (RVC; NEB S1402S) in 1× phosphate-buffered saline (PBS; Thermo AM9625) at 4 °C. The tissue was then dissolved in TRIzol (Invitrogen 15596026), and RNA was extracted using the Direct-zol RNA Miniprep Plus kit (Zymo R2072) following the manufacturer’s instructions. Total RNA was then depleted for ribosomal RNA using the HyperPrep with Ribo-Erase kit (Roche KK8560), and paired-end sequencing of 75 bases was performed on an Illumina MiSeq Instrument. RNA abundance was determined using tophat (v2.1.1) and cufflinks (v2.2.1) with default parameters, aligning against the mouse genome (GRCm39). Raw and analyzed sequencing data were deposited in the gene expression omnibus (GEO) at accession GSE297799.

#### MERFISH library design and construction

To profile gene expression in the mouse intestine, we selected a panel of 1,922 genes. The first component of this panel contained ∼200 well-established cell-type markers for major cell populations present in gut tissue (e.g., epithelial cells, fibroblasts, immune cells, endothelial cells, smooth muscle cells, cells of the enteric nervous system, interstitial cells of Cajal (ICC], and adipose cells) as well as established subtypes of these cells (e.g., B cells, T cells, etc.). The second component of this panel comprised a near comprehensive set of mouse receptors. To build a receptor list, we mined the KEGG database^84^ to compile all genes from the following categories: G-protein-coupled receptors (GPCRs; mmu4030), ligand-gated ion channels (LGICs; mmu04040), enzyme-linked receptors (ELRs, including receptor tyrosine kinases, receptor tyrosine phosphatases, receptor serine/threonine kinases, and receptor serine/threonine kinase; mmu01001 and mmu01009), nuclear receptors (NRs; mmu03310), pattern recognition receptors (PRRs, including Toll-like receptors, C-type lectin receptors, NOD-like receptors, and RIG-I-like receptors; mmu04054), complement receptors (mmu04610), cytokine receptors (mmu04050), Wnt receptors (mmu04310), Hedgehog receptors (mmu04340), Notch receptors (mmu04330), bone morphogen protein receptors (BMP; mmu04050), and Hippo receptors (mmu04392). In addition, we included all known host-derived ligands for these pathways, including cytokines (mmu04052) and ligands for the Wnt, Hedgehog, Notch, BMP, and Hippo pathways. With a few exceptions, we did not include intracellular signaling components within any of these pathways. While we aimed to be comprehensive in all receptor categories, there was one category for which we made an exception: olfactory, taste, and vomeronasal receptors. While there is ectopic expression of olfactory receptors^85^, we selected only receptors from these categories with evidence of expression from bulk RNA sequencing data (GSE143342), keeping only 791 of 1,527 genes in these categories.

Imaging-based approaches to single-cell transcriptomics can be sensitive to RNAs that are very highly expressed, as increased crowding of single-molecule fluorescence signals can corrupt the optical barcodes used to identify RNAs. In order to reduce signal crowding and improve detection sensitivity, we removed a small subset of highly expressed receptors in published bulk RNA-sequencing (GSE143342)^80^. Specifically, any RNA with an FPKM greater than 50 was removed. In addition, MERFISH obtains high quality signals by targeting each RNA with multiple probes that bind to different regions of the same RNA. Therefore, we also screened out RNAs for which the expressed isoforms are too short to support sufficient numbers of MERFISH probes. Collectively, these cuts resulted in the removal of less than 3 % of the receptors and ligands from the categories above.

MERFISH encoding probes targeting the above genes were designed for the mouse genome annotation (GRCm39) using a previously described pipeline^27,28^, with the following parameters: 30-nt target region length, GC fraction between 0.4 and 0.6, a predicted melting temperature between 65 and 75 °C, on-target binding specificity between 0.75 and 1, and 75 target regions per gene. About 10 % of the targeted genes were too short to support 75 target regions. For these genes, we designed as many regions as possible, excluding genes only if fewer than 30 target regions were possible.

To encode these genes, we used a 39-bit long, Hamming distance 4, and Hamming weight 4 barcode set that contained 2,070 barcodes, out of which 1,922 were assigned to the selected markers and receptors. The remaining barcodes served as “blank” controls, providing a direct measure of the false positive rate in each measurement. Individual bits were associated with a previously designed set of readout probes^37,86^ which were first pre-screened against the mouse tissue to identify and remove probes that had modest but measurable degrees of non-specific binding. Encoding probe template molecules were then designed by assembling (1) the reverse complement of three of the four readout probes associated with the barcode assigned to each RNA, (2) a forward primer, (3) the T7 promoter, and (4) one of the desired target regions. The three readouts associated with each target region were selected from the four appropriate readouts at random but such that there was roughly even usage of each readout among the target regions associated with a given RNA. The sequences of probe templates, as well as barcode assignment to individual genes, are provided in **Table S1**.

In addition to the 1,922 genes described above, the third component of our MERFISH panel were 12 highly expressed genes—*Chga*, *Sst*, *Pyy*, *Gcg*, *Sct*, *Nts*, *Cck*, *Ghrl*, *Gip*, *Muc2*, *Vil1*, and *Mptx2*. As we anticipated that these genes could be expressed at very high levels within the cells that express them, we encoded them with sequential barcodes and assigned each gene to a unique readout. Encoding probes for these genes were designed as above with one readout per encoding probe (**Table S1**).

The templates were ordered as an oligo pool (Twist Biosciences) and amplified following a previously described protocol^37^. Briefly, the oligo pool was amplified using limited cycle PCR (NEB M0535L) and purified via spin column (Zymo Spin V with DNA Clean & Concentrator-5, C1012-50 and D4004). The product was amplified and converted to RNA with a high yield *in vitro* transcription (NEB E2050S) and purified with home-made solid-phase reversible immobilization beads^37^ (based on Cytiva 65152105050250). The RNA products were then converted back to DNA via reverse transcription (Thermo EP0753), followed by alkaline hydrolysis to remove the RNA template and purification of DNA products by SPRI beads. Finally, the encoded probe library was adjusted to the desired concertation via ethanol precipitation.

#### Tissue collection and sectioning

Fresh frozen mouse intestinal tissue collection and cryosection were performed following a previously described protocol^37^. Briefly, mice were euthanized by isoflurane (Sigma Y0000858) treatment and cervical dislocation. Intestinal tissues were extracted, rinsed and flushed in pre-chilled 4 mM RVC in 1× PBS on ice, followed by a 1-hour incubation in the same solution at 4 °C. The tissue was then embedded in Optimal Cutting Temperature Compound (OCT; Sakura 25608-930) at 4 °C, flash frozen on dry ice, and stored at - 80 °C until cryosection.

Coverslips (Bioptechs 40-1313-03193) were silanized (Sigma 107778) and coated with poly-d-lysine and orange fluorescent beads (Santa Cruz sc-136156; Invitrogen F8800) as previously described^37^, and 10-μm tissue sections were collected on a cryostat (Leica CM1850) set to -23 °C. To improve tissue adherence, slices were briefly melted onto coverslips with fingertip heat, immediately refrozen in the cryostat chamber, and then left to dry at -23 °C inside the cryostat chamber for 2-4 hours via sublimation. After drying, the slices were fixed with 4% v/v paraformaldehyde (PFA; Electron Microscopy Sciences 15714) in 1× PBS at 4 °C for 20 minutes, and rinsed twice with 4 mM RVC in 1× PBS for 5 minutes each at 4 °C. To permeabilize samples, coverslips were submerged in 70% ethanol and stored at 4 °C in 60 mm Petri dishes sealed with parafilm for at least 12 hours. Samples were stored in this condition for no more than 1 week before proceeding with subsequent steps. RNAse-free water generated via reverse osmosis (Synergy UV, Millipore) with a Biopack polisher (Sigma CDUFBI001) was used for all steps.

#### MERFISH sample preparation

Mouse intestinal tissue slices were prepared for MERFISH imaging following a previously described protocol^37^. Briefly, an encoding probe hybridization solution (30% v/v formamide [VWR IC11FORMD002], 1 mg/mL yeast tRNA [Thermo 15401029], and 10% w/v dextran sulfate [Sigma S4030] in 2× saline-sodium citrate [SSC; Thermo AM9765]) was supplemented with 50 μM total MERFISH probe library, 2 μM anchor probe (/5Acryd/TTG AGT GGA TGG AGT GTA ATT+ TT+ TT+ TT+ TT+ TT+ TT+ TT+ TT+ TT+ T where /5Acryd/ represents an acrydite modification and T+ indicates locked nucleic acid), and 4 μg/mL Na^+^/K^+^-ATPase primary antibody (Abcam ab76020). Coverslips were removed from the 70% ethanol and rinsed twice with 30% v/v formamide in 2× SSC for 5 minutes each at room temperature. Coverslips were then placed sample-side down onto a droplet of 100 μL of the encoding probe solution on a parafilm-lined 150 mm Petri dish. A damp Kimwipe was added to the petri dish, which was then sealed, to create a humidified environment to prevent the evaporation of the hybridization solution. The staining chamber was then incubated overnight for 12-24 hours at 37 °C in a humidified incubator.

Following the first overnight incubation, the coverslip was removed from the chamber, rinsed twice with 30% v/v formamide in 2× SSC, and then stained using the same protocol as above but with 100 μL of fresh encoding probe hybridization solution in which the primary antibody was replaced with 4 μg/mL of an oligonucleotide-labeled goat-anti-rabbit secondary antibody. The coverslip was then incubated at 37 °C in a humidified incubator for an additional 24-36 hours for a total incubation time of 36-48 hours. Coverslips were carefully peeled off the parafilm with tweezers and rinsed twice in 2× SSC at room temperature before proceeding immediately to subsequent steps.

To decrease background, samples were embedded in a thin polyacrylamide file and cleared, as described previously^37^. Briefly, samples were rinsed in a hydrogel solution comprised of 4% 19:1 acrylamide/bis-acrylamide (Bio-Rad 1610144), 300 mM NaCl (Thermo AM9759), 0.03% w/v ammonium persulfate (Sigma 215589), and 0.15% v/v tetramethylethylenediamine (TEMED; Sigma T7024) in 50 mM Tris-HCl (Thermo 15568-025). A glass plate (Gorilla Scientific 6101) was cleaned and coated with Gel-Slick (Lonza 50640) following the manufacturer’s instructions. The coverslip was then inverted, sample-side-down, onto a 100 μL droplet of this gel solution on the glass plate, taking care to avoid trapping air bubbles in this thin layer of gel solution. The gel then polymerized for 2 hours at room temperature, after which the coverslip with the attached gel was carefully removed from the glass plate with a razor blade.

Samples were then incubated in a digestion buffer comprised of 800 mM guanidine HCl (Thermo 24115), 30 mM EDTA (Thermo AM9260G), 0.25% v/v Triton-X (Sigma T8787), 5% v/v Tween 20 (Sigma P1379), and 1:100 proteinase K (NEB P8107S) in 50 mM Tris-HCl at 37 °C for 12-24 hours. This digestion buffer was then replaced with freshly made digestion buffer and the digestion was repeated a second time. Following digestion, coverslips were washed four times in 2× SSC at room temperature for 5 minutes each and were stored in 2× SSC at 4 °C for up to two weeks prior to MERFISH imaging.

#### MERFISH imaging

MERFISH data collection was performed on a home-built microscope and fluidics system as previously described^32,36^. Briefly, a Nikon Ti2 microscopy body was equipped with a 60× CFI PlanApo lambda oil objective (Nikon), an objective nanopositioner (Nano F200 Mad City Labs), a Celesta light engine (Lumencor), and two CMOS cameras (Hamamatsu ORCA Flash 4.0) coupled into the microscope body via a two-camera splitter (Cairn TwinCam). The focus was maintained with a custom autofocus system. The sample was mounted in a flow chamber (Bioptechs FCS2), and fluid flow was controlled with a peristaltic pump (Gilson MP1) coupled to computer-controlled valves (Hamilton MVP and HVXM 8-5). The microscope and flow system were controlled by home-built control software (https://github.com/ZhuangLab/storm-control).

A set of custom cartridges were assembled with the following buffers required for the automated MERFISH measurement as described previously^37^. Readout probes were hybridized in a readout hybridization buffer comprised of 10% v/v ethylene carbonate (Alfa Aesar A15735-36) and 0.25% v/v Triton-X (Sigma T8787) in 2× SSC with 3 nM each of two different readout probes (**Table S1**). Excess readouts were removed in a readout wash buffer identical to the readout hybridization buffer but without readout probes. An imaging buffer comprising 4 μM Trolox-quinone, 0.5 mg/mL Trolox (Abcam AB120747), 1:500 recombinant protocatechuate 3,4-dioxygenase (rPCO; OYC Americas 46852004) and 5 mM protocatechuic acid (Sigma 37580-25G-F) in 2× SSC was used to reduce photobleaching and improve dye brightness. Fluorescent dyes were removed from readout probes with a cleavage buffer composed of 50 mM tris(2-carboxyethyl)phosphine (GoldBio TCEP25) in 2× SSC.

To prepare a sample for MERFISH imaging, the coverslip was stained with 5 mL of readout hybridization buffer containing the first pair of readout probes for 15 minutes at room temperature, followed by a 10-minute wash in 5 mL readout wash buffer supplemented with 1 mg/mL DAPI (FisherScientific D1306). Samples were briefly stored in 2× SSC prior to imaging. To perform a MERFISH measurement, a coverslip was mounted in the flow chamber and imaging buffer wash flown across the sample. 9 z-planes at 1.5 μm spacing were imaged for each field-of-view (FOV) covering a distance range of 2.5 μm to 16 μm from the coverslip surface with illumination in both the 635-nm and 750-nm channels to measure readout probes. A single z-plane was imaged with the 535-nm channel to identify the location of fiducial beads on the coverslip surface, and in the first imaging round, the same 9 z-planes were also imaged with the 405-nm and 473-nm channels to identify nuclei via the DAPI stain and the Na^+^/K^+^-ATPase membrane immunofluorescence signals. At the end of each round of imaging, the sample was incubated in cleavage buffer for 15 minutes to remove fluorescent signals from the previous round of readout probes, the sample was washed with 2× SSC for 3 minutes to remove excess TCEP, the sample incubated in the next readout hybridization mixture for 15 minutes, and then excess readout probes removed with a 5 minute incubation in readout wash buffer. Imaging buffer was then flowed across the sample and the next round of imaging was performed.

### QUANTIFICATION AND STATISTICAL ANALYSIS

#### RNA identification and quality score filtering

To identify RNAs within the MERFISH data, raw image stacks were decoded with a previously described MATLAB pipeline on the Harvard Medical School Orchestra2 cluster^37^. Briefly, this pipeline (github.com/ZhuangLab/MERFISH_analysis) identifies RNA molecules with a pixel-based, soft-decoding approach in which the normalized intensity vector of each pixel across imaging rounds is matched to that expected for each MERFISH barcode via Euclidean distance. Directly adjacent pixels that are matched to the same barcode are aggregated to form one tentative RNA molecule. This decoding process generates a set of metadata, including Euclidean distance to matched barcode (“distance”), number of pixels that form one RNA molecule (“area”), as well as the average of the Euclidean norm of the normalized signal intensity across all imaging rounds (“brightness”).

Included in these identified RNAs were a population of false positives. Previous efforts discriminated these false molecules from real molecules by thresholding on all or some of these metadata properties^37^. However, these approaches required a degree of dataset-by-dataset adjustment of metadata thresholds, in addition to leaving behind a low but potentially meaningful level of false positive, as judged by the frequency with which blank barcodes were detected. As an improvement, we developed a new approach to discriminate real from false MERFISH signals. Specifically, we used these RNA metadata properties to define a single quality score for each RNA and then applied a single quality score threshold across all data to exclude false positives.

To compute the quality score, we computed two 3D histograms of the area, brightness, and distance metadata quantities for each individual MERFISH dataset—one for all barcodes (putative real molecules combined with false positives) and the other for blank barcodes only (known false positives). In the null model that all detected RNAs with MERFISH are false positives, the fraction of blank barcodes relative to all barcodes for each bin in these histograms would be equal to the fraction of barcodes in our library that are blank barcodes. Therefore, a smaller fraction of blanks observed for any given range of metadata properties would suggest that RNAs with those metadata properties are proportionally more likely to represent true positive measurements. The use of this 3D histogram to define complex metadata cuts has been proposed previously^87^; however, here we used the ratio of these two histograms to define a quality score for each set of metadata properties. First, we normalized the ratio of the blank counts to the total counts for each RNA metadata bin by the fraction of barcodes in the library assigned to blanks. Second, we defined the quality score as one minus this quantity.

To estimate the quality score for each RNA, we computed the histograms above with area binned into bins of size 1, the logarithm of the brightness binned into bins of size 0.1, and the distance binned into bins of size 0.02. We then computed the quality score for each of these bins as described above. Finally, we assigned a unique quality score to each RNA based on a nearest neighbor interpolation to the nearest metadata bin. We found that this analysis produced a clear distinction between false positives (low quality score RNAs) and true positives (high quality score RNAs) with relatively few RNAs with intermediate quality scores (**Figure S1B**). Therefore, the actual choice of quality score threshold did not have a strong effect on the RNAs that were kept in our MERFISH analysis. We selected 0.8 as a uniform quality score threshold applied to all datasets.

In-house bulk RNA sequencing revealed a small set of genes for which the MERFISH expression level is much higher than the expected bulk RNA sequencing level (**Figure S1C**). To minimize the effect of residual false positives on our biological interpretation, we screened our data for RNAs which had evidence of large false-positive contamination by looking for RNAs detected more frequently than expected given bulk RNA-seq. We used a linear regression in logarithmic expression space to remove the predicted abundance from bulk RNA sequencing from that observed with MERFISH and identified outlier genes as those with residuals post-regression above 0.8. In addition, we removed all zero-FPKM genes expressed in any MERFISH cluster with an average abundance greater than 0.002. In total, this analysis flagged 106 genes as potentially more contaminated by false positives than the average gene. We report the location and identity of these genes but, except for *Mki67* whose expression was validated by other methods, these genes were not included in the reported single-cell analysis.

#### Cell segmentation

To assign RNAs to individual cells, we used both the immunofluorescence cell boundary stain as well as RNA identity and spatial distribution patterns. We applied 2D Cellpose to create 3D boundaries as described previously^36^. Briefly, we first trained a Cellpose^38^ model on the Na^+^/K^+^-ATPase membrane immunofluorescence (IF) from 11 FOV in 3 regions (ileum, proximal colon, and distal colon) in SPF mice, starting with the “cyto” pretrained model. Next, we applied this Cellpose model to each z-plane for each FOV, producing 2D cell boundaries. Then, we grouped 2D boundaries for the same cell across all z-planes in each field of view if pairs of boundaries within adjacent z-planes shared at least 60% of the same area by the intersection over union (IoU) metric. RNAs were then assigned to these putative cells if they fell within these 3D boundaries. The final Cellpose model is provided in the Dryad repository associated with this work.

To refine the segmentation and capture cells for which Cellpose did not define a boundary, we used Baysor^36^ (v0.5.1). Specifically, RNAs that passed the quality score threshold were given to Baysor with the identity of the Cellpose cell to which it was assigned, if any. All blank barcodes were excluded from this analysis, as was *Neat1*, a highly abundant nuclear RNA that we found could artificially drive the segmentation of nuclei from cytoplasm in the same cell. Since the quality of Baysor segmentation can be sensitive to its parameters, we first scanned a range of parameters on a subset of our data comprising three slices, one each from ileum, cecum, and distal colon. Specifically, we explored size scale (-s, from 3 to 5 in increments of 0.5), minimum molecules per cell (-m, 10 or 15), number of initial clusters for coarse segmentation (--n-clusters, from 4 to 30 in increments of 2), and Cellpose prior confidence (--prior-segmentation-confidence, from 0.7 to 1 in increments of 0.05).

For each parameter set in this screen, we used the scRNA-seq doublet-detector tool Scrublet^39^ to calculate a doublet score for each cell. We selected the final parameter set that minimized this doublet score. As a validation of this iterative segmentation approach, we observed that the qualitative quality of the UMAP of this dataset was noticeably improved with the final parameter set identified here as compared to our initial unguided best estimate of the parameter choice. Baysor was then run on all datasets with these optimized parameters (-i 1000 -m 15 -s 3.5 -- scale-std=50% --prior-segmentation-confidence=0.9 --n-clusters=10 --exclude- genes=Blank*,Neat1 --no-ncv-estimation).

To further improve our segmentation, we identified RNAs assigned to cells with limited assignment confidence by Baysor and removed all RNAs with assignment confidence less than 0.5. In total, 16% of all RNAs were removed from cells by this filtering criteria.

To evaluate the quality of our segmentation, we performed a ‘marker purity analysis.’ Specifically, we quantified the fraction of cells in each major cell class that expressed well established cell type markers. For a more stringent quality threshold, we defined expression as a single count of RNA per cell. Indeed, we found that these markers were highly enriched in the expected cell types, with very low levels of expression elsewhere. However, this analysis did show that despite our best efforts, there may remain some small degree of cross contamination of expression between adjacent cells due to imperfect segmentation.

Baysor assigns RNAs to cells but does not, technically, find the boundary of cells. Thus, when boundaries were needed for display or other calculations, we estimated these boundaries from the 2D convex hulls calculated from the location of all RNAs assigned to a given cell across all z-planes. The python packages descartes and shapely were used for convex hull calculations.

#### Sequential FISH signal quantification

To quantify the expression of the genes measured with sequential FISH, we calculated the average intensity for the pixels found within the 2D hull associated with each cell for the measured z-plane. While we included these genes to help discriminate some cell subtypes, we found that the genes measured with combinatorial barcodes were sufficient to define these subtypes. Therefore, the sequential markers (reported as z-scores of the average in-cell intensity) were only used as confirmation of enteroendocrine subtype labels. Signals for one gene, *Gip*, appeared to co-localize with the signals for *Muc2*, a marker for Goblet cells, likely due to non-specific readout probe binding. In addition, two additional markers, *Vil1* for enterocytes and *Mptx2* for Paneth cells, displayed high background fluorescence. As none of these four genes (*Gip*, *Muc2*, *Vil1*, *Mptx2*) proved necessary for defining their respective cell type clusters, we excluded them from all subsequent analyses.

#### Single-cell clustering and cell type assignment

To identify cell classes and cell types in our measurement, we first filtered cells based on size and transcript count using the following thresholds: an integrated area across all z planes within the bounds of 12 μm^2^ to 500 μm^2^, total RNA copy counts between the bounds of 15 and 2000, and a minimum number of unique genes of 5. To further reduce doublet effects from segmentation, a Scrublet^39^ doublet score was then calculated for each cell, and the top 10% of cells with the highest doublet score (above 0.22) were removed from the dataset. After this filtering, our atlas contained 2.1 M cells and 280 M transcripts.

To identify cell populations, we used standard single-cell analysis approaches as implemented in the Scanpy^82^ package (v1.9.0). Specifically, we normalized the counts within each cell to a target total sum of 2000 counts. A pseudocount of 1 was then added to all genes in all cells and the expression was natural log transformed to create logarithmic expression. In addition, we used linear regression to remove any potential effects due to the total number of unnormalized counts from this logarithmic expression, and then the residuals from this regression were z-scored.

The top 1,300 highly variable genes were used for principal component analysis (PCA). To select the number of principal components (PC) to keep for subsequent steps, we performed a pseudo-JackStraw analysis in which the maximum eigenvalue was determined from a PCA performed on expression measurements in which the cell indices associated with each gene were scrambled (therefore removing gene-gene expression covariation while preserving the expression distribution for each gene) for 10 independent randomizations. We selected only the PCs from the original analysis that had eigenvalues greater than the average of these 10 maximum values. The selected PCs were then batch-corrected with Harmony to remove batch effects between datasets, allowing for a joint analysis of cell types despite regional and microbiome-dependent differences^40^.

UMAP representations and Leiden clusters were calculated using standard approaches in Scanpy. This analysis was performed in multiple tiers to capture the natural hierarchy of cell types. Briefly, a Tier 1 UMAP was constructed from the harmonized PCs using a nearest neighbor graph with 49 PCs, 30 neighbors, a min_dist value of 0.3, and the cosine distance metric. This nearest neighbor graph was then used to construct clusters with the Leiden algorithm with a resolution value of 1.5. The expression of canonical marker genes for major cell classes—epithelial, fibroblast, immune, endothelial, interstitial cells of Cajal, smooth muscle, cells of the enteric nervous system, and adipose—were used to identify clusters associated with these classes. In the Tier 2 clustering, this process was repeated for each cell class individually to define cell types and states within these classes. In some cases, clusters were manually regrouped as a single Leiden resolution was insufficient to define all divisions within a class without introducing additional population divisions. Several cell populations (including EECs and T cells) were put through a third tier of clustering to identify established divisions within them.

All identified clusters were present across multiple replicate mice and could be assigned to expected cell populations based on gene expression and spatial location. However, a few clusters of epithelial cells were strongly driven by imaging artifacts in small regions of a few slices; these clusters were removed from subsequent analysis. In addition, we identified a population of cells that co-clustered with fibroblastic reticular cells (FRCs) but were found outside of the expected location in gut associated lymphoid tissues (GALTs) and expressed lower levels of canonical FRC markers. We labeled these cells FB6 and included them in all reported data, but as we cannot rule out that these cells represented a segmentation or clustering artifact, we do not discuss them further. Finally, we noted that some base-crypt goblet cells in the large intestine were grouped with the Paneth cell cluster in ileum, likely reflecting some shared gene expression among base-crypt secretory lineages; however, these cells do not express canonical Paneth markers outside of the ileum and were, thus, not discussed further.

In addition to the marker purity analysis described above, all final cell clusters were examined for evidence of cross-contamination with other cell populations. We noted very modest degrees of such contamination across all cell types. Nonetheless, we did notice that the Tuft cell cluster contained unclustered subsets that showed greater co-expression of markers of other epithelial populations, suggesting that this population may contain a greater degree of doublet-like features, possibly, because Baysor did not see enough instances of these exceedingly rare cells to drive the same degree of high-quality segmentation we observed in other populations.

#### Integration with scRNA-seq

As an additional layer of validation for our clustering and cell type assignment, we co-embedded our MERFISH gut atlas with published single-cell or single-nucleus sequencing datasets. Since large-scale scRNA-seq datasets that comprehensively cover all cell classes in the mouse gut were lacking, we chose, instead, to separately co-embed subsets of our data for different cell classes to existing scRNA-seq datasets of matching cell classes and regions and for which both sufficient cell numbers and cell cluster labels were available. Specifically, we used Haber et al.^8^ for ileal epithelial cells, Wang et al.^13^ for ileal immune cells, Pærregaard et al.^12^ for fibroblasts from the colon and ileum, Drokhlyansky et al.^10^ for cells of the ENS in the colon, Kalucka et al.^81^ for endothelial cells in the colon, Gehart et al.^9^ for ileal EECs, and Mayassi et al.^33^ for enterocytes of the colon that share evidence of the Scnn1g+ enterocytes described below. All scRNA-seq data were from SPF mice.

For each cell class, a set of overlapping genes between the two datasets were identified and the expression matrices concatenated. As described above, this concatenated dataset was normalized, natural log-transformed, regressed for total counts, and z-score corrected. PCA was then performed on 1,300 highly variable genes, and the number of PCs determined by the pseudo-JackStraw method described above. The kept PCs were then batch-corrected with Harmony across the datasets and data types (MERFISH vs. scRNA-seq). These corrected PCs were then used to create UMAP representations as described above.

To assess the co-integration between MERFISH and scRNA-seq, we computed a multi-categorical confusion matrix to evaluate the cell type alignment between these two data types. Specifically, for each MERFISH cell, we identified its five nearest scRNA-seq neighbors (NNs) in the batch-corrected PC space by the cosine distance metric, using only PCs kept by the pseudo-JackStraw method. Then a probability density distribution of the scRNA-seq cell types of the five NNs was calculated, using the inverse cosine distance in the PC space as weights, and aggregated across all cells in the same MERFISH cell type to yield a probabilistic scRNA-seq cell type alignment for that MERFISH cell type. Across all major cell classes, this analysis revealed an excellent level of alignment between the MERFISH labels and the published sequencing labels, validating our data quality, segmentation, clustering, and cell type assignment.

None of these previous studies, including the most recent one from Mayassi et al.^33^, identified a population of *Scnn1g*+ enterocytes in the mouse gut. Moreover, among the published epithelial datasets, only Mayassi et al. provided the necessary markers to define these populations. Therefore, to determine if evidence for this population could be found within the published data of Mayassi et al., we first extracted all colon enterocytes from SPF mice in this dataset and computed the pairwise Pearson correlation coefficients between the z-scored log-normalized expression values for key enterocyte population markers. In parallel, we co-embedded this dataset with SPF colonic enterocytes from the MERFISH atlas and mapped the sequencing cell types onto MERFISH cell types as described above. In particular, we isolated all *Scnn1g*-expressing enterocytes in the Mayassi et al. dataset and treated them as a separate group for the cell type alignment analysis. The enterocytes expressing *Scnn1g* in the Mayassi et al. dataset mapped onto the *Scnn1g*+ enterocytes in the MERFISH atlas, further supporting the existence of this enterocyte state in the mouse colonic epithelium.

#### Spatial neighborhood analysis

To analyze and extricate distinct gut anatomical layers, namely mucosa, submucosa, muscularis externa, myenteric plexus, and gut-associated lymphoid tissue (GALT), we adapted a tissue neighborhood analysis approach that has been previously described^32^, with several important modifications. Specifically, for each cell, we identified the 25 nearest neighbors based on the centroid-to-centroid distances, and a Neighborhood cell-type Composition Vector (NCV) was calculated with an inverse distance weight, which was defined as (10 μm / *d*), where 10 μm was roughly the dimension of the average cell and *d* was the centroid-to-centroid distance between the center cell and its neighbor. The weight was set to 1.0 for the center cell and capped at 1.0 as the maximum possible value for a neighboring cell. To capture both high-level and fine-grained cell type features, the NCV kept track of the relative abundance of both cell classes (Tier 1) as well as the final cell type labels (Tier 2 or Tier 3). The contribution of Tier 2 or Tier 3 components to the NCV was weighted with a factor of 2 relative to Tier 1 components.

The NCVs for all cells in the dataset, including all regions and microbiome conditions, were concatenated to form one neighborhood composition matrix (NCM). To identify spatial neighborhoods, we normalized each NCV to a total sum of 20, z-scored the NCM, performed PCA, and determined the number of PCs to keep via the jack straw method described above. These PCs were then used to create a UMAP embedding with the cosine distance metric, 30 neighbors, and a minimum distance of 0.3. To label different anatomical layers, we performed Leiden clustering on the same graph used to create the UMAP. A relatively modest resolution (0.3) was sufficient to pull out the major anatomical features. This analysis produced a few different mucosal regions, which we manually recombined. As we have observed previously^32^, the high numerical similarity of some NCVs produced challenges in UMAP convergence. Therefore, we added a random, small noise term, which varied between 0.02 and 0.2, to each entry in the NCM to aid convergence. The final spatial neighborhood labels derived from this analysis are provided in the Dryad deposition.

To quantify the relative abundance and distribution of cell types across gut regions and anatomical layers, we performed a three-tiered analysis. First, the relative abundance of cell types was calculated as the fraction of a given cell type among all cells in all SPF regions. Then, the relative abundance of cell types across regions was calculated by dividing the number in each region by the total number of that cell type in all SPF regions. Finally, the relative abundance of each cell type across gut anatomical layers was calculated separately for each SPF region, by dividing the number in each layer by the total number across all layers in that region.

#### Mucosal pseudospace axis

To define a unified spatial axis of the mucosa across regions and microbiome states, we used the spatially organized maturation of epithelial cells—from stem cells at the crypt base to mature enterocytes at the top of the mucosa. Specifically, we subset our atlas to stem cells as well as bottom, mid, top, and *Scnn1g*+ enterocytes, and we calculated a UMAP using harmonized PCs following a similar protocol as above. A diffusion map with 15 dimensions was then calculated using Scanpy based on the resulting nearest-neighbor graph. While the 1^st^ diffusion component captured inter-sample variation, the 2^nd^ and 3^rd^ components reflected the enterocyte maturation trajectory from crypt base to top mucosa. We then computed pseudotime without branching from the diffusion map of the 2^nd^ and 3^rd^ components, using the stem cell with the lowest values on these two components as the root using standard methods in Scanpy.

To extend this spatial trajectory to all mucosal populations, a mucosal pseudospace axis was constructed by propagating enterocyte pseudotime values to nearby non-enterocyte cells. For each non-enterocyte cell in the mucosa spatial neighborhood, the three nearest stem or enterocyte cells (based on centroid-to-centroid distance) were identified, and their pseudotime values averaged and assigned to that cell as its mucosal pseudospace value. For stem and enterocyte populations, the original expression-derived pseudotime was retained as the mucosal pseudospace value.

To relate mucosal pseudospace to real-space location, we computed the cumulative fraction of all mucosal cells at or below a given mucosal pseudospace value for 100 pseudospace positions sampled evenly between 0 (crypt base) and 1 (top of mucosa). The length fractions were then converted to real-space length values by scaling by the average mucosal thickness of the given region and microbiome state. To calculate the mucosal thickness, we identified the top and bottom 5% of all cells ranked by mucosal pseudospace coordinates. Each top cell was then mapped to the 5 nearest bottom cells, and the distances calculated and averaged; the distances were then averaged across all top cells in each slice to derive the slice-averaged mucosal thickness. Mapping between mucosal pseudospace and real-space length position was quantified by averaging the thickness values slice-by-slice within each region and microbiome state, followed by computation of the mean and standard error across slices.

While the spatial neighborhood analysis described in the previous section was used to delineate the mucosa from surrounding regions, the mucosal pseudospace axis provided a more fine-grained descriptor of mucosal spatial organization. Accordingly, for all analyses referring to the bottom, mid, or top mucosa, regions were defined based on mucosal pseudospace values: bottom as < 1/3, mid as 1/3 – 2/3, and top as > 2/3.

#### Cellular and transcriptomic gradients in the mucosa

Real-space coordinates, derived in relation to mucosal pseudospace coordinates as described above, were used to quantify the relative spatial distribution of all cell populations within the mucosa, separately for each region and microbiome state. The real-space coordinates were favored over the pseudospace coordinates for this analysis as they provided a more intuitive visualization of cell abundance distributions in the mucosa. To ensure robust estimation, the analysis was restricted to cell populations exceeding a minimum abundance threshold of 100 cells in the given region and microbiome condition. For each cell type, kernel density estimate (KDE) was applied on the distribution of mucosal real-space coordinates. KDE was implemented by the scipy.stats.gaussian_kde module and the kernel bandwidth was automatically selected via the “scott” method. To avoid artificial disruptions to the cell densities near either ends of the real-space axis, boundary correction was applied by padding the distribution of real-space values for 0.1 fractional unit beyond the minimum or maximum real-space value. Specifically, all data points within 0.1 fractional unit of either end of the real-space axis were reflected across the end and added to the existing real-space value distribution. The KDE-smoothed cell density vectors were normalized so that the maximum value equaled 1, visualized with heat map, and reported for mucosal cell types in both SPF and GF gut regions.

Similar to cell abundance gradients, gene expression gradients along the mucosal pseudospace axis were estimated using KDE. In short, in each mucosal cell type with more than 100 counts in a given region or microbiome state, RNAs of each gene were tabulated for the mucosal pseudospace values of the cells they are found in. The resulting pseudospace values were then smoothed with KDE and normalized by the cell abundance KDE over the same range of mucosal pseudospace coordinates.

To implement this pipeline, several considerations were made. First, since sufficient cell density was required for robust KDE estimate of gene expression, for each mucosal cell type in a given region or condition, only the mucosal pseudospace range over which the given cell type was sufficiently abundant was used for gene expression gradient calculations. To determine the pseudospace range where cell density is sufficient, the following standard was applied. Cell counts were first computed in 100 bins along the pseudospace axis. The lower boundary of the range was defined as the lowest pseudospace value above which all bins contained at least 10 cells. Similarly, the upper boundary was defined as the highest pseudospace value below which all bins satisfied this same threshold. KDE analyses proceeded only if the resulting valid range exceeded 0.1 pseudospace unit (the whole mucosa is 1 pseudospace unit), ensuring adequate support for reliable cell density and gene expression estimation.

To estimate cell densities for the purpose of normalizing gene expression, a KDE was computed for the total counts of the selected cell type along the pseudospace axis, following a protocol like the one described above. Genes were then filtered so that only ones with natural log-normalized expression levels above 0.2 in each cell type in the given region and microbiome state were estimated for KDE densities. To estimate gene expression densities, for each gene, a transcript count vector and a pseudospace coordinate vector were extracted from all cells of a given type. The pseudospace vector was expanded such that each pseudospace value was replicated according to the number of transcripts in the corresponding cell. If the resulting vector contained more than 100 values, a KDE was applied to obtain a continuous expression profile along the pseudospace axis. This profile was then normalized by the KDE of total cell counts to control for underlying cell density. The normalized curve was further smoothed using a second round of KDE and finally scaled so that the maximum value was equal to 1. To visualize the gene expression gradients in enterocytes, telocytes, and macrophages, hierarchical clustering with Euclidean distance metric and “complete” inter-cluster distance was applied for the purpose of ordering the gradients by similarity. The final set of gene expression gradients for each cell type in each region and microbiome state was reported in **Table S2**.

#### Mapping mucosal pseudospace to scRNA-seq

To validate mucosal gene expression gradients independently of MERFISH measurements and to extend these gradients transcriptome-wide, we mapped mucosal positional coordinates onto published scRNA-seq datasets via co-integration with MERFISH data. Integration was performed as previously described. For each cell in the scRNA-seq dataset that corresponded to a mucosal cell type, the five nearest MERFISH neighbors of the same type were identified within the co-embedding principal component space by the Euclidean distance metric, and mucosal pseudospace values were transferred by averaging the positional scores of these neighbors.

Following pseudospace transfer, each scRNA-seq dataset was analyzed independently of MERFISH data to ensure an unbiased assessment of transcriptome-wide spatial patterns. Preprocessing steps included count normalization to a target sum of 10,000, natural log-transformation, and selection of highly variable genes (min_mean = 0, min_disp = -0.2). Total UMI counts were regressed out, and the residual expression matrix was z-scored. Principal component analysis (PCA) was performed, and the number of components retained was determined using the pseudo-JackStraw method as described above. Batch correction was applied using either Harmony^40^ or ComBat^88^, with the latter used when average batch sizes were fewer than 200 cells due to poor Harmony performance under such conditions. A nearest-neighbor graph and UMAP embedding were then computed using standard Scanpy procedures. The transferred mucosal pseudospace values were overlaid on the scRNA-seq UMAPs, revealing spatial gradients consistent with the axis of gene expression variation.

#### Mutual information (MI) between transcriptome and pseudospace

To quantitatively assess the influence of mucosal position on gene expression across different mucosal cell populations, latent mutual information (LMI) was computed^58^. LMI calculations were performed using the latentmi package. For each cell population within each mucosal region and microbiome state, cells were subsetted and included only if at least 100 cells were present. The model was run in GPU mode with the following parameters: 15 dimensions for the latent space (N_dims = 15), half the data used as the validation set (validation_split = 0.5), and a maximum number of 2000 rounds of iterations (epochs = 2000). The resulting LMI scores represent the degree to which transcriptomic variation is explained by spatial positions along the mucosal axis.

To identify genes that contribute to spatial expression gradients within specific cell populations along the mucosal pseudospace axis, we calculated the normalized mutual information (NMI) between the expression of individual genes and mucosal pseudospace. For each gene within a given cell population and condition, we estimated mutual information using sklearn.feature_selection.mutual_info_regression. To normalize the resulting mutual information values to fall between 0 and 1, we divided them by the square root of the product of the differential entropies of pseudospace locations and gene expression, both computed using scipy.stats.differential_entropy. Prior to these calculations, both pseudospace and natural log-normalized gene expression values were z-scored to ensure scale comparability. To handle the entropy of gene expression distributions containing a large fraction of zeros, we modeled expression as a two-component mixture of zero and non-zero values, assigning zero entropy to the former and estimating the latter after adding a small noise term, modeled with a Gaussian distribution with 0 mean and 10^-6^ variance, to avoid numerical instability. NMI was computed only for cell populations with at least 100 cells in the given region and condition, and for genes with average natural log-normalized expression greater than 0.2. The distributions of NMI values in select cell populations in SPF ileum were visualized. The LMI values of cell populations as well as NMI values of genes can be found in **Table S3**.

#### Mucosal gene gradient conservation analysis via hierarchical clustering

To assess the conservation of spatial gene expression gradients in the mucosal pseudospace across mucosal regions and microbiome states, we performed hierarchical clustering of the KDE-smoothed gene expression gradients previously derived. To ensure robust gradient estimation as well as to facilitate comparison across conditions, only cell types with well-defined gradients over the pseudospace range of 0.35 - 0.85 in all regions and microbiome conditions were considered for this analysis. This led us to focus on key populations, including enterocytes, goblet cells, telocytes, and FB1 interstitial fibroblasts. In each of these populations and for each region and microbiome state, only genes above 0.2 natural log-normalized expression were considered for gradient pattern comparison.

For comparison of gradient similarity across regions in the SPF dataset and between SPF and GF, all gene expression gradients were concatenated and hierarchically clustered using a complete linkage algorithm and Euclidean distance. This analysis identified five major gradient patterns: constant, slightly increasing, increasing, decreasing, and slightly decreasing. While a small subset of genes in enterocytes exhibited non-monotonic (e.g., peak in the middle) expression, these were too infrequent to form a distinct cluster.

To quantify the conservation of gradient patterns across SPF regions, we computed the frequency with which a given gene-in-cell type gradient assigned to a particular cluster in one region co-occurred with the same or alternative clusters in other regions. Among genes defined in at least two SPF regions, most cluster transitions occurred between the constant and slightly increasing or slightly decreasing clusters, indicating high overall conservation of gradient patterns. A similar co-occurrence analysis comparing SPF and GF states within the same region revealed a high level of gradient preservation across microbiome conditions.

#### Quantifying mucosal gradient similarity via Jensen-Shannon distance

To quantify the conservation of gene expression gradients across regions and microbiome states, we used the Jensen-Shannon (J-S) distance, a symmetric measure of dissimilarity between density distributions, implemented by the scipy.spatial.distance.jensenshannon module. For each cell type (stem cell and enterocyte, goblet, telocyte, and FB1), we first computed the pairwise J-S distances between all genes expressed above a threshold (0.2 in natural log-normalized expression) within each individual region and microbiome state, capturing the natural intra-regional variation in gene expression spatial patterns. We then computed inter-regional J-S distances for each gene, comparing its expression gradients across all regions in the SPF dataset where it was sufficiently expressed. As expected, inter-regional J-S distances for the same gene were consistently lower than intra-regional distances between different genes, indicating that most gene gradients are conserved across regions.

To identify genes with high regional variability, we defined outliers as those whose inter-regional J-S distance exceeded the 75th percentile of the intra-regional J-S distance distribution of all regions. Genes were then classified as conserved or variable within each cell type and receptor class and visualized using stacked bar plots. The majority of gene gradients were found to be regionally conserved.

A similar approach was applied to compare spatial gene expression gradients between SPF and GF conditions within the same region. J-S distances between microbiome states were markedly smaller than baseline intra-regional variation. Upon applying the same threshold—defined as the 75th percentile of inter-regional and inter-microbiome variation—to identify gradient variability, the vast majority of gene expression gradients were likewise classified as conserved. This reinforced the conclusion that the majority of gene expression gradients were conserved across both regions and microbiome states.

Finally, to quantify the spatial co-localization between chemokine expression and immune cell abundance in the mucosa, we computed the J-S distance between the spatial distributions of chemokines and chemokine-receptor-expressing immune cells along the real-space mucosal axis. The spatial distributions were computed following the KDE-smoothing protocols for cell abundance and gene expression as described above. Chemokine distributions were calculated as density (counts per cell) among all mucosal cells, while immune cell distributions were derived from total cell counts. Chemokine-receptor interactions were curated from the OmniPath^89^ database, with human gene names converted to mouse orthologs using pybiomart and verified manually. Only immune cell populations expressing the cognate chemokine receptor above a threshold of 0.2 in natural log–normalized expression were included. For chemokines, a lower expression threshold of 0.01 was applied to account for dilution across mucosal cell types. J-S distances between chemokine and immune cell distributions were calculated separately within each gut region in the SPF dataset to characterize region-specific spatial coordination between chemokine expression and immune infiltration.

#### Spatially informed receptor-ligand interaction

To perform a spatially informed receptor-ligand interaction analysis, we first subset to all cells found within each mucosal neighborhood, defined by three equal bins of the mucosal pseudospace axis as described above. CellPhoneDB^90^ receptor-ligand interaction analysis was used to map cell-cell communications within each anatomical and functional zones separately for each SPF and GF regions. The following parameters were applied: an expression threshold of 0, meaning we performed permutation tests on all receptor-ligand pairs; 1000 permutations; FDR correction on the p-values; the ‘clusters’ option for the correction axis; and an alpha value of 0.05 for significance. Interactions between mucosal fibroblasts (telocytes and FB1) with stem cells and enterocytes as well as macrophages were extracted and plotted, separately for each region, microbiome state, and mucosal zone. Spatially informed ligand-receptor interaction data for the selected cell type interactions were reported in **Table S4**.

#### Differential gene expression

To identify transcriptomic differences across SPF gut regions and between SPF and GF conditions, we performed differential gene expression (DEG) analysis using individual gut slices as biological replicates. For regional comparisons, each SPF region was compared to the aggregate of all other regions. For microbiome-dependent comparisons, each SPF region was compared separately to its GF counterpart. Within each condition, we first subset cells by region, microbiome state, and cell type, then gene expression across cells of the same type was averaged within each slice. These per-slice values were used to calculate the mean expression in each condition and to estimate the statistical significance of differences between conditions.

To compute the log-fold expression change, the ratio between the averaged normalized expressions (in linear scale, not the log-transformed or z-scored values) was calculated between any given regions or microbiome states, followed by log_2_ transformation of this ratio. To determine the significance of this ratio, a two-sided Student’s t-test was applied to calculate a p-value from the per-slice expression values for each condition. These p-values were then corrected for multiple hypothesis testing using the Benjamini-Hochberg method with a false-discovery rate (FDR) target of 0.05.

For tabulating genes that are differentially expressed across regions or between microbiome states, we defined meaningful DEGs as those with a log-normalized expression above 0.2 in either condition, at least a 2-fold change in expression, and an FDR-corrected p-value below 0.05. For both sets of comparisons, a gene was considered a regional- or microbiome-dependent DEG if it satisfied these criteria in any cell type in each cell class. For regional comparisons, a gene was counted as a DEG if it was upregulated in at least one region relative to the rest. The fraction of genes within each receptor and cell class that are DEGs across regions or between microbiome conditions were plotted. The full list of DEGs is provided in **Table S6**.

#### Receptor and microbiota metabolite database integration

To investigate how microbiota metabolites interact with the gut in a spatially resolved and cell-type-specific fashion, we integrated our receptor atlas with a microbiota metabolite-receptor database we assembled from a variety of published databases as described below.

To construct a microbiota-metabolite-receptor database, we started with the gutMGene^67^ microbiota-metabolite-receptor database, which compiles literature associations between microbial metabolites and receptors. To complement this existing database, we took two comprehensive receptor-ligand databases, GLASS^64^ (for GPCRs) and NRLiST BDB^65^ (for nuclear receptors), and screened the ligands based on their presence in a comprehensive microbiota metabolite database, MiMeDB^66^, keeping only interactions in GLASS or NRLiSt where the ligand was a metabolite in MiMeDB. We then examined all target genes in gutMGene to ensure that they were a receptor gene instead of a gene in a downstream pathway. To enrich this final database, we supplemented it with chemical taxonomy information derived from HMDB^91^ to allow metabolites to be explored based on common chemical categories (e.g., short-chain fatty acids, alkylamines, etc.). We termed our final database the Microbiota Metabolite-Receptor Database (MMRDB; **Table S7**).

To integrate MMRDB with our MERFISH receptor atlas, we used the drug2cell pipeline^68^. To explore potential targets of microbiota metabolites based on our measured receptor profiles, we provided the MMRDB to drug2cell as a Python dictionary. We translated mouse gene names to human names using the python package mousipy and ran drug2cell on the expression profiles measured by MERFISH for all cells seen in SPF mice. Metabolite – cell interaction scores from drug2cell were used for downstream analysis, with one important modification: for a compound with multiple target receptors expressed in the same cell, instead of averaging the interaction scores as the default in the drug2cell pipeline, we summed the interaction scores. This modification was made to ensure that weak interactions between a compound and some receptors would not mask strong interactions between the same compound and other receptors expressed in the same cell type.

To tabulate the percentage of microbiota metabolites that could target each of the major gut cell classes (epithelial, immune, fibroblast, endothelial, smooth muscle, ENS, or ICC), a metabolite was considered as interacting if its interaction score with any cell type in a given cell class in any given region in the SPF dataset was above a threshold of 0.5. In order to identify the most targeted cell classes of select microbiota metabolite categories, or the most targeted cell types of select metabolites, we calculated z-scores across all cell types (or classes) of the interaction score with a given metabolite (or metabolite category).

#### Dimensionally reduced spatially variable gene (SVG) analysis

In order to describe the gene expression heterogeneity around the gut cross-sectional circumference, we developed a dimensionally reduced spatially variable gene (SVG) approach that captured the circumferential axis with a 1D coordinate. This method presents a preliminary version of a more sophisticated analysis described elsewhere^92^, yet it proved sufficient for capturing the key features highlighted in our results. Specifically, a modified version of the dimension reduction method Isomap^93^ was used to estimate the 1D manifold that best approximated the 2D coordinates of the cells from each cell type in each major anatomical layer: mucosa, submucosa, and muscularis propria. Isomap began by constructing a k-nearest neighbor (kNN) graph of the 2D cell coordinates and then applied Djikstra’s algorithm to find the shortest path between all pairs of cells. k was set small enough to only include close neighbors, so that this graph geodesic would approximate the manifold distance. Then the graph distance matrix was embedded in one dimension using multi-dimensional scaling (MDS). Specifically, for each cell with coordinates (*x*_*i*_, *y*_*i*_) we obtained a corresponding 0 ≤ *t*_*i*_ ≤ 1 that described the position of that cell along the one-dimensional manifold. In the context of gut cross sections, Isomap sometimes could not be directly applied since the tissue structure was a closed loop. In this case we began by dividing the cells into two semicircles and applied Isomap to each arc independently. Then the two arcs were combined to parameterize the location of the cell along the loop.

Once the one-dimensional parameter *t*_*i*_ was obtained, we analyzed the gene expression data as a continuous series along this variable. Specifically, let *Y*_*ig*_ be the count for gene *i* in cell *g*; we then model

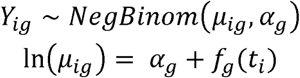

where *α*_*g*_ is baseline expression, *n*_*i*_ is the total number of counts (across all genes) for cell *i*, and *f*_*g*_(*t*_*i*_) is some smooth function of *t*_*i*_. exp (*f*_*g*_(*t*_*i*_)) can be interpreted as the expression fold change of cell *i* relative to baseline. The model is a type of generalized additive model (GAM) and is fit using the mgcv package in R^94,95^. We also apply ash^96^ to shrink the estimate of *f_g_* towards 0 to reduce sensitivity to low count genes. The criteria for finding a gene with patchy expression is based on the *peak:*

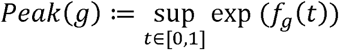

The method also records where the peak occurs, the baseline expression, and the number of local maxima of *f*_g_.

We then filtered the results of this analysis by selecting genes with a peak exceeding 15, an absolute expression greater than 0.03 counts per cell, and with at least two independent occurrences across all SPF slices, yielding a list of 58 unique SVG – cell pairs (**Table S5**). Among them, a large proportion could be attributed to broad variable gene expression that arose due to slice morphology or residual mucosal gradient patterns. However, we also identified genes with non-uniform (“patchy”) expression patterns around the gut cross-sectional circumference, such as the interferon stimulated gene (ISG) patches and genes associated with the follicle-associated epithelium (FAE). These patterns were only apparent in the results for enterocytes but not other cell populations.

