## Supplementary material for "An image-based transcriptomics atlas reveals the regional and microbiota-dependent molecular, cellular, and spatial structure of the murine gut": SI Figures and Captions

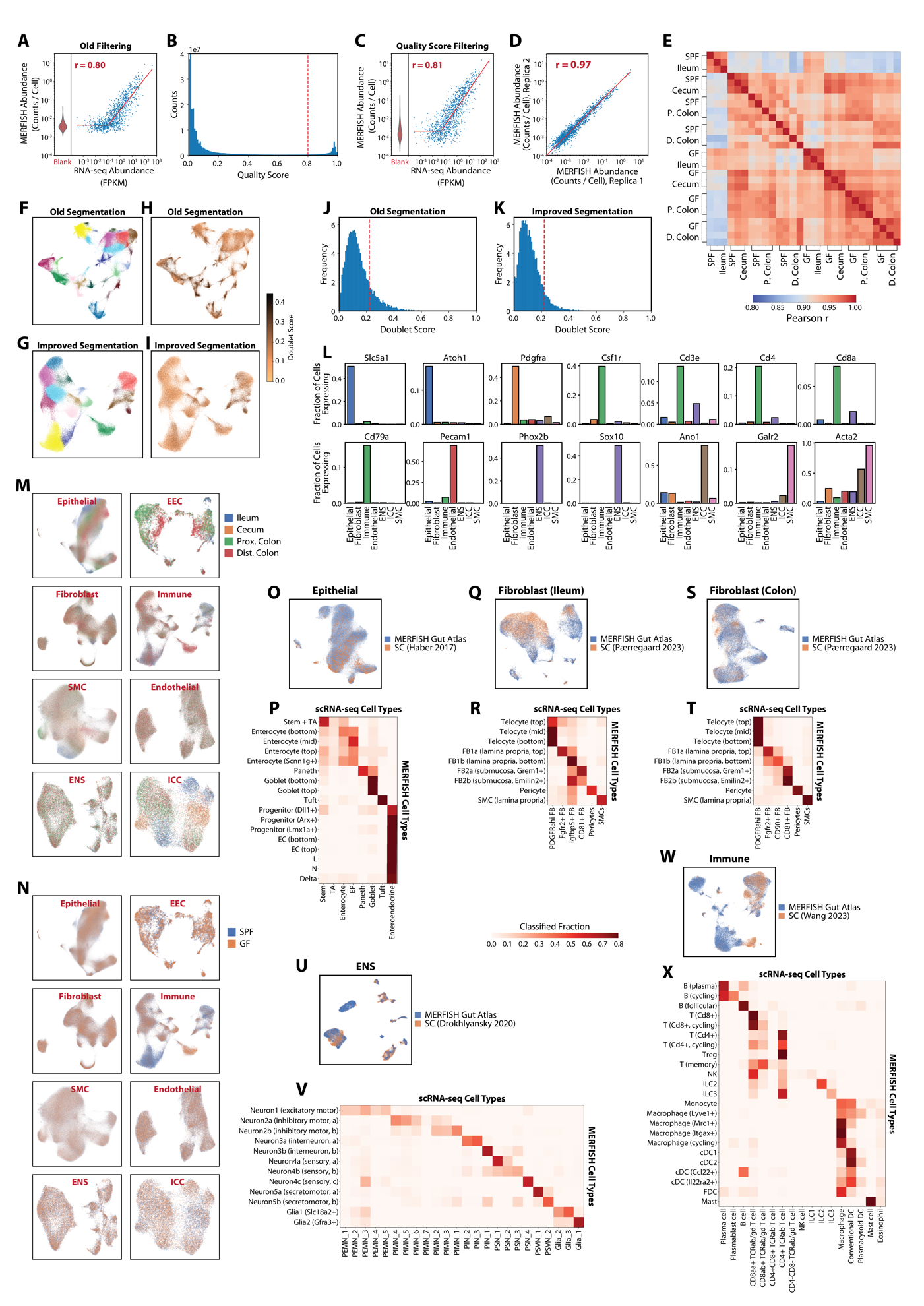

**Figure S1**. **Related to Figure 1. Validation and benchmarking of MERFISH measurements.**

(A) Scatter plot of the average RNA copy number per cell determined via MERFISH versus bulk RNA-seq from the distal colon of SPF mice (right) and the observed distribution of copy number per cell of the ‘Blank’ (control) barcodes determined via MERFISH in the same measurement (left). Bulk RNA-seq abundance is measured in fragments per kilobase per million reads (FPKM). The Pearson correlation coefficient between the logarithmic expression values is listed (r). The red line is a guide to the eye. The MERFISH counts per cell were determined via the old filtering method^37^. Only RNAs with non-zero FPKM are plotted.

(B) Histogram of quality scores for all MERFISH barcodes in the same dataset as shown in (A). The dashed red line represents the quality score threshold used throughout this work to discriminate foreground from background molecules.

(C) As in (A) but for the average copy number per cell determined via MERFISH after the new quality-score filtering method is applied.

(D) Scatter plot of the average RNA copy number per cell determined for one MERFISH measurement of the SPF ileum versus another measurement of the SPF ileum taken from a different mouse. The Pearson correlation coefficient between the logarithmic expression values is listed (r).

(E) The pairwise Pearson correlation coefficients measured between the logarithmic expression values for all MERFISH measurements. SPF: Specific-pathogen free. GF: Germ-free. P. colon: Proximal colon. D. colon: Distal colon.

(F-I) UMAP representation of a subset of cells imaged with MERFISH colored by Leiden cluster (F,G) or via the measured doublet score (H,I) prior to (F,H) or after (G,I) optimization of Baysor segmentation parameters.

(J,K) Histogram of the doublet score measured for the data in (F-I) before (J) and after (K) optimization of Baysor segmentation parameters.

(L) Fraction of each of the major cell classes expressing even a single count of the listed marker genes for these classes. Listed markers include those of epithelial (*Slc5a1* and *Atoh1*), fibroblast (*Pdgfra*), immune (*Csf1r*, *Cd3e*, *Cd4*, *Cd8a*, and *Cd79a*), endothelial (*Pecam1*), the enteric nervous system (ENS; *Phox3b* and *Sox10*), interstitial cells of Cajal (*Ano1*), and smooth muscle (*Galr2* and *Acta2*) populations.

(M,N) UMAP representation of all cells for each of the major cell classes identified in Tier 1 clustering (**Figure 1C**) colored by gut region (M) or microbiome state (N).

(O) UMAP representation of the co-embedding of epithelial cells measured in this work (MERFISH Gut Atlas) and scRNA-seq published elsewhere^8^ (Haber 2017). Both datasets are from SPF ileum.

(P) Fraction of published scRNA-seq cluster labels in the vicinity of cells assigned the given MERFISH label in the co-integration in (O) (STAR Methods).

(Q,R) As in (O,P) but for SPF ileal fibroblasts with data from (Paerregaard 2023)^12^.

(S,T) As in (O,P) but for SPF colonic fibroblasts with data from (Paerregaard 2023)^12^.

(U,V) As in (O,P) but for SPF colonic cells of the enteric nervous system with data from (Drokhlyansky 2020)^10^.

(W,X) As in (O,P) but for SPF ileal immune cells with data from (Wang 2023)^13^.

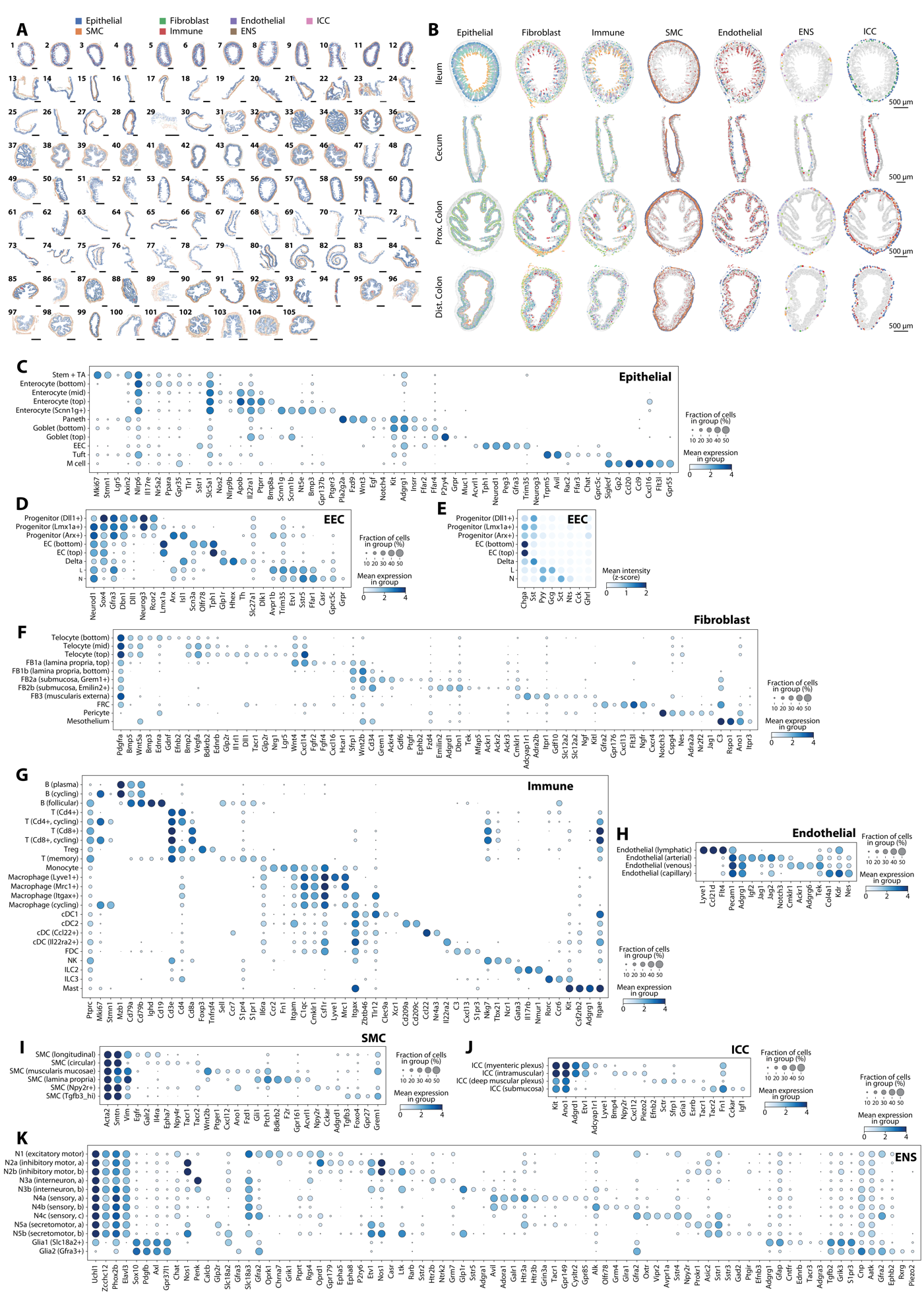

**Figure S2**. **Related to Figure 1. Cell Type Marker Expression and Spatial Distributions.**

(A) Spatial distribution of all major cell classes in all measured slices in this atlas, colored by cell class. Slice 1-12: SPF ileum. Slice 13-30: SPF cecum. Slice 31-37: SPF proximal colon. Slice 38-46: SPF distal colon. Slice 47-62: GF ileum. Slice 63-83: GF cecum. Slice 84-93: GF proximal colon. Slice 94-105: GF distal colon. SPF: specific-pathogen-free. GF: germ-free. ENS: enteric nervous system. SMC: smooth muscle cell. ICC: interstitial cell of Cajal. Scale bars: 500 µm.

(B) Spatial distribution of cell centroids in four representative slices of the SPF mouse with cells colored by cell types within each major class as in **Figure 1D**. All other cells are marked in gray. Scale bars: 500 µm.

(C) Average gene expression within each of the epithelial cell clusters across all regions of SPF mice. Color indicates the average logarithmic normalized expression within the cluster and dot size indicates the fraction of cells that express at least one count of the listed gene.

(D) As in (C) but for enteroendocrine cells (EEC).

(E) The average fluorescence intensity measured within different EEC types for the sequential stains associated with abundant EEC subset markers. Expression is measured in z-score (STAR Methods).

(F) As in (C) but for fibroblasts.

(G) As in (C) but for immune cells.

(H) As in (C) but for endothelial cells.

(I) As in (C) but for smooth muscle cells (SMC).

(J) As in (C) but for interstitial cells of Cajal (ICC).

(K) As in (C) but for the enteric nervous system (ENS). N: neuron.

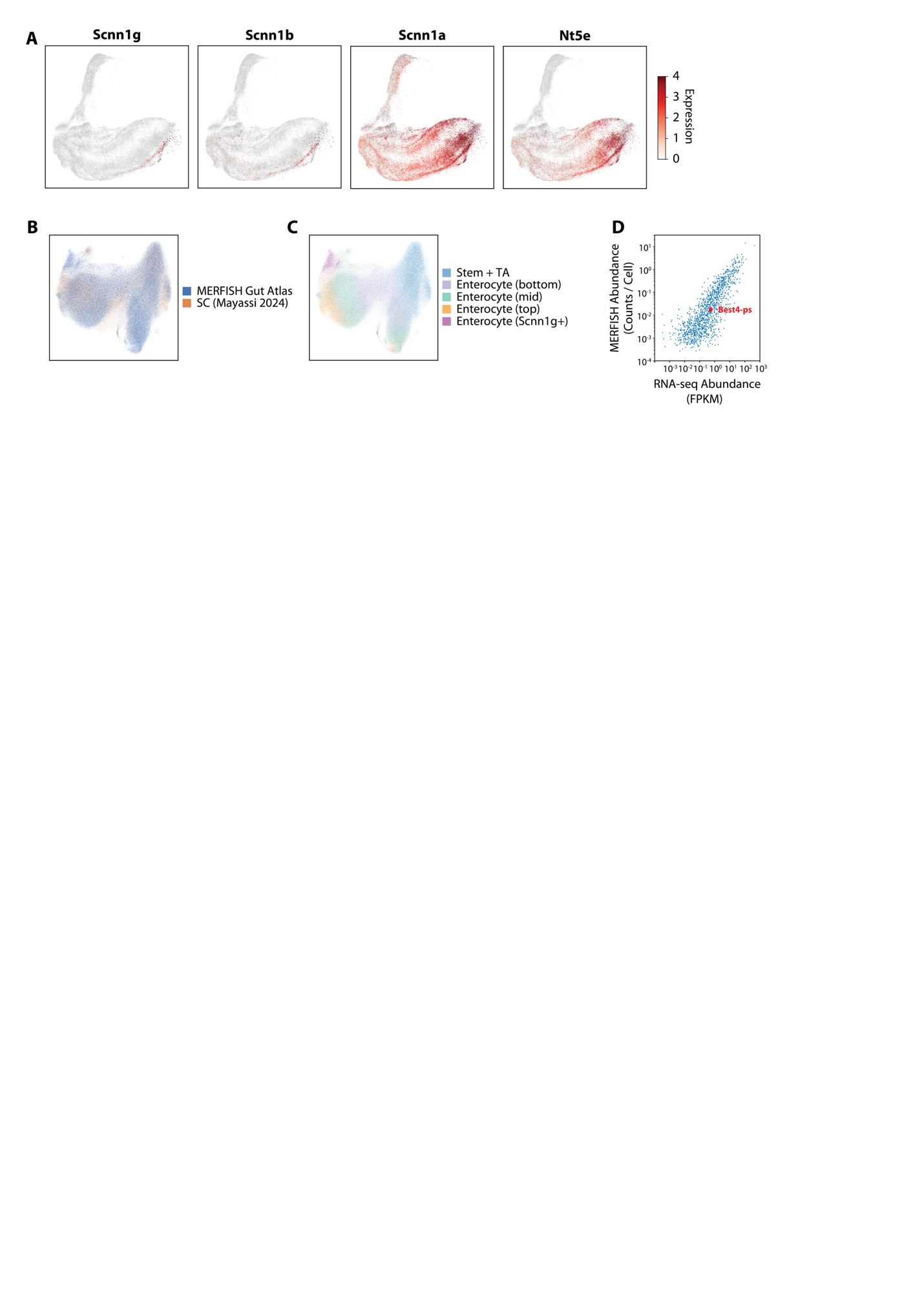

**Figure S3**. **Related to Figure 2. Additional support for Scnn1g+ enterocytes.**

(A) UMAP representation of colonic enterocytes from SPF mice take from published scRNA-seq in Mayassi et al.^33^ Cells are colored by the log-normalized expression of the listed genes. A subpopulation of mature enterocytes co-expressing *Scnn1a*, *Scnn1b*, *Scnn1g*, and *Nt5e* is visible. *Best4-ps* was not reported in this study.
(B) UMAP representation of the co-embedding of colonic enterocytes measured in this work (MERFISH Gut Atlas) and reported by Mayassi et al.^33^ colored by dataset.
(C) As in (B) but colored by labels assigned to MERFISH cells or propagated to scRNA-seq cells.
(D) Scatter plot of the average RNA copy number per cell determined via MERFISH versus bulk RNA-seq from the distal colon of SPF mice. Reproduced from **Figure S1C**. The expression of *Best4-ps* is highlighted.

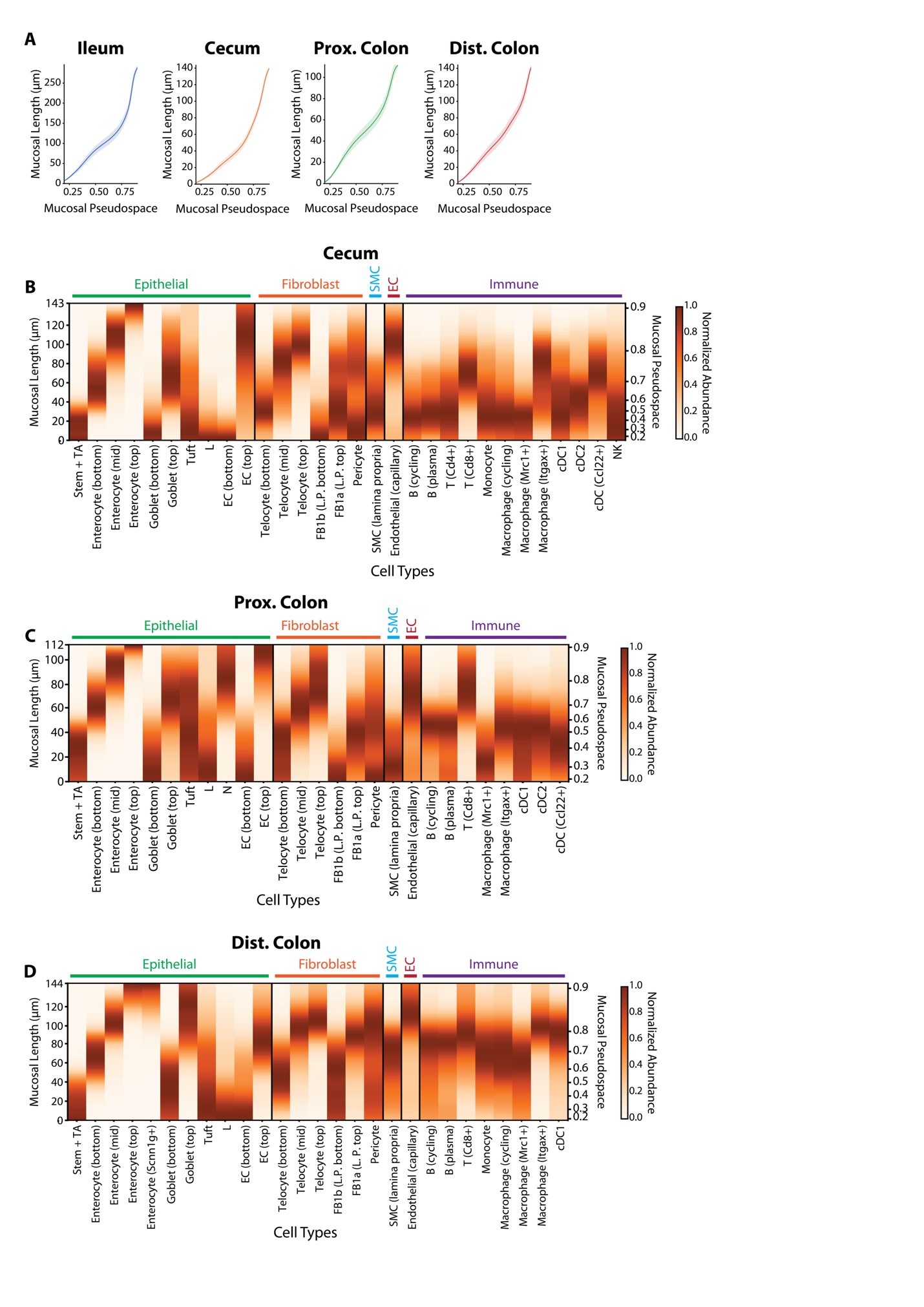

**Figure S4**. **Related to Figure 3. Spatial organization of cell types in the intestinal mucosa in SPF mice.**

(A) The relationship between mucosal pseudospace and mucosal distance from the base of the mucosa for each of the measured regions in the SPF gut. Lines represent the average across all slices and shaded regions represent the 95% confidence interval (slice numbers in **Figure S2A**). Similar to **Figure 3H** but plotting actual, instead of normalized, mucosal length.

(B) Average distribution of all major mucosal cell populations along the length of the cecal mucosa in SPF mice determined by mucosal pseudospace (left) or mucosal distance from the crypt (right). Only mucosal populations with sufficient abundances are shown (STAR Methods).

(C) As in (B) but for the SPF proximal colon.

(D) As in (B) but for the SPF distal colon.

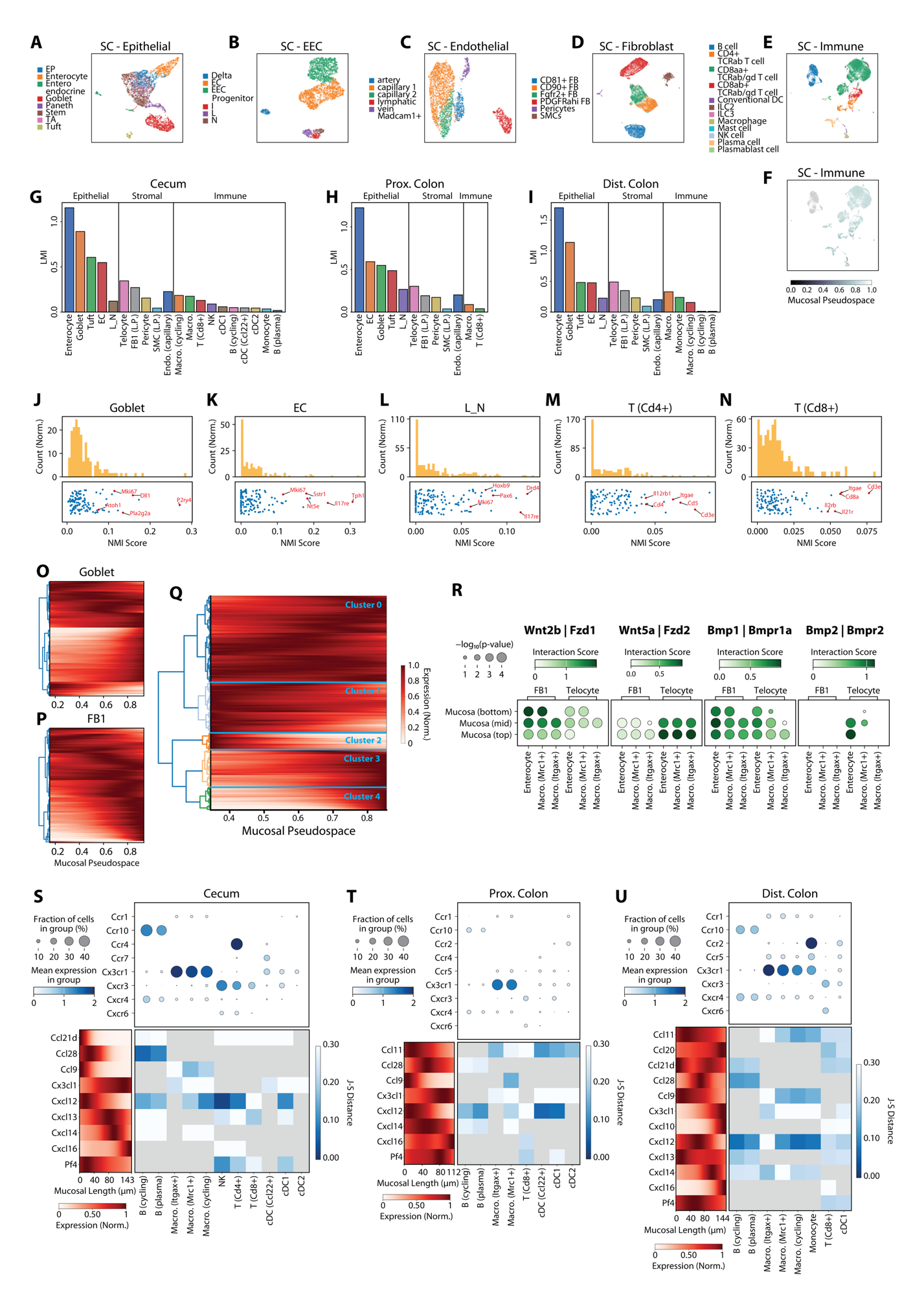

**Figure S5**. **Related to Figure 4. Gene expression variations along the mucosal axis and the potential consequences for regions of the SPF intestine.**

(A) UMAP representation of epithelial populations from Haber et al.^8^ colored by published cell type labels. UMAP is the same as in **Figure 4D**.

(B) As in (A) but for enteroendocrine cells from Gehart et al.^9^.

(C) As in (A) but for endothelial cells from Kalucka et al.^81^.

(D) As in (A) but for fibroblasts from Pærregaard et al.^12^.

(E) As in (A) but for immune cells from Wang et al.^13^. The immune cell UMAP was not included in **Figure 4D**.

(F) UMAP as in (E) colored by the imputed mucosal pseudospace. Cells not found in the mucosa are colored gray. Note that we see modest covariation of mucosal pseudospace and UMAP position, suggesting that for the genes we measured, there is only modest dependence on mucosal position within immune populations.

(G) Latent mutual information (LMI) between gene expression and mucosal pseudospace for all major mucosal cell populations in the SPF cecum.

(H) As in (G) but for the SPF proximal colon.

(I) As in (G) but for the SPF distal colon.

(J) Histogram of normalized mutual information (NMI) scores for all genes in goblet cells in the ileal mucosa of SPF mice (top) with the distribution of NMI for individual genes (bottom). Notable high NMI genes are listed.

(K) As in (J) but for ileal enterochromaffin cells (EC).

(L) As in (J) but for L and N cells, which are known to be the same developmental lineage spatially distributed along the mucosal axis^9^.

(M) As in (J) but for *Cd4*+ T cells.

(N) As in (J) but for *Cd8*+ T cells.

(O) Average expression of individual genes versus mucosal pseudospace for all goblet cells in the ileal mucosa of SPF mice (right) sorted via hierarchical clustering (left).

(P) As in (O) but for all FB1 fibroblasts.

(Q) Average expression of individual genes versus mucosal pseudospace for all enterocytes, goblet cells, telocytes, and FB1 fibroblasts seen in any region of SPF or GF mice (right) sorted by hierarchical clustering (left). The tree is colored by cluster groups of similar spatial gene expression patterns (STAR Methods).

(R) Spatially prioritized receptor ligand interaction scores (STAR Methods) between mucosal fibroblasts (FB1 and telocytes) and different epithelial and macrophage populations for cells in the distal colon of SPF mice. Dot size represents significance while color indicates the interaction score.

(S) Average expression of key chemokine receptors in the listed mucosal cell populations (top). Average spatial expression of chemokines across all mucosal cells versus mucosal position (left). J-S distance (color) between the chemokine expression profile and the spatial distribution of cell types expressing the cognate chemokine receptor. Gray indicates that the corresponding cell type does not express the specific chemokine receptor above background levels (STAR Methods). Included are only cells from the cecal mucosa of SPF mice.

(T) As in (S) but for the proximal colon of SPF mice.

(U) As in (S) but for the distal colon of SPF mice.

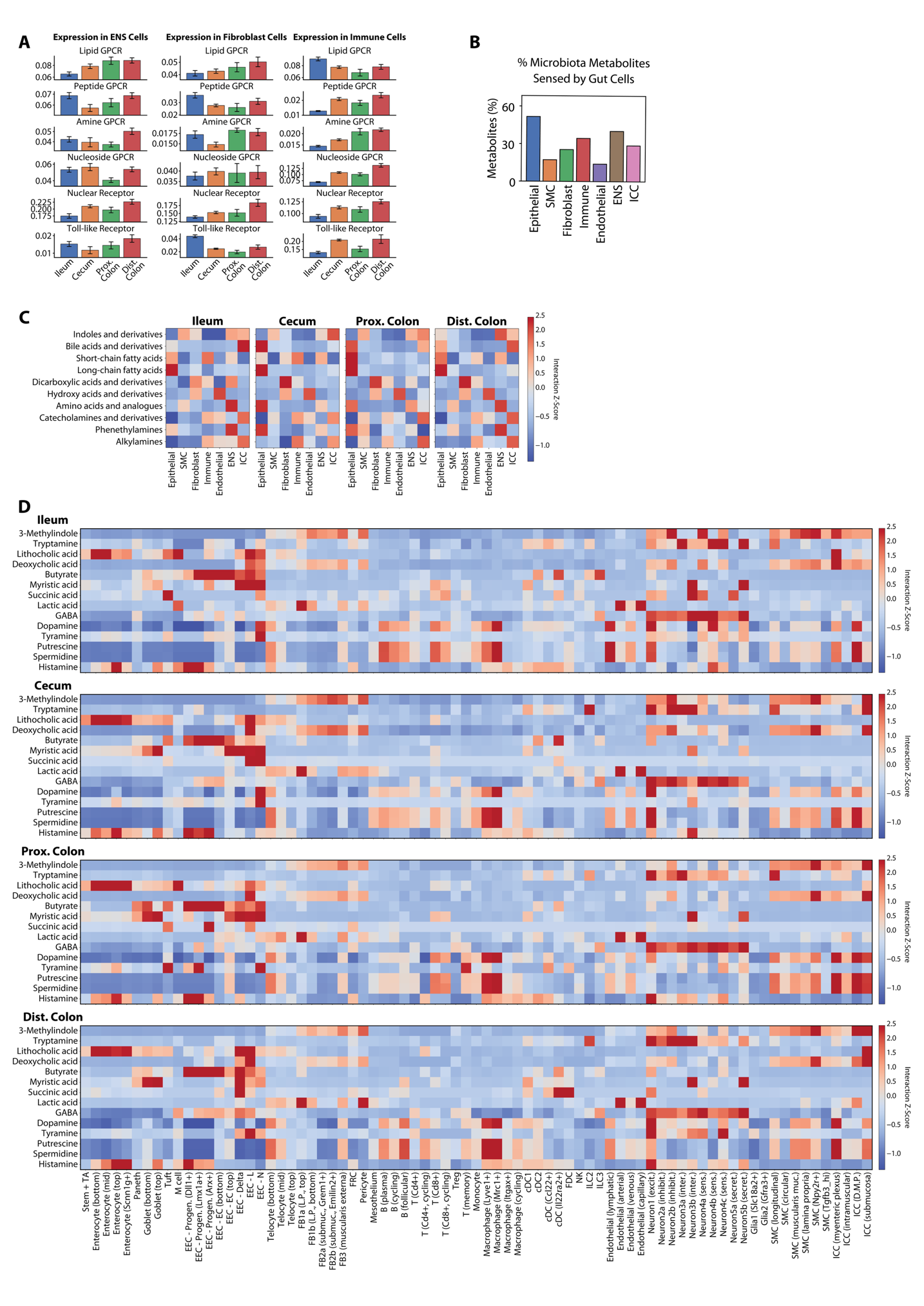

**Figure S6**. **Related to Figure 6. Regional specialization and conservation of receptor expression and responsivity to microbially derived small molecules.**

(A) Average expression of receptor categories within the listed major cell classes across regions in SPF mice. Bars and error bars represent the mean and standard error across slices (slice numbers in **Figure S2A**).
(B) Fraction of microbiota-derived metabolites listed in the MMRDB that interact with any cell type within each of the major cell classes.
(C) Predicted interaction strength (represented as z-scores across cell classes) between specific classes of microbiota-derived metabolites and major gut cell classes in each region of the SPF mouse.
(D) As in (C) but for the interaction between specific microbiota-derived metabolites and all gut cell types in the listed regions.

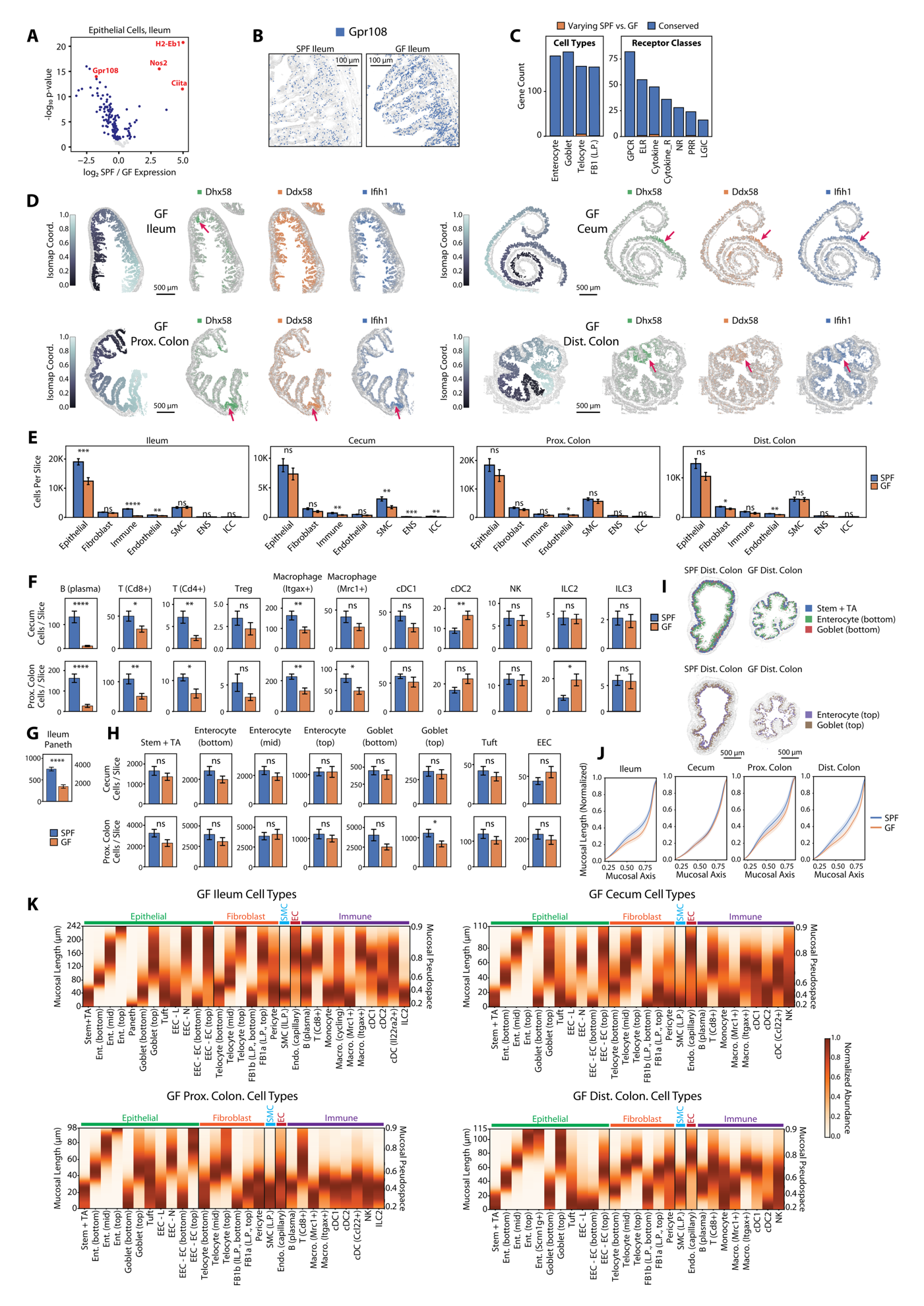

**Figure S7**. **Related to Figure 7. Selective tissue-scale, cellular, and transcriptomic remodeling of the mouse gut in the absence of the microbiome.**

(A) Logarithmic p-values for the differential expression of genes in all epithelial cells between SPF and GF conditions versus the log_2_-fold change in expression. Notable genes are marked in red. P-values were generated from a t-test and are FDR corrected (STAR Methods).

(B) Spatial distribution of RNAs of *Gpr108* (color) on top of all RNAs (gray) in a representative ileal slice of the SPF or GF mouse. Scale bar: 100 µm.

(C) The number of genes that are conserved (blue) or varying (orange) in their mucosal spatial distributions between SPF and GF states of the same region, among all genes expressed in both microbiome states in at least one region within the listed cell populations or receptor categories (STAR Methods).

(D) Spatial distribution of mucosal cells colored by circumferential isomap coordinates (left; STAR Methods) on top of non-mucosal cells (gray) with the spatial distribution of RNAs (right) colored by the listed genes on top of all RNAs (gray) in an example slice from each gut region of a GF mouse. Arrows mark a patch of coordinated interferon-stimulated gene expression within epithelial cells. Scale bar: 500 µm.

(E) Average number of cells per slice in each major cell class for each gut region of the SPF or GF mice. Bars and error bars represent the mean and standard error across slices; *, **, ***, and **** indicate p-values less than 0.05, 10^-2^, 10^-3^, and 10^-4^, respectively, while ns indicates a p-value greater than 0.05 as determined by a t-test (slice numbers in **Figure S2A**).

(F) Average number of select immune cells per slice in cecum or proximal colon of the SPF or GF mice. Bars and error bars represent the mean and standard error across slices (slice numbers in **Figure S2A**).

(G) As in (F) but for Paneth cells in ileum.

(H) As in (F) but for select epithelial cells.

(I) Spatial abundance and distribution of select cell types in the top or bottom mucosa in a representative slice of the distal colon of a SPF or GF mouse.

(J) The relationship between mucosal pseudospace and actual mucosal length, normalized by the mucosal thickness of the specified region and microbiome state. Lines represent the average across all slices and shaded regions represent the 95% confidence interval (slice numbers in **Figure S2A**).

(K) Average distribution of all major mucosal cell populations along the length of the mucosa determined by mucosal pseudospace (left) or mucosal distance from the crypt (right), in each gut region of GF mice. Only mucosal populations with sufficient abundances are shown (STAR Methods).

**Table S1. Related to Figure 1. Oligonucleotide sequences related to MERFISH (provide as a separate xslx file).**

In the sheet titled ‘Targeted Genes’, the ‘Gene’ column lists the name of each targeted RNA and the ‘Barcode’ column lists the binary barcode assigned to it. Entries that start with ‘Blank’ represent barcodes not assigned to an RNA and serve as a measure of false positive rates. In the sheet titled ‘Encoding Probes’, the column ‘Name’ lists a name for the template molecule associated with each MERFISH encoding probe while the column ‘Sequence’ lists the sequence associated with that template. In the sheet titled ‘Readout Probes’, the column ‘Name’ contains the name for the readout probe, the column ‘Sequence’ contains the sequence, and the column ‘Associated Bit or Gene’ contains the barcode bit to which the readout is associated if the entry is numeric or the gene to which the readout is assigned if the entry is a gene name. These genes were measured with sequential non-combinatorial stains. Readouts were conjugated to either Alexa488, Cy5, or Alexa750, as indicated in their name and sequence, via a disulfide bond, as indicated by ‘-S-S-’.

**Table S2. Related to Figures 3 and 4. Spatial organization of cells and gene expression in the mucosa (provided as a separate xslx file).**

In the sheet titled “Cell Abundance Gradients”, the ‘cell_type’ column lists the name of the cell type for which the abundance gradient is calculated. The ‘region’ column lists the gut region for which those cells were characterized. The ‘microbiome’ column lists the microbiome state, either specific-pathogen-free (SPF) or germ-free (GF). The ‘total_count’ column lists the total number of the listed cell type in the listed region and microbiome state. Column 5 and onwards list the normalized abundance of the given cell in the given region and microbiome state at the mucosal real space position in the heading (STAR Methods). The real space position is represented as a fraction of the total mucosal length with 0 for crypt base and 1 for top mucosa, and the cell abundance vectors are normalized to a maximum of 1. In the sheet titled ‘Gene Expression Gradients’, the ‘gene’ column lists the gene for which the expression gradient is calculated. The ‘cell_type’ column lists the name of the cell type expressing this gene. The ‘region’ column lists the gut region for which those genes were characterized. The ‘microbiome’ column lists the microbiome state, either SPF or GF. The ‘mean_expression’ column lists the mean log-normalized expression of the given gene in the given cell type, region, and microbiome state. Column 6 and onwards list the normalized expression of the given gene in the given cell type, region, and microbiome state at the mucosal pseudospace position in the heading (STAR Methods). The pseudospace position is represented as a number from 0 to 1, with 1 representing top mucosa. The expression vectors are normalized to a maximum of 1.

**Table S3. Related to Figure 4. Latent mutual information (LMI) and normalized mutual information (NMI) between gene expression and mucosal pseudospace positions (provided as a separate xslx file).**

In the sheet titled ‘LMI’, the ‘cell_type’ column lists the name of the cell type for which the LMI score is calculated. The ‘region’ column lists the gut region for which those cells were characterized. The ‘microbiome’ column lists the microbiome state, either specific-pathogen-free (SPF) or germ-free (GF). The ‘total_cells’ column lists the total number of the listed cell type in the listed region and microbiome state. The ‘lmi_with_space’ column lists the LMI score between the transcriptome of the given cell type and mucosal pseudospace positions (STAR Methods). In the sheet titled ‘NMI’, the ‘gene’ column lists the gene for which the NMI score is calculated. The ‘cell_type’ column lists the name of the cell type expressing this gene. The ‘region’ column lists the gut region for which those genes were characterized. The ‘microbiome’ column lists the microbiome state, either SPF or GF. The ‘total_cells’ column lists the total number of the listed cell type in the listed region and microbiome state. The ‘nmi_with_space’ column lists the NMI score between the expression of the given gene and mucosal pseudospace positions (STAR Methods).

**Table S4. Related to Figure 4. Spatially informed receptor-ligand interactions for example mucosal cell populations (provided as a separate xslx file).**

The ‘region’ column lists the profiled gut region. The ‘microbiome’ column lists the microbiome state, either specific-pathogen-free (SPF) or germ-free (GF). The ‘spatial_neighborhood’ column lists the specific anatomical region under consideration. The ‘ligand_cell_type’ column lists the cell type expressing the ligand. The ‘receptor_cell_type’ column lists the cell type expressing the receptor. The ‘ligand’ column lists the ligand gene (translated to human gene name). The ‘receptor’ column lists the receptor gene (translated to human gene name). The ‘mean [CellPhoneDB]’ column lists the CellPhoneDB interaction score. The ‘p_value [CellPhoneDB]’ column lists the p-value for the interaction. The ‘FDR’ column indicates whether the p-value is below an FDR-corrected p-value threshold of 0.05.

**Table S5. Related to Figure 5. Spatially variable genes (SVGs) detected around the circumference of gut cross-sectional slices (provided as a separate xslx file).**

The ‘gene’ column lists the name of the SVG. The ‘cell_type’ column lists the name of the cell type in which this SVG is detected. The ‘layer’ column lists the gut anatomical layer in which this SVG is detected. The ‘region’ column lists the gut region in which this SVG is detected. The ‘microbiome’ column lists the microbiome state, either specific-pathogen-free (SPF) or germ-free (GF). The ‘slice_full_name’ column lists the name of the slices in which this SVG is detected. The ‘gene_description’ column lists a description of this gene if it is a receptor or ‘nan’ if not. The ‘receptor_type’ columns lists the receptor category for receptors and ‘nan’ for other genes.

**Table S6. Related to Figure 6. Differentially expressed genes (DEGs) across gut regions or between microbiome states (provided as a separate xslx file).**

In the sheet titled ‘Regional Upregulation’, the ‘region’ column lists the specific-pathogen-free (SPF) gut region that is compared against the rest for the purpose of DEG analysis. The ‘cell_type’ column lists the name of the cell type expressing the given DEG. The ‘gene_name’ column lists the name of the DEG. The ‘mean_region [log normalized]’ column lists the log-normalized mean regional expression of the given gene in the given cell type. The ‘mean_rest [log normalized]’ lists the log-normalized mean expression of the given gene in the given cell type in all other regions. The ‘mean_region [linear normalized]’ column lists the normalized (without log-transform) mean regional expression of the given gene in the given cell type. The ‘mean_rest [linear normalized]’ column lists the normalized (without log-transform) mean expression of the given gene in the given cell type in all other regions. The ‘ratio [log2]’ column lists the log2-transformed ratio between the mean linear expression in the given region versus all other regions. The FDR-corrected ‘p-value’ columns lists the p-value for the differential expression. The ‘FDR’ column indicates whether the FDR-corrected p-value is below a threshold of 0.05. In the sheet titled ‘SPF vs. GF’, the ‘region’ column lists the gut region that is considered for the purpose of DEG analysis between SPF and germ-free (GF) conditions. The ‘cell_type’ column lists the name of the cell type expressing the given DEG. The ‘gene_name’ column lists the name of the DEG. The ‘mean_SPF [log normalized]’ column lists the log-normalized mean expression of the given gene in the given cell type and region in the SPF condition. The ‘mean_GF [log normalized]’ lists the log-normalized mean expression of the given gene in the given cell type and region in the GF condition. The ‘mean_SPF [linear normalized]’ column lists the normalized (without log-transform) mean expression of the given gene in the given cell type and region in the SPF condition. The ‘mean_GF [linear normalized]’ column lists the normalized (without log-transform) mean expression of the given gene in the given cell type and region in the GF condition. The ‘ratio [log2]’ column lists the log2-transformed ratio between the mean linear expression in the SPF versus GF condition. The ‘p-value’ columns lists the p-value of the statistical significance. The ‘FDR’ column indicates whether the p-value is below an FDR-corrected p-value threshold of 0.05.

**Table S7. Related to Figure 6. The Microbiota Metabolite - Receptor Database (MMRDB) and receptor – drug and receptor – metabolite interactions (provided as a separate xslx file).**

In the sheet titled ‘MMRDB’, the ‘Gene’ column lists the name of the receptor or downstream gene targeted by the microbiota metabolite. The ‘UniProt ID’ column lists the UniProt ID of the protein product of the gene. The ‘Metabolite’ column lists the common name of the metabolite proposed to interact with the gene. The ‘InChl’ column lists the International Chemical Identifier (InChI) of the metabolite. The ‘InChIKey’ column lists the InChIKey representation of the metabolite. The ‘PubChem CID’ column lists the PubChem chemical ID of the metabolite. The ‘CHEMBL’ column lists the EMBL chemical ID, if available. The ‘ChEBI’ column lists the Chemical Entities of Biological Interest ID of the metabolite, if available. The columns ‘Kingdom’, ‘Super class’, ‘Class’, ‘Sub Class’, ‘Direct Parent‘, and ‘Alternative Parents’ list the chemical classification of the metabolite according to the Human Metabolome Database (HMDB)^91^. The column ‘Source’ lists the database source from which this gene – metabolite interaction was derived. The column ‘Direct Interaction’ indicates whether the targeted gene is a direction receptor (TRUE) or downstream gene (FALSE). The column ‘Microbiota-Derived Metabolite’ indicates whether the metabolite is known to be produced by the human gut microbiome (TRUE) or not (FALSE). In the sheet titled ‘Microbiota Metabolite - Cell’, the column ‘cell_type’ lists the cell type. The column ‘metabolite’ lists the common name of the microbiota metabolite. The columns ‘SPF XX, interaction_score [drug2cell]’ lists the interaction score between the given metabolite and cell type where XX indicates the region of the SPF gut calculated by drug2cell, using MMRDB as the interaction database. The columns ‘GF XX, interaction_score [drug2cell]’ lists the interaction score between the given metabolite and cell type where XX indicates the region of the GF gut calculated by drug2cell, using MMRDB as the interaction database.
